# Temporal dynamics of QTL effects on vegetative growth in *Arabidopsis thaliana*

**DOI:** 10.1101/2020.06.11.145953

**Authors:** Rhonda C. Meyer, Kathleen Weigelt-Fischer, Dominic Knoch, Marc Heuermann, Yusheng Zhao, Thomas Altmann

**Author notes:** Corresponding author: Rhonda C. Meyer. Both authors contributed equally (Rhonda C. Meyer, Kathleen Weigelt-Fischer).

## Abstract

We assessed early vegetative growth in a population of 382 accessions of *Arabidopsis thaliana* using automated non-invasive high-throughput phenotyping. All accessions were imaged daily from seven to 18 days after sowing in three independent experiments and genotyped using the Affymetrix 250k SNP array. Projected leaf area (PLA) was derived from image analysis and used to calculate relative growth rates (RGR). In addition, initial seed size was determined. The generated data sets were used jointly for a genome-wide association study that identified 238 marker-trait associations (MTAs) individually explaining up to 8 % of the total phenotypic variation. Co-localisation of MTAs occurred at 33 genomic positions. At 21 of these positions, sequential co-localisation of MTAs for two to nine consecutive days was observed. The detected MTAs for PLA and RGR could be grouped according to their temporal expression patterns, emphasising that temporal variation of MTA action can be observed even during the vegetative growth phase, a period of continuous formation and enlargement of seemingly similar rosette leaves. This indicates that causal genes may be differentially expressed in successive periods. Analyses of the temporal dynamics of biological processes are needed to gain important insight into the molecular mechanisms of growth-controlling processes in plants.

**Highlight:** A genome-wide association study including the factor time highlighted that early plant growth in Arabidopsis is governed by several medium and many small effect loci, most of which act only during short phases of two to nine days.

## INTRODUCTION

Plant growth is a complex process integrating many genetic, metabolic and environmental factors at the level of cells, tissues, organs or whole plants. Growth in the model plant system *Arabidopsis thaliana* occurs in a sequence of distinct yet partially overlapping phases (Boyes *et al*., 2001), germination, seedling establishment, vegetative growth with successive appearance of leaves and progressive development of the root system, floral transition, flowering, seed production, senescence, each of which may be initiated and controlled by a network of different processes and responses to environmental cues (Beemster *et al*., 2005; Dubois *et al*., 2017; Schippers, 2015; Silva *et al*., 2016; Tisné *et al*., 2008; Weng *et al*., 2016). In this context, quantitative trait locus (QTL) mapping and genome-wide association analyses have often been applied to identify QTL/alleles for biomass and other growth-related traits. Examples include QTL for leaf area, growth rates and dry weight (El-Lithy *et al*., 2004; Lisec *et al*., 2008), for seed germination, seed longevity or seed dormancy (Clerkx *et al*., 2004; Nguyen *et al*., 2012), and for complex traits such as leaf shape (Juenger *et al*., 2005), or epistatic QTL for shoot and root growth (Bouteillé *et al*., 2012). In several cases, the genes underlying the QTL could be identified (Bentsink *et al*., 2006; Coluccio *et al*., 2010; Loudet *et al*., 2005; Riewe *et al*., 2016; Todesco *et al*., 2010). However, growth analyses were often restricted to one or a few time points during the development and consequently detected mostly cumulative effects (Zhu, 1995). The establishment of automated non-invasive high-throughput phenotyping systems (Furbank and Tester, 2011) allowed in-depth studies of many aspects of plant growth in model and crop plants, including Arabidopsis (Dornbusch *et al*., 2012; Granier *et al*., 2006; Lyu *et al*., 2017; Tisné *et al*., 2013), maize (Cabrera□Bosquet *et al*., 2016; Junker *et al*., 2015; Zhang *et al*., 2017), rice (Al-Tamimi *et al*., 2016; Campbell *et al*., 2015; Schilling *et al*., 2015), barley (Honsdorf *et al*., 2014; Neumann *et al*., 2017; Wang *et al*., 2019a), pea (Humplík *et al*., 2015), lentil (Muscolo *et al*., 2015) and rapeseed (Fanourakis *et al*., 2014; Kjaer and Ottosen, 2015; Pommerrenig *et al*., 2018). In particular, these automated platforms enabled almost continuous monitoring of plant growth and development at many time points during development. In Arabidopsis, a genome wide association study (GWAS) of projected leaf area at 12 different time points, parameters derived from growth models, and final biomass data revealed time-specific and general QTL affecting plant growth (Bac-Molenaar *et al*., 2015). Temporal patterns for growth and developmental traits have also been described for maize (Muraya *et al*., 2017), barley (Neumann *et al*., 2017), triticale (Liu *et al*., 2014), wheat (Ren *et al*., 2018), and rapeseed (Knoch *et al*., 2020; Wang *et al*., 2015). Taken together, these findings clearly show a need for time-resolved analyses of plant growth to detect loci showing temporal restricted expression patterns. We applied daily automated imaging to a population of 382 natural Arabidopsis accessions and performed genome-wide association analyses throughout early vegetative phases to address the following questions: (i) Can we resolve dynamic, time-restricted contributions of loci for early growth by a time course analysis? (ii) Does initial seed size affect vegetative growth (iii) Can we draw links to known QTL and loci? (iv) Are we able to identify candidate genes underlying the observed marker-trait-associations?

## MATERIALS AND METHODS

### Plant materials and growth conditions

The 382 Arabidopsis accessions (Table S1) were amplified together, and the number of siliques restricted to six per plant. Seeds from this amplification were sown in a controlled environment growth-chamber. After two days of stratification at 5°C in constant darkness, seeds were germinated and seedlings acclimated under a 16/8 h day/night regime with 16/14°C, 75% relative humidity, and 140 ± 10 μmol m^-2^ s^-1^ light intensity for three days. Parameters were then adjusted to 20/18°C, 60/75% relative humidity and 140 ± 10 µmol m^-2^ s^-1^ photosynthetically active radiation (PAR) from Whitelux Plus metal halide lamps (Venture Lighting Europe Ltd., Rickmansworth, Hertfordshire, England) still under a 16/8 h day/night regime. 12-well trays with a well size of 38×38×78 mm, cut from QuickPot QP 96T trays (HerkuPlast, Ering, Germany), were filled with a mixture of 85% (v) red substrate 2 (Klasmann-Deilmann GmbH, Geeste, Germany) and 15% (v) sand. Plants were watered with 45 ml water at 7 and 9 days after sowing (DAS), and then every other day until 19 DAS with 55 ml water, to maintain approximately 70% field capacity.

Plants were grown in three independent experiments over one year, arranged in a randomised complete block design with three replicates per experiment. Each replicate consisted of four individual plants grown in the same 12-well tray.

### Genotyping of accessions with 250K SNP chip

As no public 250k SNP data (Horton *et al*., 2012) were available for 64 Arabidopsis accessions, DNA of the missing accessions was hybridised to the Affymetrix 250K SNP Array (DNAVision, Charleroi, Belgium), and raw data subjected to the analysis pipeline established by Nordborg and colleagues (Atwell *et al*., 2010). Distribution of SNPs across the genome was visualized using the SNP-density plot function of the R package ‘rMVP’, a Memory-efficient, Visualization-enhanced, and Parallel-accelerated tool for genome-wide association studies, with bin size set to 10,000 bp. The package is available at github: https://github.com/XiaoleiLiuBio/rMVP.

The Araport database (Hanlon *et al*., 2015; Rosen *et al*., 2014; www.araport.org) and Polymorph1001 (1001GenomesConsortium, 2016; http://tools.1001genomes.org/polymorph) were used to classify SNPs in candidate genes.

### Population structure

Population structure was analysed using the software package STRUCTURE, version 2.3.4 (Pritchard *et al*., 2000). Population clustering for K= 1 to 10 using the ‘admixture’ model was performed with a burn-in period of 50,000, 50,000 MCMC replications and five iterations per K. Two approaches were combined to determine the best value for K, L(K) as described by Rosenberg *et al*. (2001), and ΔK introduced by Evanno *et al*. (2005).

### High-throughput non-invasive phenotyping

We assessed vegetative growth of the 382 Arabidopsis accessions at 12 different time points during development using the IPK automated phenotyping facility for small plants (Junker *et al*., 2015; https://www.ipk-gatersleben.de/en/phenotyping).

Plants were imaged daily between 7 and 18 days after sowing (DAS), and dry weight was determined at 20 DAS. The germination time was defined as the time of emergence of the cotyledons, and determined by manually scanning the top view fluorescent images taken from three days after sowing onwards. Projected leaf area (PLA) measurements were extracted from top view images in the visible light range using IAP (Klukas *et al*., 2014) and used to calculate relative growth rates (RGRs) as in Eq.1 in overlapping three-day intervals.

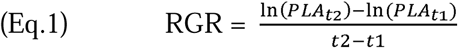

To measure seed size traits, 20 seeds per accession were fixed on an A4 sheet including a size standard and scanned with an Epson Expression 10000XL flatbed scanner (Seiko Epson Corporation, Suwa, Japan) at a resolution of 1200 dpi. Seed width, length and area were extracted using the custom program “Evaluator” (Meyer *et al*., 2012). The Evaluator algorithm isolates the seed image from the background based on differences in pixel intensities, creates a contour boundary and counts the pixels inside the boundary as a measure of area. Length and width of each seed are determined based on the seed’s orientation.

## Statistical analyses

Adjusted phenotypic means were extracted as best linear unbiased estimates (BLUEs) using the GenSTAT 17^th^ Edition (VSNi, Hempstead, UK) procedure REML and the following mixed linear model:

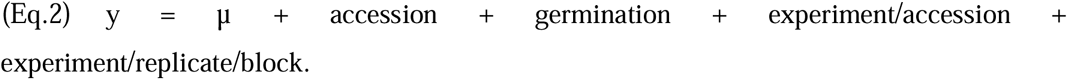

Plant genotypes (accession) were considered as fixed factor effects with days to emergence of cotyledons (germination proxy) as covariate. Combinatorial interactions between each set of experiments, replicates within the experiments and blocks (8 carriers moving together in the phenotyping facility) within the replicates were considered as random factor effects. Broad-sense heritability was calculated using the same mixed linear model, but with genotype as random factor:

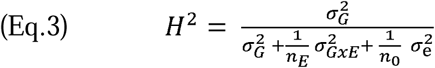

where *σ*_g_ is the genetic variance (accessions), *σ*_GxE_ is the variance of the experiment/accession interaction, *σ*_e_ is the error variance, n_E_ is the average number of experiments per accessions (n_E_=3), and *n*_*0*_ is the number of individual plants for each accession (n_0_=36; He *et al*., 2016). The following statistical analyses were performed in R version 3.4.4 software environment for statistical computing and graphics (RCoreTeam, 2018), and RStudio Version 1.1.383. Pearson correlations and associated *p*-values were estimated using the function ‘rcorr’ from the R package ‘Hmisc’ (V 4.1.1, https://cran.r-project.org/web/packages/Hmisc/index.html).

GWAS was performed with 26 traits and 212,142 SNP markers using FarmCPU (Liu *et al*., 2016). Principal components (PCs) to adjust for population structure were extracted from the GAPIT output (Lipka *et al*., 2012; Tang *et al*., 2016). For each trait, QQ plots were inspected to choose an appropriate number of PCs within the limits set by the STRUCTURE analysis (2-4 populations). The maxLoop parameter was increased to 30 and the optimal threshold for *p*-value selection of the model in the first iteration was estimated by the FarmCPU.P.Threshold function with 1,000 permutations and set to 0.000085 for all traits. Subsequently, *p*-values of marker-trait-associations were adjusted for multiple comparisons using FDR (Benjamini and Hochberg, 1995). Only associations with adjusted *p*-values below the FDR threshold of 0.05 were included in further analyses. The phenotypic variance explained (PVE%) by a significant marker was estimated in R as described in Knoch *et al*. (2020).

Linkage disequilibrium (LD) was individually measured for each chromosome as *r*^*2*^, the square of the allelic correlation coefficient of the pairwise physical distance between the 109,178 homozygous SNP markers with the R package *LDheatmap* (Shin *et al*., 2006). A modified equation (Marroni *et al*., 2011; Remington *et al*., 2001) based on expectations for *r*^*2*^ (Hill and Weir, 1988) was used to estimate the decay of *r*^*2*^ with distance implemented in R:

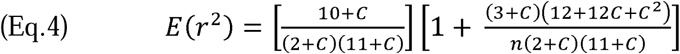

where n is the effective population size (764 gametes of 382 individuals) and c is the recombination fraction between sites and C = 4nc. The arbitrary C is estimated fitting a nonlinear model using the nls function in R and starting with C = 0.1. The estimated C is than refitted into the equation to model adjusted LD values aligned for their Euclidian distance along the chromosome. The intercept of the half maximum adjusted LD with the Euclidian pairwise distance between SNPs was the half LD decay value of the population. To estimate the degree of random co-localisation, permutation analyses were performed, distributing the detected associations randomly to all markers and extracting the number of co-localisations. This procedure was repeated 100,000 times.

## RESULTS

### Description of the mapping population

The 382 accessions were selected to represent a wide geographic distribution (Fig. S1), with a focus on accessions for which public 250K SNP data were available (Atwell *et al*., 2010). Hundred and one of these accessions were previously analysed in a nitrogen use efficiency study (Kuhlmann *et al*., 2020). For 64 accessions no public SNP data were available at the time. DNA of the missing accessions was extracted and hybridised to the Affymetrix 250K SNP Array. The raw data were subjected to the analysis pipeline established by Nordborg and colleagues (Atwell *et al*., 2010). The total number of 214,052 identified SNPs (call method 75; (Horton *et al*., 2012) was reduced to 212,142 SNPs by filtering for a minor allele frequency above 2% and missing values below 5% for use in GWAS. The SNP density plot (Fig. 1) reveals an even distribution of the markers across the genome. Overall population structure was low, with the first ten principal components (PCs) yielding a cumulative R^2^ of only 16.16% (3.67, 2.31, 2.02, 1.74, 1.33, 1.28, 1.07, 0.97, 0.95, 0.83 %, respectively). The mean Ln probability (L(K)) and the mean difference between successive likelihood values of K (ΔK) derived from the STRUCTURE output indicated an optimum K, i.e. number of subpopulations, between 2 and 4 (Fig. S2). The genome-wide half maximum LD decay was found to occur at a pairwise physical SNP distance of 3.37 kb. LD decay was also determined for each chromosome separately and amounted to 2.91 kb, 4.26 kb, 3.13 kb, 3.11 kb, and 3.82 kb for chromosomes 1 to 5, respectively.

**Figure 1:**
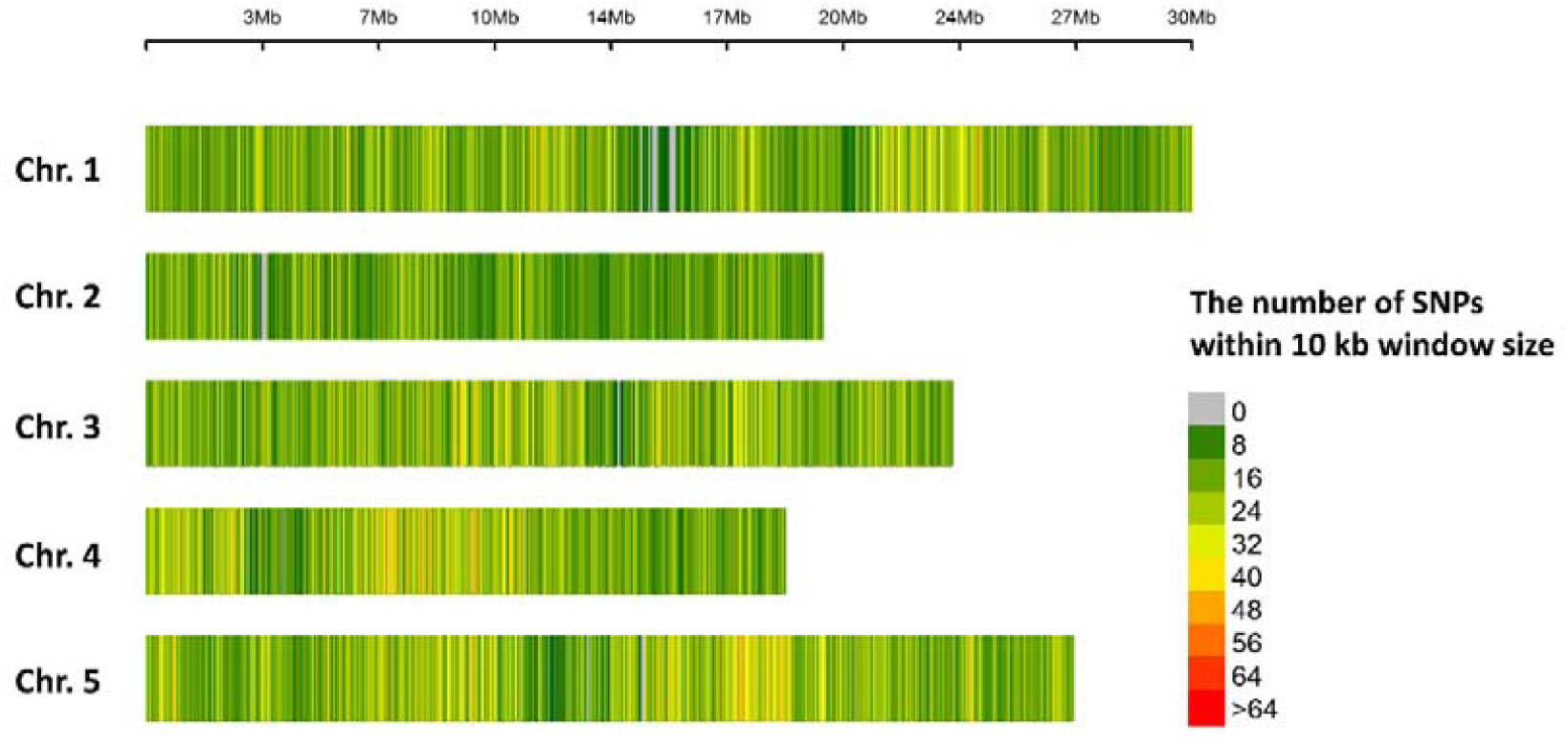
SNP density plot illustrating even SNP distribution across the genome. Shown is the genome-wide SNP marker distribution across the five Arabidopsis chromosomes. 212,142 unique, single-copy SNPs were binned in 10 kb intervals. The marker density is indicated by the colour legend (green to red) on the right side. Grey colour indicates regions without SNPs.

### Analysis of traits

Between 7 and 18 days after sowing (DAS), plants were phenotyped on a daily basis using top view visible light images. Best linear unbiased estimates (BLUEs) of projected leaf area (PLA) and relative growth rates (RGR) were obtained using a mixed linear model (Eq. 2). Three models were evaluated, all of which contained accession (genotype) as fixed factor: model 1 incorporated the day of emergence of the cotyledons (proxy for germination) as a covariate to account for different germination time points (3-7 DAS); model 2 included the seed size as covariate in the fixed model, and model 3 included both germination and seed size. Only germination showed a significant effect, and therefore model 1 with accession as fixed factor and germination time as covariate was used to obtain adjusted mean values. The same model with accession as random factor was used to estimate broad-sense heritabilities of PLA and RGR (Table S2). Heritabilities were moderate to high, ranging from 66% for RGR15_17 to 93% for seed area.

Pearson correlations were calculated between all phenotypic traits. The corresponding heatmap is presented in Fig. 2. The traits are clearly separated into three groups, corresponding to biomass (PLA and DW20), seed traits, and growth rates (RGR), respectively. Seed traits are positively correlated with PLA, and negatively with RGR, both at a low level. Correlations between PLA over time are all positive and highly significant. In contrast, correlation between RGRs over time are generally lower and switch from positive to negative during the late phase starting 14 DAS. This switch is even more pronounced in correlations between PLA and RGR.

**Figure 2:**
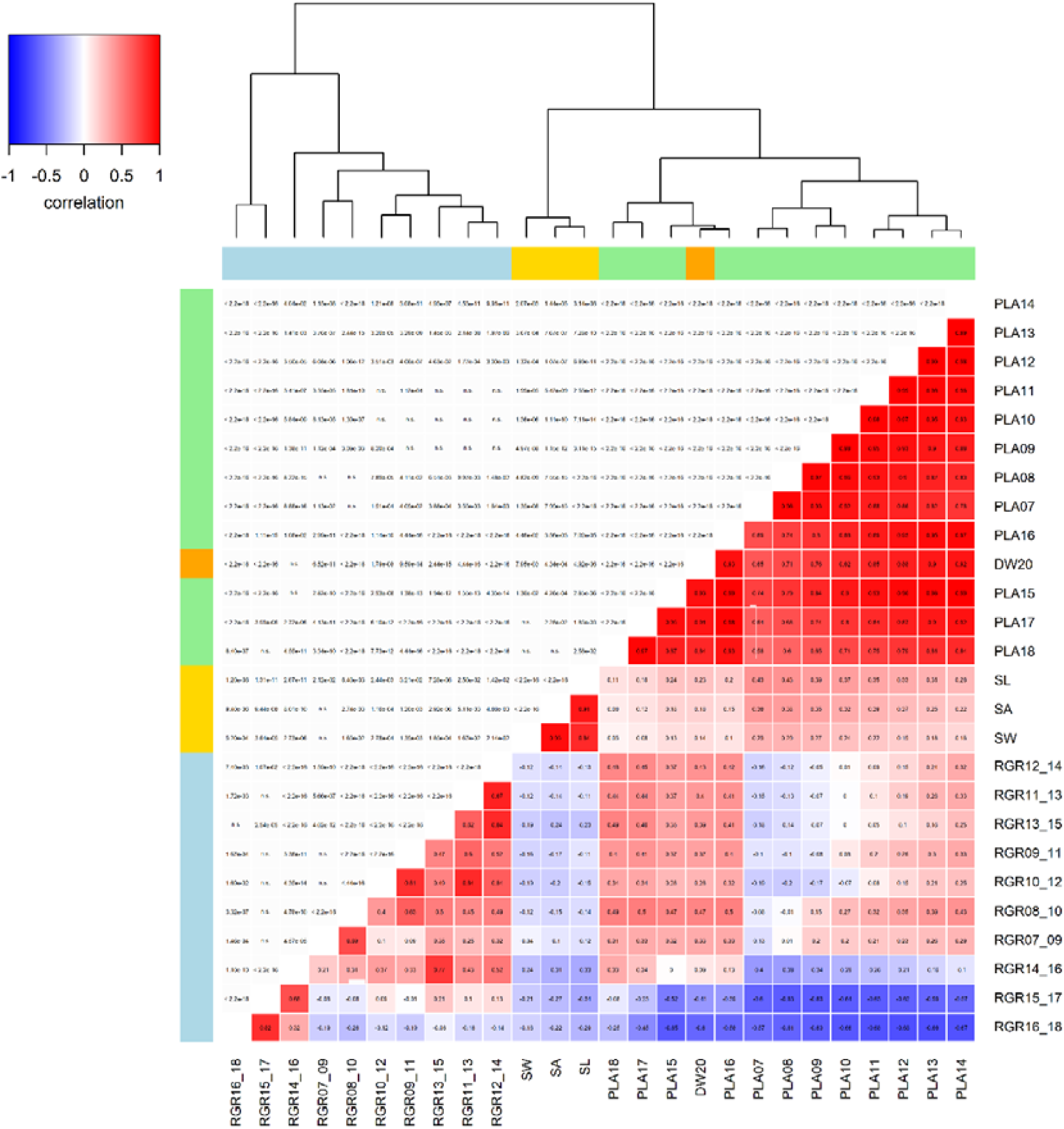
Correlation heatmap and hierarchical clustering of phenotypic means. Correlation heatmap and hierarchical cluster analysis of adjusted means of endpoint biomass (DW20), projected leaf area (PLA) and relative growth rates (RGR) over time, as well as seed area (SA), seed length (SL) and seed width (SW). The lower triangle displays the coefficients (r), the upper triangle the statistical significance (*p*-values). Colour scale for r: red, high correlation; blue, low correlation. Hierarchical clustering: colour differences in sidebars indicate the different trait groups within the clusters.

### Genome-wide association study

Genome-wide association studies (GWAS) were performed using the 26 phenotypic traits (dry biomass at 20DAS, PLA at 12 time points, RGR at 10 intervals, 3 seed traits) and 212,142 SNPs in FarmCPU (Liu *et al*., 2016). Correction for population structure was obtained by inclusion of the FarmCPU kinship matrix (Liu *et al*., 2016), and in addition inclusion of principal components (PCs); the optimal number of PCs for each trait (Table S3) was selected based on QQ plots (Fig. S3). Overall, 238 significant (*p*-value_(FDR)_ ≤ 0.05) marker-trait associations (MTAs) were discovered, explaining between 0.1% and 8.1% of the estimated phenotypic variance (Table S3). Final biomass (dry weight at 20 DAS, DW20) resulted in 10 MTAs, while the time-resolved projected leaf area (PLA) yielded 111 MTAs, and the relative growth rates (RGR) 85 MTAs; 32 MTAs were found for seed traits (seed area SA, seed length SL, seed width SW). The next step consisted in a search for co-localisations; two MTAs were considered co-localised if they were positioned within the chromosome-specific LD decay threshold from each other. MTAs of different traits co-localised at 33 positions (Table S3). In a permutation analysis with 100,000 repeats a maximum of 4 co-localisations per iteration was detected, consistent with the low number of detected associations (n=238) in relation to the number of markers (n=212,142).

MTAs for final biomass and leaf areas over time only shared three positions, no common MTAs were detected for final biomass and RGRs. Surprisingly, one co-localisation was found between seed area and RGR10_12. To explore similarities between our results and QTL reported in the literature, with physical distances available (Bac-Molenaar *et al*., 2015; El-Lithy *et al*., 2004; Knoch *et al*., 2017; Lisec *et al*., 2008; Meyer *et al*., 2010), we searched for co-localisations within a 10 kb interval around the SNP marker. The larger interval was chosen to harmonise our search with previous studies (Bac-Molenaar *et al*., 2015; Kim *et al*., 2007). MTAs co-localised (Table 1) at one position with a QTL for PLA extracted from Bac-Molenaar *et al*. (2015), at another position with a QTL for PLA identified by Meyer *et al*. (2010), and at three positions with metabolic QTL identified by Knoch *et al*. (2017) and Lisec *et al*. (2008). Two overlaps with known flowering genes were found (Table 1), one of which also coincides with the co-localised growth QTL of Bac-Molenaar *et al*. (2015).

**Table 1:**
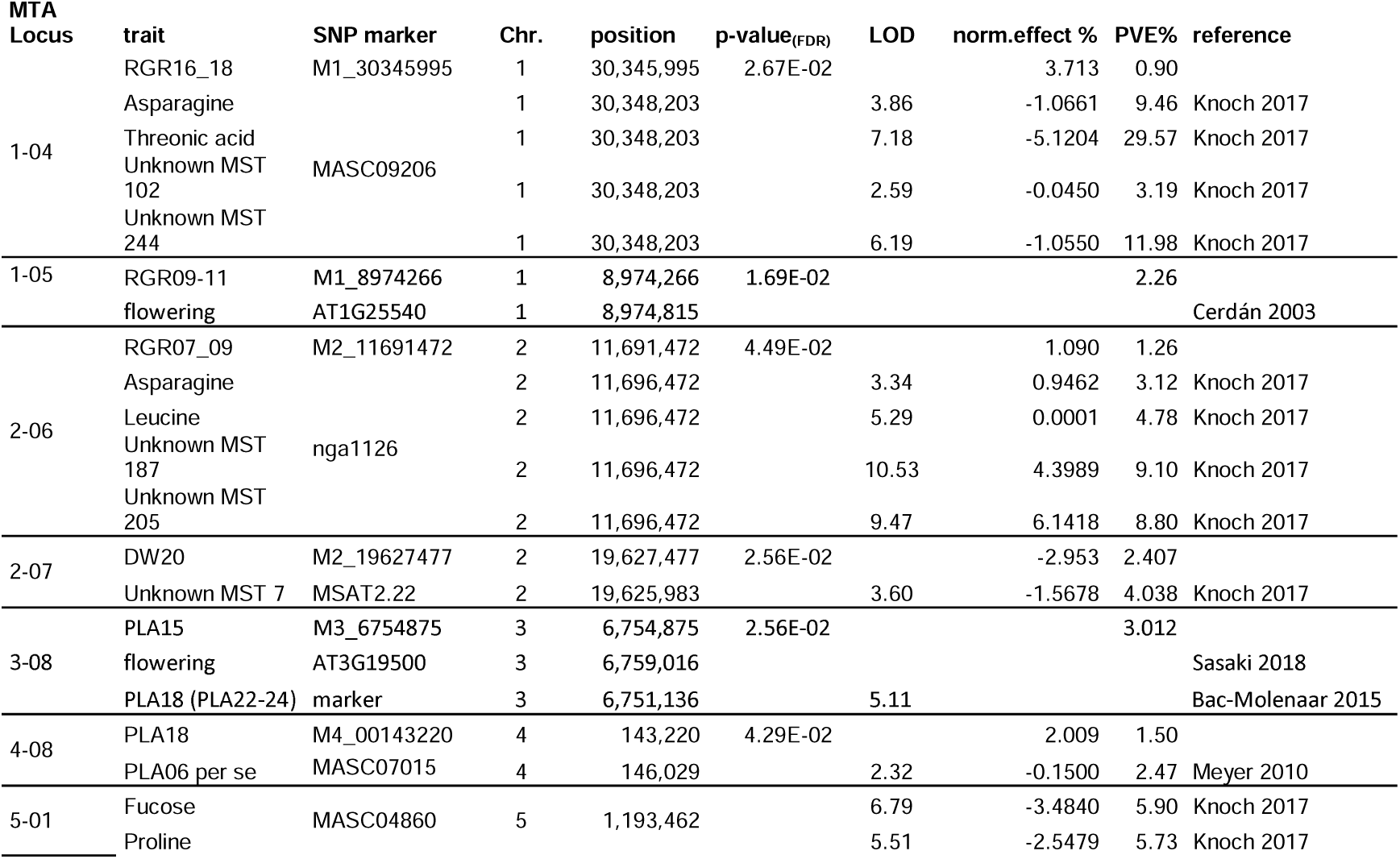
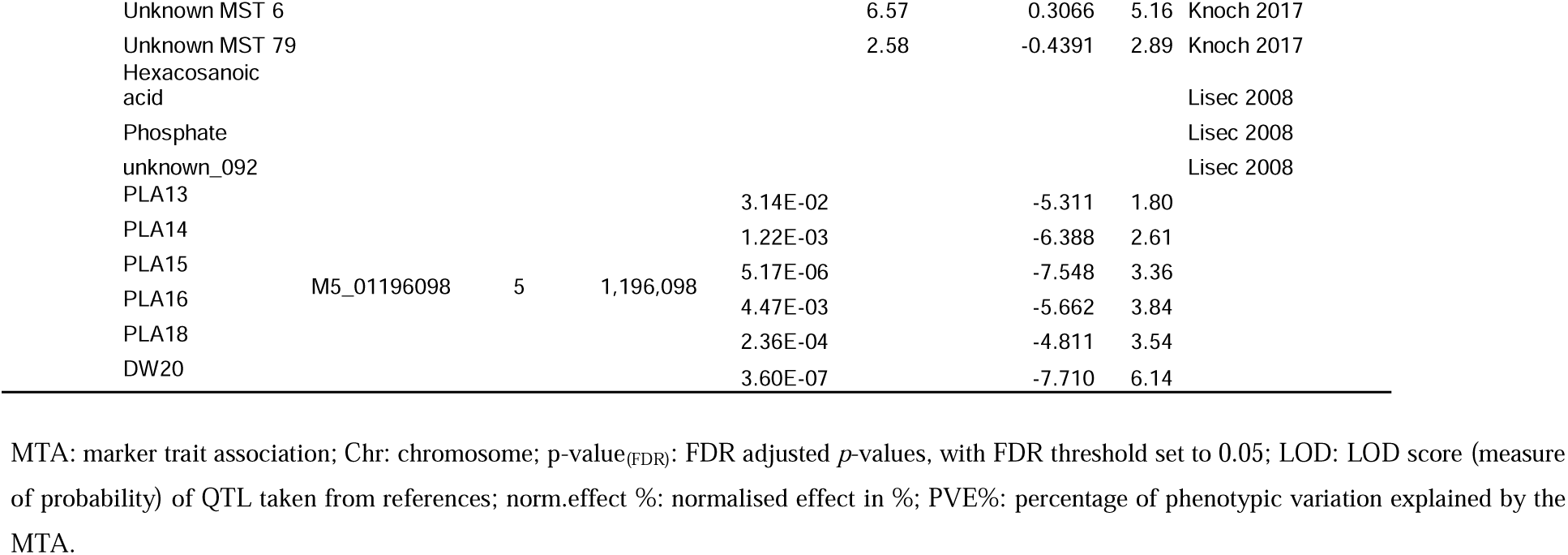
Co-localisation of detected MTAs with known growth, metabolite and flowering time QTL

We were particularly interested in the occurrence of growth MTAs over time, and investigated robust MTAs that were significant on at least two consecutive days. Overall, MTAs at 21 positions fulfilled this criterion, 15 PLA (Fig. 3A) and 6 RGR (Fig. 4A) loci. At two dynamic PLA loci, a MTA for RGR15-17 also co-localised, while a MTA for PLA12 co-localised at one dynamic RGR locus, but with reverse effect (Fig. 3B). A reversal of effects over time occurred in dynamic MTAs for RGR (Fig. 4B) only. Interestingly, one dynamic MTA for PLA coincided with a QTL previously described for metabolites in leaves (Lisec *et al*., 2008) and seeds (Knoch *et al*., 2017; Fig. S4).

**Figure 3:**
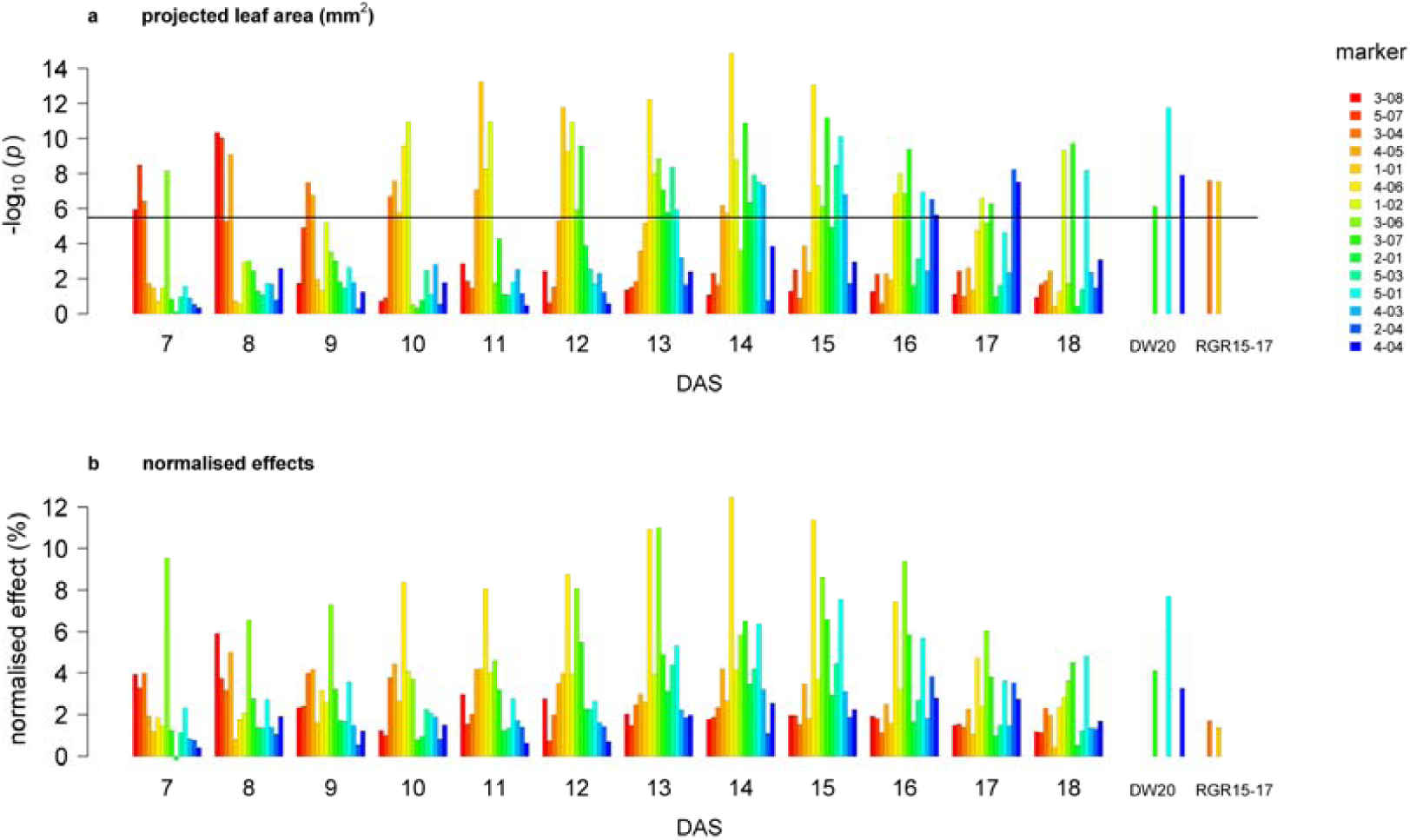
Dynamic MTAs for projected leaf area. Probabilities of 15 dynamic marker trait associations (MTA) (a) for projected leaf area over time (days after sowing, DAS), with normalised effects (b). The sign of the allelic effect is determined by the alphabetic order of the respective nucleotides of the SNP marker: a positive sign refers to the second allele in alphabetic order. The horizontal line indicates the significance threshold corresponding to *p*-value_(FDR)_ < 0.05. Also shown are co-localised MTAs for endpoint biomass (DW20) and relative growth rate between 15-17 DAS (RGR15-17). Different MTAs are represented by different colours, see legend. (Colours across figures 3 and 4 are not comparable)

**Figure 4:**
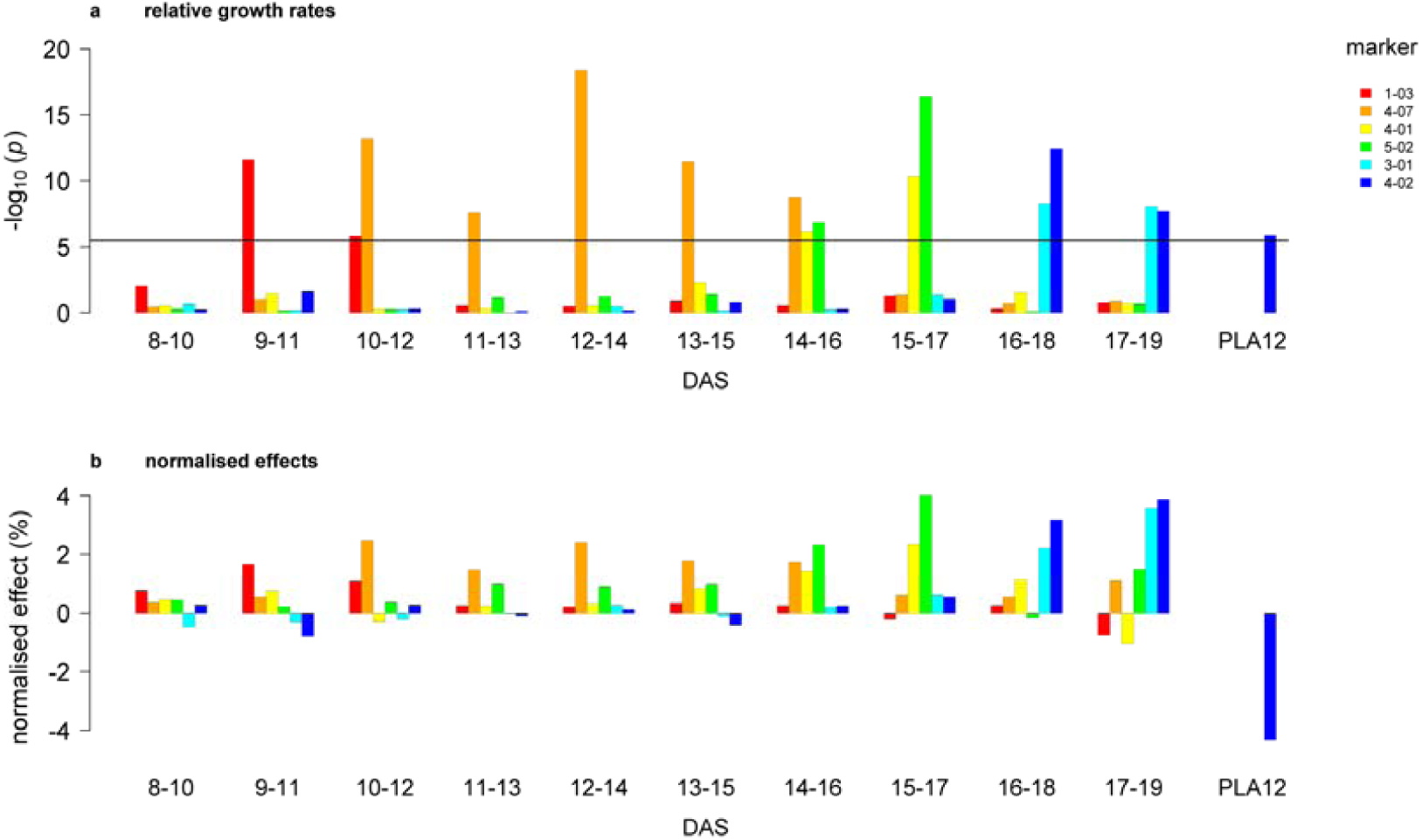
Dynamic QTL for relative growth rates. Probabilities for six dynamic marker trait associations (MTA) (a) for relative growth rates over time (days after sowing, DAS), with normalised effects (b). The sign of the allelic effect is determined by the alphabetic order of the respective nucleotides of the SNP marker: a positive sign refers to the second allele in alphabetic order. The horizontal line indicates the significance threshold corresponding to *p*-value_(FDR)_< 0.05. Also shown is a co-localised MTA for projected leaf area at 12 DAS (PLA12). Different MTAs are represented by different colours, see legend. (Colours across figures 3 and 4 are not comparable)

The dynamic MTAs were only significant during restricted periods ranging from two to nine days (Fig. S4), none were significant over the whole time (12 days). For PLA, we found three early QTL, two early/intermediate QTL, two early/intermediate/late QTL, one intermediate QTL, four intermediate/late QTL, two late and one QTL with breaks between the early, intermediate and late phases (Fig. S4). For RGR we detected one early QTL, one early/intermediate QTL, two intermediate/late QTL and two late QTL (Fig. S4). In the next step we looked for genes situated within the respective chromosome LD decay interval of each significant marker. In total we found 78 genes in or immediately adjacent to the MTA region, encoding one miRNA, three t-RNAs, five long non-coding RNAs, ten transposable elements and 59 genes encoding (putative) proteins (Table S4). Four significant markers associated with protein-coding genes (AT1G07680, AT1G60750, AT2G30690, AT3G07020) directly caused non-synonymous changes.

### Candidate genes

The confidence intervals around sixteen MTAs contained a total of 30 genes annotated to be involved in growth, cell wall, signalling, or transcription regulation (Table S5), with up to five genes in an interval. Additional SNPs and small insertions/deletions (indels) were identified in the candidate genes using the 250K SNP array data (Horton *et al*., 2012), Araport JBROWSE (Krishnakumar *et al*., 2015) and Polymorph 1001 (1001GenomesConsortium, 2016), yielding 1133 SNPs with high or moderate impact in the coding, promoter or UTR regions of 22 of these candidate genes (Table S5). Further mining of available databases and literature for possible links to plant growth let to a reduced list of nine most promising candidate genes for seven dynamic MTAs (Table 2).

**Table 2:** List of candidate genes for growth-related trait variation

| MTA Locus | trait | stage | gene | name | symbol | annotation | expression in |
| --- | --- | --- | --- | --- | --- | --- | --- |
| 1-01 | PLA | 10-13 | AT1G07670 | endomembrane-type CA-ATPase 4 | ECA4 | regulation of ABA and BL levels | germinating seed, seedling root |
| 1-03 | RGR | 8-11 | AT1G60790 | trichome birefringence-like protein (DUF828) | TBL2 | cell wall organization or biogenesis | seedling, young leaves |
| 1-06 | SA<br>RGR | seed<br>10-12 | AT1G16060 | ARIA-interacting double AP2 domain protein | WRI3 | involved in regulating seedling growth. | flower, petiole mature leaf |
| 3-01 | RGR | 15-18 | AT3G07020 | UDP-Glycosyltransferase superfamily protein | UGT80A2 | lipid glycosylation; reduced seed size | root, leaves, stem, flower |
|  |  |  | AT3G07030 | ALBA DNA/RNA-binding protein | ALBA | nucleic acid binding | stem, root, mature leaves |
| 3-04 | RGR | 15-17 | AT3G23730 | xyloglucan endotrans-glucosylase/hydrolase 16 | XTH16 | cell wall modification | germinating seed, root apex, young leaves, flower |
| 3-07 | PLA | 12-20 | AT3G49380 | plant-specific IQD family | IQD15 | CaM-dependent Ca<sup>2+</sup> signalling | seedling root, root, maturing seed |
|  |  |  | AT3G49390 | CTC-interacting domain 10 | CID10 | RNA-binding protein RBP37 | seedling root, root, leaves, maturing seed |
| 4-03 | PLA | 14-15 | AT4G13620 | Integrase-type DNA-binding superfamily protein | ERF062 | transcription regulation | seedling root, root |

List of the nine most promising candidate genes indicating the relevant MTA locus, the associated trait (PLA: projected leaf area, RGR: relative growth rate, SA: seed area) and stage (as time period given in days after sowing). The expression is derived from the eFP Browser Klepikova Atlas (Klepikova *et al.*, 2016; Winter *et al.*, 2007).

## DISCUSSION

Unravelling the genetic basis of complex traits governing plant performance and discovering the underlying molecular mechanisms remains a major undertaking in plant biology. The main aim of this study was the identification of genetic factors that influence early vegetative growth in Arabidopsis over a period of 12 days (7 to 18 DAS). We applied daily automated high□throughput phenotyping in the IPK phenotyping platform for small plants (Junker et al., 2015) to a diverse collection of 382 *Arabidopsis thaliana* accessions, extracted data for projected leaf area (PLA) at 12 time points, and calculated relative growth rates based on PLA. Previous studies have demonstrated that in Arabidopsis, biomass is highly correlated with leaf area (Leister *et al*., 1999; Meyer *et al*., 2004), enabling us to use PLA as a proxy for biomass.

Hierarchical clustering of the phenotypic data revealed a separation between early/intermediate (7 – 14) and late (15-18) phases. One possible explanation may be the increasing overlap of leaves, and therefore underestimation of PLA, at the later stages. It may also reflect morphological differences between leaves appearing at different stages (heteroblasty; Berardini *et al*., 2001). Similarly, the pronounced switches in correlations between PLA and RGR may indicate distinct growth phases, in particular floral transition in the shoot apical meristem (SAM). The transition from vegetative to reproductive SAM is terminal in the annual Arabidopsis, occurs before any visible sign of flowering and slows down vegetative leaf growth (Cookson *et al*., 2007). In our long-day conditions, some accessions started bolting as early as 18 days after sowing. Another possibility is a link to the appearance of leaves, as speculated for rapeseed (Knoch *et al*., 2020).

For all analysed time points, a total of 236 associations with endpoint biomass, the 22 growth-related and the three seed traits were detected at *p*-value_(FDR)_ ≤ 0.05. Most of the detected MTAs explained only a small percentage of phenotypic variance (< 5 PVE%, Table S3). In total, only 9 (3.8 %) MTAs with larger effects (> 5 PVE%) were detected, similar to a study in rapeseed (Knoch *et al*., 2020), confirming that plant growth results from the cumulative effects of the interaction of numerous small effect genes. We found a surprisingly large number of associations for RGR, individually explaining up to 8% phenotypic variance, the highest value found in this study. Robust phenotypic values obtained by calculating RGR over rolling 3-day intervals certainly contributed to the successful GWAS.

Several MTAs co-localised with previously described QTL, with the caveat that physical marker positions are only available for a restricted number of studies. This is particularly true for flowering time, where only eight QTL were available for comparison (Alonso-Blanco *et al*., 1998; Clarke *et al*., 1995); therefore known flowering genes (Sasaki *et al*., 2018; Srikanth and Schmid, 2011; Wellmer and Riechmann, 2010) were also included. The flowering time gene *PFT1* (*AT1G25540*, (Cerdán and Chory, 2003) co-localising with MTA1-05 for RGR09-11 is also involved in the control of organ size (Xu and Li, 2011) and the transcriptional regulation of genes involved in cell elongation and cell wall composition (Seguela-Arnaud *et al*., 2015). The flowering time gene *AT3G19500* encodes a bHLH DNA-binding superfamily protein that is part of the genetic network underlying flowering time regulation; its expression is positively correlated with flowering time, and negatively correlated with the expression of FLC (Sasaki *et al*., 2018). This gene mapped to the same region as MTA3-08 for PLA15 from this study, and MTA 3/6.8 for PLA18 identified by Bac-Molenaar *et al*. (2015). According to their experimental set-up, plants were transferred from stratification 4, 5, or 6 days after sowing, and this day was counted as day 1, therefore their PLA18 corresponds to PLA at 22-24 days after sowing. Despite 244 common accessions, this is the only shared MTA between these two growth studies, most likely due to different growth conditions and measurement at different time points. A large influence of even slightly different growth conditions was demonstrated in a comparison of the growth of three Arabidopsis accessions across ten laboratories (Massonnet *et al*., 2010). Therefore, the influence on growth of candidate genes located in this MTA is potentially stable across various environmental conditions.

The co-localisation between MTAs for seed area and RGR10_12 was unexpected, as seed area displayed no significant effect in the mixed linear analysis. Seed size has been shown to influence early vegetative growth in Arabidopsis (Elwell *et al*., 2011; Meyer *et al*., 2004), but this effect can be neutralised by restricting the number of siliques per plant (Meyer *et al*., 2004). However, the shared MTA region contains *AtWRINKLED3 (AT1G16060)*, which encodes an AP2-domain protein that interacts with a positive regulator of the ABA response (ARIA) and is involved in regulating seedling growth (Lee *et al*., 2009). Conversely, ABA has been shown to be involved in endosperm development (Cheng *et al*., 2014), and seed size is at least partially determined by endosperm growth (Sun *et al*., 2010). The observed link may well reflect different actions of the same gene during different developmental phases. Similarly, the co-localisation of a QTL for seed proline content (Knoch *et al*., 2017) with MTAs for PLA between 13 and 18 DAS and dry weight at 20 DAS may be due to the influence of parental seed composition on growth and development of the next generation (Alonso-Blanco *et al*., 1999; Elwell *et al*., 2011). In early studies in wheat, seed proline content was positively correlated with seedling growth (Lowe *et al*., 1972). Proline has been associated with general stress tolerance (Ashraf and Foolad, 2007); it accumulates in maturing Arabidopsis seeds (Chiang and Dandekar, 1995) where it seems essential for embryo development (Funck *et al*., 2012) and stimulates Arabidopsis germination (Hare *et al*., 2003). Given these findings, it is conceivable that differences in seed proline content may translate into growth differences. The associated candidate gene *At5g04275* encodes miRNA172, which is has been implicated in early vegetative development in Arabidopsis (Martin *et al*., 2010) and in proline accumulation under drought stress in potato (Yang *et al*., 2013), and which shows higher abundance in fast growing Arabidopsis mutants overexpressing purple acid phosphatase 2 (Liang *et al*., 2014).

The daily imaging performed during the phenotyping experiments allowed the analysis of the temporal dynamics of detected growth QTL. To address robust associations, only MTAs significant at two consecutive time points were considered for a detailed analysis. A total of 21 of these associations were detected, 15 for projected leaf area and six for relative growth rate. The elucidation of growth dynamics by means of time-dependent QTL analysis has been addressed in several studies in model and crop plant species, including Arabidopsis (Bac-Molenaar *et al*., 2016; Bac-Molenaar *et al*., 2015; Marchadier *et al*., 2019; Meyer *et al*., 2010), Setaria (Feldman *et al*., 2017), rice (Al-Tamimi *et al*., 2016; Campbell *et al*., 2017; Wu *et al*., 2018), maize (Muraya *et al*., 2017; Wang *et al*., 2019b; Zhang *et al*., 2017), barley (Neumann *et al*., 2015; Pham *et al*., 2019), rye (Miedaner *et al*., 2018; Würschum *et al*., 2014) and rapeseed (Knoch *et al*., 2020; Wang *et al*., 2015), with phenotyping frequencies varying from daily to weekly. The high temporal resolution provided by the present study coupled to the advantages of the Arabidopsis model system (small plant size, small and annotated genome, plethora of publicly available genetic and genomic resources) and the fast LD decay in our population facilitate the identification of putative candidate genes. In concordance with a previous genome-wide association study in Arabidopsis (Bac-Molenaar *et al*., 2015), we detected only period-specific MTAs affecting growth, and none significant over the whole time. Only three MTAs for endpoint biomass (DW20) co-localised with sequential MTAs for PLA, all in the intermediate to late phase; none overlapped with MTAs for RGR. Similar observations were made during the analyses of plant growth dynamics in maize (Muraya *et al*., 2017) and rapeseed (Knoch *et al*., 2020). Determining only endpoint biomass for input in a GWAS therefore severely limits the number of growth controlling genetic factors that can be detected.

Of the 59 putative protein-encoding genes located within dynamic MTA regions for PLA and RGR, eleven were annotated as encoding hypothetical proteins and another eleven genes could not be assigned a function. The remaining 37 genes were screened in available databases (Araport, TAIR, eFP Browser) and literature for possible links to plant growth, reducing the list to 30 candidate genes. Of these candidates, nine genes displayed relevant expression patterns (leaves, roots, seedlings) and/or mutant growth behaviour; three genes contained both deleterious and missense SNPs, six genes harboured only moderate effect SNPs in the coding region, promoter or UTRs. Moderate effect SNPs may be of particular interests in attempts to identify alleles modulating the growth performance, without the possible pleiotropic effects caused by gene disruption. The low number of genes in the MTA regions should facilitate validation using time- and tissue-resolved expression analyses. *AtECA4* (AT1G07670) encoding an endomembrane-type CA-ATPase 4 is a possible candidate within MTA1.1 for PLA10-14. Nguyen *et al*. (2018) showed that *AtECA4* is involved in the recycling of endocytosed cargo proteins such as ABCG25 and BRI1 from the trans-Golgi network/early endosome to the plasma membrane. This process has been described to be crucial to regulate homeostasis of the cellular ABA levels, and brassinolide (BL)-mediated signalling for growth. Mutant *eca4* plants showed multiple phenotypes including enhanced ABA sensitivity, increased resistance to dehydration and NaCl stresses, and more robust vegetative growth of shoots and roots. Candidate gene AT1G60790 (*AtTBL2*) is located within MTA1-03 for RGR08-11, belongs to the ‘trichome birefringence like’ (TBL) gene family with 46 members in Arabidopsis and clusters in the same clade as *TBR, TBL1* and *TBL4* (Gao *et al*., 2017). *TBR* (AT5G06700) and *TBL29/ESK1* (AT3G55990) are involved in cell wall biogenesis and modification with mutants showing impaired growth (Bischoff *et al*., 2010; Lefebvre *et al*., 2011; Xiong *et al*., 2013). In rice, trichome birefringence-like (tbl) mutants affected in xylan *O*-acetylation displayed a stunted growth phenotype (Gao *et al*., 2017). The MTA3-01 region for RGR15-18 contained two possible candidate genes: AT3G07020 and AT3G07030. AT3G07020 encodes an UDP-glucose:sterol glucosyltransferase, 80 UGT80A2, that is required for steryl glycosides and acyl steryl glycosides, and mutant *ugt80A2* seedlings have been described to show reduced root growth, with overall minor effects on plant growth (DeBolt *et al*., 2009), and a lower seed mass (Stucky *et al*., 2014). The second gene, AT3G07030, encodes an ALBA DNA/RNA-binding protein potentially involved in transcription regulation (Goyal *et al*., 2016). In Arabidopsis, ALBA proteins have been associated with rhizoid and root hair growth, with mutants *alba1* and *alba2* displaying reduced elongation (Honkanen *et al*., 2016). *AtXTH16* (AT3G23730) is the most likely candidate for MTA3-06 (PLA12-16). XTH16 is a xyloglucan endotransglucosylase/hydrolase 16, potentially involved in cell wall modifications (Sasidharan *et al*., 2010). Recent studies revealed that the expression of *AtXTH16* is GA-induced and *PKL*-dependent and correlates with larger plants (Park *et al*., 2017). Among the three genes within the MTA3-07 region for PLA12-20, AT2G49380 (*iqd15*) belongs to one of the plant-specific IQD families that have been described as scaffold-like proteins containing the IQ67 calmodulin binding domain that may link CaM-dependent Ca2+ signalling to cell function, shape, and growth (Bürstenbinder *et al*., 2017). Another possible candidate could be the cytokinin responsive gene AT3G49390 (*CID10*); however, T-DNA insertion mutants did not show an altered growth phenotype (Bravo *et al*., 2005). AT4G13620, adjacent to the MTA4-03 region (PLA14-15), encodes the ethylene-responsive transcription factor ERF062 belonging to the DREB subgroup A6 within the ERF/AP2 transcription factor superfamily (Weber and Hellmann, 2009), and has been shown to be nitrate responsive (Menz *et al*., 2016).

The functional diversity of the candidate genes identified in this study is yet another reminder of the complexity of plant growth, which necessitates the coordinated action of a large number of genes active at different time points during development.

## CONCLUSIONS

In this study we analysed the early growth of a diversity population of Arabidopsis accessions using high-throughput phenotyping at a high temporal resolution, and detected both single timepoint (general) and multiple timepoint (dynamic) MTAs. The inspection of the genes located in the MTA regions delivered potential targets for in-depth time-resolved functional analyses. The scarcity of shared QTL between endpoint biomass and PLA (proxy for biomass) or RGR over time illustrates the need for analyses of the temporal dynamics of biological processes to gain important insight into the molecular mechanisms of growth-controlling processes in plants.

## Supporting information

List of Supplementary Data

## ACKNOWLEDGMENTS

We thank Alexandra Rech, Iris Fischer, Monika Gottowik, Beatrice Knüpfer, Manuela Kretschmann, Marion Michaelis, Ingo Mücke and Gunda Wehrstedt for excellent technical assistance.

This research was supported through institutional funds of the Leibniz Institute of Plant Genetics and Crop Plant Research (IPK) and did not receive any specific grant from funding agencies in the public, commercial, or not-for-profit sectors.

## SUPPLEMENTARY DATA

Table S1: Overview of accessions used in this study

Table S2: Adjusted means and broad sense heritability of phenotypic data

Table S3: Overview of the 238 detected MTAs

Table S4: Overview of the 79 genes in or adjacent to LD interval around significant marker Table S5: List of 30 potential candidate genes

Figure S1: Geographic origin of the 382 analysed Arabidopsis accessions

Figure S2: Assessment of population structure

Figure S3: QQ plots with inclusion of PCs for population structure correction

Figure S4: Duration of dynamic MTA and co-localisation with known QTL

