## Supplementary Table S1 for "Temporal dynamics of QTL effects on vegetative growth in *Arabidopsis thaliana*"

| name | stock nr | array ID call method 75 | array ID new | latitude | longitude | location |
|---|---|---|---|---|---|---|
| 11PNA4 | CS76084 | 1245 | | 42.0945 | -86.3253 | Bainbridge, MI |
| 328PNA | CS76085 | 422 | | 42.0945 | -86.3253 | Bainbridge, MI |
| Aa-0 | CS28007 | 216 | | 50.9167 | 9.57073 | Aua/Rhön |
| Ak-1 | | 1058 | | 48.0683 | 7.62551 | Achkarren/Freiburg |
| Akita | 252AV | | 1516 | 39.43 | 140.06 | Yurihonjo, Akita |
| Alc-0 | CS76088 | 239 | | 40.31 | -3.22 | Alcala de Henares/Madrid |
| ALL1-2 | CS76089 | 363 | | 45.2667 | 1.48333 | Allassac |
| Alst-1 | CS28013 | 217 | | 54.8 | -2.4333 | Alston Moor |
| Amel-1 | CS28014 | 470 | | 53.448 | 5.73 | Ameland-Firehouse |
| An1 | CS76091 | 55 | | 51.2167 | 4.4 | Antwerpen |
| An-2 | CS28017 | 349 | | 51.2167 | 4.4 | Antwerpen |
| Ang-0 | CS28018 | 471 | | 50.3 | 5.3 | Angleur |
| Ang-0 | N949 | 184 | | 50.3 | 5.3 | Angleur |
| Ang-1 | N951 | | 1526 | 50.3 | 5.3 | Angleur |
| Ann-1 | CS28049 | 374 | | 45.9 | 6.13028 | Annecy |
| Arby-1 | CS28051 | 472 | | 59.4308 | 16.7999 | Asby/Helgaröby |
| Ba-1 | CS28053 | 335 | | 56.5459 | -4.79821 | Blackmount |
| Ba1-2 | CS76093 | 240 | | 56.4 | 12.9 | Båstad |
| Baa-1 | CS28054 | 375 | | 51.3333 | 6.1 | Baarlo |
| Bay-0 | CS76094 | 134 | | 49 | 11 | Bayreuth |
| Bch-1 | | | 1548 | 49.5166 | 9.3166 | Buchen/Lauenburg |
| Bd-0 | N963 | 1062 | | 52.4584 | 13.287 | Bensheim/Bergstraße |
| Be-0 | N964 | | 1546 | 49.6803 | 8.6161 | Bensheim/Bergstraße |
| Be-1 | CS28063 | 301 | | 49.6803 | 8.6161 | Bensheim/Bergstraße |
| Belmonte494 | CS76095 | 487 | | 42.1167 | 12.4833 | Belmonte in Sabina |
| Benk-1 | CS28064 | 350 | | 52 | 5.675 | Bennekom |
| Bg-2 | CS76096 | 214 | | 47.6479 | -122.305 | Seattle, WA, University of Washington |
| Bl-1 | N968 | | 1559 | 44.5041 | 11.3396 | Bologna |
| Bla-1 | CS76097 | 242 | | 41.6833 | 2.8 | Blanes/Gerona |
| Bla-11 | N984 | 1066 | | 41.6833 | 2.8 | Blanes/Gerona |
| Blh-1 | CS76098 | 243 | | 48 | 19 | Bulhary |
| Blh-1 | N1030 | | 1557 | 48 | 19 | Bulhary |
| Blh-2 | CS28090 | 336 | | 48 | 19 | Bulhary |

| name | stock nr | array ID call method 75 | array ID new | latitude | longitude | location |
|---|---|---|---|---|---|---|
| Boot-1 | CS28091 | 376 | | 54.4 | -3.2667 | Boot, Eskdale |
| Bor-1 | CS76099 | 56 | | 49.4013 | 16.2326 | Borky/Brno, Moravia |
| Bor-4 | CS76100 | 135 | | 49.4013 | 16.2326 | Borky/Brno, Moravia |
| Br-0 | N994 | 136 | | 49.2 | 16.6166 | Brno |
| Bs-1 | N997 | 185 | | 47.5 | 7.5 | Basel |
| Bs-2 | CS28097 | 445 | | 47.5 | 7.5 | Basel |
| Bsch-0 | CS28099 | 218 | | 40.0167 | 8.6667 | Buchschlag/Frankfurt |
| Bsch-2 | N1004 | 1067 | | 40.0167 | 8.6667 | Buchschlag/Frankfurt |
| Bu-0 | CS76103 | 245 | | 50.5 | 9.5 | Burghaun/Rhön |
| Bu-2 | N1008 | | 1570 | 50.5 | 9.5 | Burghaun/Rhön |
| Bur-0 | CS76105 | 137 | | 54.1 | -6.2 | Burren |
| C24 | CS76106 | 138 | | 40.2077 | -8.42639 | Coimbra |
| Ca-0 | CS28128 | 220 | | 50.2981 | 8.26607 | Camberg/Taunus |
| Cal-0 | N1063 | | 1591 | 53.2699 | -1.64293 | Calver |
| CAM-16 | CS76107 | 1075 | | 48.2667 | -4.58333 | Camaret-sur-Mer |
| CAM-61 | CS76108 | 208 | | 48.2667 | -4.58333 | Camaret-sur-Mer |
| Can-0 | N1064 | | 1568 | 29.2144 | -13.4811 | Lanzarote, Mirador de Rio |
| Cen-0 | CS76110 | 247 | | 49 | 0.5 | Caen |
| Cha-0 | CS28133 | 473 | | 46.0333 | 7.1167 | Champex |
| Chat-1 | CS28135 | 397 | | 48.0717 | 1.33867 | Châteaudun |
| Chi-0 | N1073 | | 1527 | 53.7502 | 34.7361 | Chisdra |
| CIBC-17 | CS76111 | 60 | | 51.4083 | -0.6383 | Ascot, Imperial College London - Silwood Park Campus |
| CIBC-2 | CS28140 | 423 | | 51.4083 | -0.6383 | Ascot, Imperial College London - Silwood Park Campus |
| CIBC-4 | CS28141 | 351 | | 51.4083 | -0.6383 | Ascot, Imperial College London - Silwood Park Campus |
| CIBC-5 | CS28142 | 377 | | 51.4083 | -0.6383 | Ascot, Imperial College London - Silwood Park Campus |
| Cit-0 | CS28158 | 378 | | 43.3779 | 2.54038 | Citou, Aude |
| Cl-0 | N1082 | | 1549 | (null) | (null) | |
| Cnt-1 | CS28160 | 398 | | 51.3 | 1.1 | Canterbury |
| Co | CS28161 | 342 | | 40.2077 | -8.42639 | Coimbra |
| Co-2 | CS28163 | 399 | | 40.12 | -8.25 | Coimbra |
| Co-3 | N1088 | 1261 | | 40.12 | -8.25 | Coimbra |
| Co-4 | CS28165 | 166 | | 40.12 | -8.25 | Coimbra |
| Col-0 | CS76113 | 139 | | 38.3 | -92.3 | Gückingen |

| name | stock nr | array ID call method 75 | array ID new | latitude | longitude | location |
|---|---|---|---|---|---|---|
| Com-1 | CS28193 | 352 | | 49.416 | 2.823 | Compiègne |
| CSHL-5 | CS28181 | 167 | | 40.8585 | -73.4675 | Cold Spring Harbor Laboratory, Long Island, NY |
| Ct-1 | CS76114 | 62 | | 37.3 | 15 | Catania |
| CUR-3 | CS76115 | 482 | | 45 | 1.75 | Curemonte |
| Cvi-0 | CS76116 | 143 | | 15.1111 | -23.6167 | Cap Verde Islands |
| Da(1)-12 | CS28201 | 221 | | (null) | (null) | |
| Da-0 | CS28200 | 400 | | 49.8724 | 8.65081 | Darmstadt |
| Db-0 | CS28202 | 446 | | 50.3055 | 8.324 | Dombachtal/Taunus |
| Db-1 | N1103 | | 1581 | 50.3055 | 8.324 | Dombachtal/Taunus |
| Di-1 | CS28208 | 168 | | 47 | 5 | Dijon |
| Dijon-M | N919 | | 1592 | 47 | 5 | Dijon |
| Do-0 | CS28210 | 447 | | 50.7224 | 8.2372 | Donsbach/Westerwald |
| Dr-0 | N1114 | 1047 | | 51.051 | 13.7336 | Dresden |
| Dra-0 | N1116 | | 1602 | 49.4167 | 16.2667 | Drahonin |
| Dra-2 | CS28214 | 424 | | 49.4167 | 16.2667 | Drahonin |
| DraIV1-14 | CS76119 | 430 | | 49.4112 | 16.2815 | Drahonín, Moravia |
| DraIV1-5 | CS76120 | 311 | | 49.4112 | 16.2815 | Drahonín, Moravia |
| DraIV6-16 | CS76122 | 312 | | 49.4112 | 16.2815 | Drahonín, Moravia |
| DraIV6-35 | CS76123 | 295 | | 49.4112 | 16.2815 | Drahonín, Moravia |
| Duk | CS76124 | 252 | | 49.1 | 16.2 | Dukovany, Moravia |
| Durh-1 | N22552 | | 1528 | 54.7761 | -1.5733 | Durham |
| Ede-1 | CS28217 | 448 | | 52.0333 | 5.66667 | Ede-Station |
| Eden-1 | CS28218 | 83 | | 62.877 | 18.177 | Eden |
| Edi-0 | CS76126 | 64 | | 56 | -3 | Edinburgh |
| Ei-2 | | | 1539 | 50.3 | 6.3 | Eifel |
| Ei-4 | CS28224 | 1049 | | 50.3 | 6.3 | Eifel |
| El-0 | N1134 | | 1589 | 51.5105 | 9.68253 | Ellershausen |
| En-1 | N1137 | 181 | | 50 | 8.5 | Enkheim |
| Enkheim-D | N920 | | 1550 | 50 | 8.5 | Enkheim |
| Ep-0 | CS28236 | 222 | | 50.1721 | 8.38912 | Eppenheim/Taunus |
| Er-0 | N1142 | | 1600 | 49.5955 | 11.0087 | Erlangen |
| Es-0 | CS28241 | 379 | | 60.1997 | 24.5682 | Espoo |
| Est-0 | CS28243 | 223 | | 58.3 | 25.3 | Estland |

| name | stock nr | array ID call method 75 | array ID new | latitude | longitude | location |
|---|---|---|---|---|---|---|
| Est-1 | CS76127 | 146 | | 58.3 | 25.3 | Estonia |
| Fei-0 | CS28250 | 147 | | 40.92 | -8.54 | Santa Maria da Feira |
| Fi-0 | N1157 | | 1561 | 50.5 | 8.0167 | Frickhofen |
| Fi-1 | CS28252 | 202 | | 50.5 | 8.0167 | Frickhofen |
| Fr-2 | N1169 | | 1572 | 50.1102 | 8.6822 | Frankfurt |
| Fr-4 | CS28268 | 449 | | 50.1102 | 8.6822 | Frankfurt |
| Ga-0 | CS76133 | 66 | | 50.3 | 8 | Gabelstein |
| Ga-2 | CS28274 | 401 | | 50.3 | 8 | Gabelstein |
| Gd-1 | CS76134 | 254 | | 53.5 | 10.5 | Gudow, SH |
| Ge-1 | CS28277 | 224 | | 46.5 | 6.08 | Geneva |
| Ge-2 | N1190 | | 1582 | 46.5 | 6.08 | Geneva |
| Gel-1 | CS28279 | 402 | | 51.0167 | 5.86667 | Geleen |
| Gie-0 | CS28280 | 225 | | 50.584 | 8.67825 | Gießen |
| Go-0 | CS28282 | 403 | | 51.5338 | 9.9355 | Göttingen |
| Gö-2 | N1196 | | 1525 | 51.5338 | 9.9355 | Göttingen |
| Golm-1 | | | 1593 | 52.416213 | 12.969162 | Golm, MPI |
| GOT-7 | CS22608 | | 1518 | 51.5338 | 9.9355 | Göttingen |
| Gr | | | 1529 | 47 | 15.5 | Graz, St |
| Gr-1 | CS76137 | 256 | | 47 | 15.5 | Graz, St |
| Gr-5 | CS28326 | 169 | | 47 | 15.5 | Graz, St |
| Gre-0 | N1210 | | 1536 | 43.178 | -85.2532 | Greenville, MI |
| Gu-0 | N22617 | 87 | | 50.3 | 8 | Gückingen |
| Gu-1 | CS28332 | 404 | | 50.3 | 8 | Gückingen |
| Gy-0 | CS76139 | 67 | | 49 | 2 | La Minière |
| Ha-0 | CS28336 | 170 | | 52.3721 | 9.73569 | Hannover |
| Hau-0 | CS28343 | 353 | | 55.675 | 12.5686 | Hauniensis |
| Hey-1 | CS28344 | 226 | | 51.25 | 5.9 | Heythuysen |
| Hh-0 | CS28345 | 354 | | 54.4175 | 9.88682 | Hohenlieth |
| Hi-0 | CS76140 | 1138 | | 52 | 5 | Hilversum |
| Hl-3 | N1232 | | 1540 | 52.1444 | 9.37827 | Holtensen |
| Hn-0 | CS28350 | 171 | | 51.3472 | 8.28844 | Hennetalsperre |
| Hod | CS76141 | 259 | | 48.8 | 17.1 | Hodonin |
| HOG | N922 | | 1551 | 38.717 | 69.712 | Obigarm |

| name | stock nr | array ID call method 75 | array ID new | latitude | longitude | location |
|---|---|---|---|---|---|---|
| Hoh-1 | | | 1562 | 48.7129 | 9.2116 | |
| Hovdala-2 | CS76143 | 570 | | 56.1 | 13.74 | Hovdala |
| HR-10 | N22597 | 88 | | 51.4083 | -0.6383 | Ascot, Imperial College London - Silwood Park Campus |
| HR-5 | CS28353 | 68 | | 51.4083 | -0.6383 | Ascot, Imperial College London - Silwood Park Campus |
| Hs-0 | CS76145 | 262 | | 52.24 | 9.44 | Hannover/Stroehen, NI |
| HSm | CS76146 | 263 | | 49.33 | 15.76 | Horni Smrcne |
| In-0 | CS76147 | 264 | | 47.5 | 11.5 | Isenburg/Innsbruck |
| Is-1 | N1242 | | 1583 | 50.5 | 7.5 | Isenburg/Neuwied |
| Je-0 | CS28364 | 355 | | 50.927 | 11.587 | Jena |
| Je-54 | N924 | | 1594 | 49 | 15 | Stankov |
| JEA | CS76148 | 439 | | 43.6833 | 7.33333 | Saint-Jean-Cap-Ferrat |
| Jea | 25AV | | 1519 | 43.41 | 7.2 | St Jean Cap Ferrat |
| Jl-3 | CS28369 | 405 | | 49.31 | 16.61 | Vranov/Brno |
| Jm-1 | CS28373 | 406 | | 49 | 15 | Jamolice |
| Kä-0 | CS76149 | 265 | | 47 | 14 | Kärnten |
| Kas-2 | CS76150 | 69 | | 35 | 77 | Kashmir |
| Kb-0 | N1269 | | 1541 | 50.183 | 8.5 | Kronberg/Taunus |
| KBS-Mac-8 | CS76151 | 390 | | 42.405 | -85.398 | Kellogg Biological Station, Michigan State University, Hickory Corners, MI |
| Kelsterbach-2 | CS28382 | 407 | | 50.0667 | 8.5333 | Kelsterbach, HE |
| Kelsterbach-4 | CS76152 | 249 | | 50.0667 | 8.5333 | Kelsterbach |
| Kil-0 | N1271 | 529 | | 55.6395 | -5.66364 | Killean |
| Kin-0 | CS76153 | 70 | | 43.362776 | -85.25472 | Kindalville, MI |
| Kl-0 | N1274 | 530 | | 50.95 | 6.9666 | Köln |
| Kl-5 | CS28394 | 302 | | 50.95 | 6.9666 | Köln |
| Kn-0 | CS28395 | 450 | | 54.8969 | 23.8924 | Kaunas |
| KNO-11 | CS28407 | 337 | | 41.2816 | -86.621 | Knox, IN |
| Kno-18 | CS76154 | 71 | | 41.2816 | -86.621 | Knox, IN |
| Koln | CS76155 | 266 | | 51 | 7 | Köln |
| Kondara | N916 | | 1558 | 38.48 | 68.49 | Kondara gorge |
| Kr-0 | CS28419 | 408 | | 51.3317 | 6.55934 | Krefeld |
| Kro-0 | CS28420 | 409 | | 50.0742 | 8.96617 | Klein-Krotzenburg |
| Krot-2 | CS28423 | 410 | | 49.631 | 11.5722 | Krottensee, Thüringen |
| Kz-1 | N22606 | 91 | | 49.5 | 73.1 | Karagandy |

| name | stock nr | array ID call method 75 | array ID new | latitude | longitude | location |
|---|---|---|---|---|---|---|
| Kz-9 | N22607 | 113 | | 49.5 | 73.1 | Karagandy |
| LAC-3 | CS76157 | 318 | | 47.7 | 6.81667 | Lachapelle-sous-Chaux |
| LAC-5 | CS76158 | 1074 | | 47.7 | 6.81667 | Lachapelle-sous-Chaux |
| Lan-0 | N1304 | | 1552 | 55.6739 | -3.78181 | Lanark |
| Laud-1 | N22555 | | 1563 | 55.7 | -2.75 | Lauder |
| Lc-0 | CS76159 | 268 | | 57 | -4 | Loch Ness |
| LDV-14 | CS76160 | 1072 | | 48.5167 | -4.06667 | Landivisiau |
| LDV-25 | CS76161 | 847 | | 48.5167 | -4.06667 | Landivisiau |
| LDV-34 | CS76162 | 433 | | 48.5167 | -4.06667 | Landivisiau |
| LDV-58 | CS76163 | 209 | | 48.5167 | -4.06667 | Landivisiau |
| Ler-1 | CS76164 | 150 | | 47.984 | 10.8719 | Landsberg am Lech |
| Li-3 | CS28454 | 303 | | 50.3833 | 8.0666 | Limburg |
| Li-5:2 | CS28457 | 380 | | 50.3833 | 8.0666 | Limburg |
| Li-6 | CS28459 | 381 | | 50.3833 | 8.0666 | Limburg |
| Li-7 | CS28461 | 411 | | 50.3833 | 8.0666 | Limburg |
| Liarum | CS76166 | 238 | | 55.95 | 13.85 | Liarum |
| Limeport | N8070 | | 1573 | 40.5088 | -75.4472 | Limeport, PA |
| Lip-0 | CS76168 | 270 | | 50 | 19.3 | Lipowiec/Chrzanow |
| Lis-1 | CS76169 | 271 | | 56 | 14.7 | Listershuvud |
| LL-0 | CS76172 | 72 | | 41.59 | 2.49 | Llagostera |
| Ll-OF-095 | CS76165 | 440 | | 40.7777 | -72.9069 | Brookhaven, NY - Organic Farm |
| Lm | | | 1595 | 48 | 0.5 | Le Mans |
| Lm-2 | CS76173 | 164 | | 48 | 0.5 | Le Mans |
| Lom1-1 | CS76174 | 275 | | 56.09 | 13.9 | Lommarp |
| Löv-5 | N22575 | | 1520 | 62.801 | 18.079 | Lövvik |
| Lp2-2 | CS76176 | 73 | | 49.38 | 16.81 | Lipovec/Brno |
| Lp2-6 | CS76177 | 74 | | 49.38 | 16.81 | Lipovec/Brno |
| Lu | | | 1569 | 55.71 | 13.2 | Lund |
| Lz-0 | CS76179 | 75 | | 46 | 3.3 | Lezoux |
| Map-42 | CS76180 | 368 | | 42.166 | -86.412 | Maple Lane, Benton Harbor, MI |
| Mc-0 | CS28490 | 356 | | 54.6167 | -2.3 | Mickle Fell |
| Me-0 | N1365 | 524 | | 51.9183 | 10.1138 | Merchtshausen, HE |
| Mh-0 | CS28492 | 227 | | 50.95 | 7.5 | Mühlen/Ostpreußen |

| name | stock nr | array ID call method 75 | array ID new | latitude | longitude | location |
|---|---|---|---|---|---|---|
| Mh-1 | N1368 | 536 | | 50.95 | 7.5 | Mühlen/Ostpreußen |
| MIB-15 | CS76181 | 431 | | 47.3833 | 5.31667 | Mirebeau-sur-Bèze |
| MIB-22 | CS76182 | 313 | | 47.3833 | 5.31667 | Mirebeau-sur-Bèze |
| MIB-28 | CS76183 | 236 | | 47.3833 | 5.31667 | Mirebeau-sur-Bèze |
| MIB-84 | CS76184 | 432 | | 47.3833 | 5.31667 | Mirebeau-sur-Bèze |
| MNF-Pot-48 | CS76187 | 321 | | 43.595 | -86.2657 | Manistee National Forest, Baldwin, MI |
| Mnz-0 | CS28495 | 412 | | 50.001 | 8.26664 | Mainz |
| MOG-37 | CS76189 | 1071 | | 48.6667 | -4.06667 | Sibiril |
| Mrk-0 | CS76191 | 77 | | 49 | 9.3 | Märkt/Baden |
| Ms-0 | N905 | | 1531 | 55.7522 | 37.6322 | Moskow |
| Mt-0 | CS76192 | 78 | | 32.34 | 22.46 | Martuba/Cyrenaika |
| Mz-0 | CS76193 | 92 | | 50.3 | 8.3 | Merzhausen/Taunus |
| N13 | CS76194 | 61 | | 61.36 | 34.15 | Konchezero, Karelien |
| N4 | CS28510 | 357 | | 61.84423 | 34.36777 | Solomennoye |
| N7 | CS28513 | 382 | | 61.36 | 34.15 | Pinguba |
| Na-1 | CS76195 | 277 | | 47.5 | 1.5 | Nantes |
| Nc-1 | CS28527 | 451 | | 48.6167 | 6.25 | Ville-en-Vermois |
| Nd | N1636 | | 1542 | 50 | 10 | Niederzenz/Arnstein |
| Nd-1 | CS76197 | 93 | | 50 | 10 | Niederzenz/Arnstein |
| NFA-10 | CS76198 | 94 | | 51.4083 | -0.6383 | Ascot |
| NFA-8 | CS76199 | 154 | | 51.4083 | -0.6383 | Ascot |
| NFC-20 | CS28550 | 383 | | 51.4083 | -0.6383 | Ascot |
| No-0 | CS28564 | 228 | | 51.0581 | 13.2995 | Nossen/Halle |
| Nok-1 | CS28568 | 358 | | 52.24 | 4.45 | Noordwijk |
| Nok-2 | N1402 | | 1564 | 52.24 | 4.45 | Noordwijk |
| Nw-0 | CS28573 | 229 | | 50.5 | 8.5 | Neuweilnau |
| Nw-2 | CS28575 | 413 | | 50.5 | 8.5 | Neuweilnau |
| NW3 | N1414 | | 1574 | 50.5 | 8.5 | Neuweilnau |
| Nz1 | CS28578 | 414 | | -37.7871 | 175.283 | Hamilton, NZ |
| Ob-1 | CS28580 | 304 | | 50.2 | 8.5833 | Oberursel/Friedhof |
| Old-1 | CS28583 | 415 | | 53.1667 | 8.2 | Oldenburg |
| Or-0 | CS28587 | 416 | | 50.3827 | 8.01161 | Oranienstein |
| Ors-1 | CS28848 | 387 | | 44.7203 | 22.3955 | Orsova |

| name | stock nr | array ID call method 75 | array ID new | latitude | longitude | location |
|---|---|---|---|---|---|---|
| Ors-2 | CS28849 | 341 | | 44.7203 | 22.3955 | Orsova |
| Ost-0 | CS28588 | 280 | | 60.25 | 18.37 | Osthammar |
| Ove-0 | N1434 | 1279 | | 53.3422 | 8.42255 | Ovelgoenne |
| Ove-0 | N1435 | | 1585 | 53.3422 | 8.42255 | Ovelgoenne |
| Oy-0 | CS76203 | 96 | | 60.23 | 6.13 | Oystese |
| Oy-1 | | | 1596 | 60.23 | 6.13 | Oystese |
| Pa-1 | N1439 | 281 | | 38.07 | 13.22 | Palermo |
| Pa-2 | CS28595 | 230 | | 38.07 | 13.22 | Palermo |
| PAR-3 | CS76205 | 316 | | 46.65 | -0.25 | Parthenay |
| PAR-4 | CS76206 | 436 | | 46.65 | -0.25 | Parthenay |
| Per-1 | CS76210 | 282 | | 58 | 56.3167 | Perm |
| Petergof | CS76211 | 283 | | 59 | 29 | Petergof |
| PHW-10 | CS28610 | 425 | | 51.29273 | 0.40907 | West Malling, Kent |
| PHW-13 | CS28613 | 426 | | 51.29273 | 0.40907 | West Malling, Kent |
| PHW-14 | CS28614 | 359 | | 51.29273 | 0.40907 | West Malling, Kent |
| PHW-20 | CS28620 | 305 | | 51.29273 | 0.40907 | West Malling, Kent |
| PHW-22 | CS28622 | 306 | | 51.4167 | -1.7167 | Marlborough |
| PHW-26 | CS28626 | 474 | | 50.6728 | -3.8404 | Chagford, Devon |
| PHW-28 | CS28628 | 338 | | 50.35 | -3.5833 | Dartmouth, Devon |
| PHW-31 | CS28631 | 475 | | 51.4666 | -3.2 | Ely, Norfolk |
| PHW-33 | CS28633 | 427 | | 52.25 | 4.5667 | Lisse, Keukenhof |
| PHW-35 | CS28635 | 339 | | 48.6103 | 2.3086 | Brétigny-sur-Orge |
| PHW-36 | CS28636 | 293 | | 48.6103 | 2.3086 | Brétigny-sur-Orge |
| PHW-37 | CS28637 | 428 | | 48.6103 | 2.3086 | Brétigny-sur-Orge |
| Pi-0 | N1455 | | 1521 | 47.04 | 10.51 | Pitztal/Tirol |
| Pla-0 | CS28640 | 307 | | 41.5 | 2.25 | Playa de Aro |
| Pn-0 | CS28645 | 203 | | 48.0653 | -2.96591 | Pontivy |
| Pna-10 | N22571 | 118 | | 42.0945 | -86.3253 | Benton Harbor, MI |
| Pna-17 | CS28647 | 119 | | 42.0945 | -86.3253 | Benton Harbor, MI |
| Pna-17 | N22570 | 1532/119 | | 42.0945 | -86.3253 | Benton Harbor, MI |
| Po-0 | N1470 | | 1543 | 50.7167 | 7.1 | Poppelsdorf |
| Pog-0 | CS28650 | 308 | | 49.2655 | -123.206 | Point Grey, BC |
| Pr-0 | CS28651 | 231 | | 50.1448 | 8.60706 | Frankfurt-Praunheim |

| name | stock nr | array ID call method 75 | array ID new | latitude | longitude | location |
|---|---|---|---|---|---|---|
| Pro-0 | CS28652 | 97 | | 43.25 | -6 | Proaza, Asturias |
| Pt-0 | N1478 | | 1554 | 53.476 | 10.6065 | Poetrau/Lauenburg |
| Pu2-23 | CS76215 | 98 | | 49.42 | 16.36 | Prudka/Doubravnik |
| PU2-24 | CS28663 | 384 | | 49.42 | 16.36 | Prudka/Doubravnik |
| Pu2-7 | N22592 | 120 | | 49.42 | 16.36 | Prudka/Doubravnik |
| Pyl-1 | 8AV | | 1579 | 44.39 | 1.1 | Le Pyla (33115) |
| Ra-0 | CS76216 | 99 | | 46 | 3.3 | Randan, Puy-de-Dome |
| Rak-2 | CS76217 | 284 | | 49 | 16 | Raksice/Krumlov |
| Ren-1 | CS76218 | 100 | | 48.5 | -1.41 | Rennes/Montanel |
| Ren-11 | N22611 | 121 | | 48.5 | -1.41 | Rennes |
| Rhen-1 | CS28685 | 360 | | 51.9667 | 5.56667 | Rhenen |
| Ri-0 | CS28686 | 1333 | | 49.1632 | -123.137 | Richmond, BC |
| RLD-1 | N913 | | 1590 | (null) | (null) | |
| RLD-2 | CS28688 | 1256 | | 56.25 | 34.3167 | Rschew |
| Rmx-A02 | N22568 | 122 | | 42.036 | -86.511 | St. Joseph, MI |
| Rmx-A180 | CS76220 | 101 | | 42.036 | -86.511 | St. Joseph, MI |
| Rou-0 | CS28692 | 385 | | 49.4424 | 1.09849 | Rouen |
| RRS-10 | CS22689 | 155 | | 41.5609 | -86.4251 | North Liberty, IN |
| RRS-7 | CS28713 | 156 | | 41.5609 | -86.4251 | North Liberty, IN |
| Rsch-0 | | | 1597 | 56.3 | 34 | Rschew/Starize |
| Rsch-4 | CS76222 | 285 | | 56.3 | 34 | Rschew/Starize |
| Rubeszhnoe | | | 1601 | 49 | 38.28 | Rubezhnoe |
| S96 | | 417 | | (null) | (null) | |
| Santa Clara | N8069 | | 1544 | 37.21 | -121.16 | Newman, CA |
| Sap-0 | CS76224 | 287 | | 49.49 | 14.24 | Slapy |
| Sapporo-0 | CS28724 | 232 | | 43.0553 | 141.346 | Sapporo |
| Sav-0 | CS28725 | 172 | | 49.1833 | 15.8833 | Slavice |
| Sav-0 | CS76225 | 288 | | 49.1833 | 15.8833 | Slavice |
| Se-0 | CS76226 | 102 | | 38.3333 | -3.53333 | San Eleno |
| Sei-0 | CS28729 | 233 | | 46.5438 | 11.5614 | Seis am Schlern/Siusi |
| Sg-1 | CS28732 | 477 | | 47.6667 | 9.5 | St. Georgen |
| Sh-0 | CS28734 | 204 | | 51.6832 | 10.2144 | Schwiegershausen |
| Shahdara | CS76227 | 157 | | 38.35 | 68.48 | Shakdara, Pamiro-Alai |

| name | stock nr | array ID call method 75 | array ID new | latitude | longitude | location |
|---|---|---|---|---|---|---|
| Si-0 | CS28739 | 205 | | 50.8738 | 8.02341 | Siegen |
| SLSP-30 | CS76228 | 212 | | 43.665 | -86.496 | Silver Lake State Park, Mears, MI |
| Sorbo | N931 | 123 | | 38.35 | 68.48 | Sorbo |
| Sp-0 | CS28743 | 294 | | 52.5339 | 13.181 | Berlin/Spandau |
| Sq-1 | N22600 | 126 | | 51.4083 | -0.6383 | Ascot |
| Sq-8 | CS76230 | 103 | | 51.4083 | -0.6383 | Ascot |
| St-0 | CS76231 | 289 | | 59 | 18 | Stockholm |
| Ste-0 | CS28750 | 478 | | 52.6058 | 11.8558 | Stendal |
| Ste-3 | CS76232 | 488 | | 42.03 | -86.514 | Stevensville, MI |
| Stw-0 | N1538 | | 1555 | 52 | 36 | Stobowa/Orel |
| Ta-0 | CS76242 | 290 | | 49.5 | 14.5 | Tabor |
| TAMM-2 | CS22604 | 159 | | 60 | 23.5 | Tammisari |
| Tamm-27 | N22605 | 127 | | 60 | 23.5 | Tammisari |
| TDr-1 | CS76245 | 322 | | 55.7683 | 14.1386 | Degeberga |
| TDr-3 | CS76248 | 1250 | | 55.7686 | 14.1381 | Degeberga |
| Te-0 | N1550 | | 1576 | 60.0585 | 23.2982 | Tenela |
| Tha-1 | CS28758 | 479 | | 52.08 | 4.3 | Den Haag |
| Ting-1 | CS28759 | 386 | | 56.5 | 14.9 | Tingsryd |
| Tiv-1 | CS28760 | 429 | | 41.96 | 12.8 | Tivoli |
| Tol-0 | N8020 | | 1587 | 41.6639 | -83.5553 | Toledo, OH |
| Tottarp-2 | CS76251 | 291 | | 55.95 | 13.85 | Tollarp |
| TOU-A1-115 | CS76252 | 177 | | 46.6667 | 4.11667 | Toulon-sur-Arroux |
| TOU-A1-116 | CS76253 | 1073 | | 46.6667 | 4.11667 | Toulon-sur-Arroux |
| TOU-A1-43 | CS76255 | 435 | | 46.6667 | 4.11667 | Toulon-sur-Arroux |
| TOU-A1-62 | CS76256 | 210 | | 46.6667 | 4.11667 | Toulon-sur-Arroux |
| TOU-A1-96 | CS76258 | 315 | | 46.6667 | 4.11667 | Toulon-sur-Arroux |
| TOU-C-3 | CS76259 | 437 | | 46.6667 | 4.11667 | Toulon-sur-Arroux |
| TOU-E-11 | CS76260 | 541 | | 46.6667 | 4.11667 | Toulon-sur-Arroux |
| TOU-H-13 | CS76262 | 317 | | 46.6667 | 4.11667 | Toulon-sur-Arroux |
| TOU-I-17 | CS76263 | 237 | | 46.6667 | 4.11667 | Toulon-sur-Arroux |
| TOU-I-2 | CS76264 | 365 | | 46.6667 | 4.11667 | Toulon-sur-Arroux |
| TOU-I-6 | CS76265 | 366 | | 46.6667 | 4.11667 | Toulon-sur-Arroux |
| Ts-1 | CS76268 | 161 | | 41.7194 | 2.93056 | Tossa de Mar |

| name | stock nr | array ID call method 75 | array ID new | latitude | longitude | location |
|---|---|---|---|---|---|---|
| Ts-5 | N22648 | 128 | | 41.7194 | 2.93056 | Tossa de Mar |
| Tscha-1 | CS28779 | 173 | | 47.0748 | 9.9042 | Tschagguns |
| Tsu-0 | CS28780 | 480 | | 34.43 | 136.31 | Tsushima |
| Tsu-1 | N1640 | 160 | | 34.43 | 136.31 | Tsushima |
| Tu-0 | N1567 | 334 | | 45 | 7.5 | Torino |
| Tul-0 | N1570 | | 1598 | 43.2708 | -85.2563 | Turk Lake/Greenville, MI |
| Ty-0 | CS28786 | 361 | | 56.4278 | -5.23439 | Taynuilt |
| Udul1-34 | CS76269 | 468 | | 49.2771 | 16.6314 | Utechov |
| Uk-1 | CS28787 | 206 | | 48.0333 | 7.7667 | Umkirch |
| Uk-2 | CS28788 | 165 | | 48.0333 | 7.7667 | Umkirch |
| Uk-4 | N1581 | | 1534 | 48.0333 | 7.7667 | Umkirch |
| UKID48 | CS76273 | 213 | | 54.7 | -2.7 | Lazonby |
| UKNW06-059 | CS76275 | 298 | | 54.4 | -3 | Ambleside, Lake District |
| UKNW06-060 | CS76276 | 299 | | 54.4 | -3 | Ambleside, Lake District |
| UKNW06-386 | CS76277 | 300 | | 54.6 | -3.1 | Keswick, Lake District |
| UKSE06-429 | CS76287 | 323 | | 51.3 | 0.4 | Hadlow |
| UKSE06-466 | CS76288 | 442 | | 51.2 | 0.4 | Paddock Wood |
| UKSE06-482 | CS76289 | 546 | | 51.2 | 0.6 | Staplehurst |
| UKSE06-520 | CS76290 | 547 | | 51.3 | 1.1 | Upper Harbledown |
| UKSE06-628 | CS76291 | 491 | | 51.1 | 0.4 | Scotney Castle |
| Ull2-3 | CS76293 | 104 | | 56.0648 | 13.9707 | Ullstorp, Skane |
| Ull-2-5 | CS28792 | 105 | | 56.0648 | 13.9707 | Ullstorp, Skane |
| Uod-1 | N22612 | 129 | | 48.3 | 14.45 | Ottenhof/Weingraben |
| Uod-7 | CS76296 | 106 | | 48.3 | 14.45 | Ottenhof/Weingraben |
| Utrecht | CS28795 | 418 | | 52.0918 | 5.1145 | Utrecht |
| Van-0 | CS76297 | 162 | | 49.3 | -123 | Vancouver, University of British Columbia, BC |
| Var-2-1 | CS28798 | 107 | | 55.58 | 14.334 | Varhallarna, Skane |
| Ven-1 | CS28800 | 481 | | 52.0333 | 5.55 | Veenendaal |
| Wa-1 | CS28804 | 174 | | 52.3 | 21 | Warsaw |
| Wa-1 | N22644 | 131 | | 52.3 | 21 | Warsaw |
| Wag-3 | CS28808 | 340 | | 51.9666 | 5.6666 | Wageningen-Asserpark |
| Wag-4 | CS28809 | 362 | | 51.9666 | 5.6666 | Wageningen -Genetics |
| Wag-5 | CS28810 | 419 | | 51.9666 | 5.6666 | Wageningen-trans |

| name | stock nr | array ID call method 75 | array ID new | latitude | longitude | location |
|---|---|---|---|---|---|---|
| WAR | CS28812 | 420 | | 41.7302 | -71.2825 | Lincoln Woods State Park, RI |
| Wc-2 | CS28814 | 207 | | 52.6 | 10.0667 | Westercelle |
| Wei-(1) | N1639 | | 1577 | 47.25 | 8.26 | Weiningen |
| Wei-0 | CS76301 | 108 | | 47.25 | 8.26 | Weiningen |
| Wil | CS6888 | | 1588 | 54.6833 | 25.3167 | Wilna/Towniskaia |
| Wil-2 | N1596 | | 1599 | 54.6833 | 25.3167 | Wilna/Towniskaia |
| Wl-0 | CS28822 | 235 | | 47.9299 | 10.8134 | Wildbad |
| Ws | CS28823 | 175 | | 52.3 | 30 | Wassilewskija |
| Ws-0 | N22623 | 109 | | 52.3 | 30 | Wassilewskija |
| Ws-2 | N22659 | 132 | | 52.3 | 30 | Wassilewskija |
| Ws-3 | N1638 | | 1524 | 52.3 | 30 | Wassilewskija |
| Wt-3 | CS28833 | 309 | | 52.3 | 9.3 | Wietze |
| Wt-5 | CS76304 | 110 | | 52.3 | 9.3 | Wietze |
| Yo-0 | CS28843 | 111 | | 37.45 | -119.35 | Yosemite Nat. Park |
| Zdr-1 | N22588 | 133 | | 49.3853 | 16.2544 | Zdarec/Brno, Moravia |
| Zdr-6 | CS28845 | 112 | | 49.3853 | 16.2544 | Zdarec/Brno, Moravia |
| Zdrl2-24 | CS76307 | 324 | | 49.3853 | 16.2544 | Zdarec/Brno, Moravia |
| Zdrl2-25 | CS76308 | 492 | | 49.3853 | 16.2544 | Zdarec/Brno, Moravia |
| Zü-1 | CS28847 | 310 | | 47.3667 | 8.55 | Zürich |
