## Supplementary Table S2 for "Temporal dynamics of QTL effects on vegetative growth in *Arabidopsis thaliana*"

| accession | DW20 | PLA07 | PLA08 | PLA09 | PLA10 | PLA11 | PLA12 | PLA13 | PLA14 | PLA15 | PLA16 | PLA17 | PLA18 | RGR07_09 |
|---|---|---|---|---|---|---|---|---|---|---|---|---|---|---|
| $H^2$ | 0.82 | 0.83 | 0.81 | 0.79 | 0.79 | 0.79 | 0.79 | 0.79 | 0.78 | 0.79 | 0.81 | 0.81 | 0.81 | 0.7508 |
| 11PNA4 | 10.04 | 6.16 | 9.06 | 13.72 | 20.35 | 32.50 | 49.63 | 74.01 | 108.60 | 156.61 | 217.89 | 293.76 | 370.38 | 0.391 |
| 328PNA | 10.04 | 6.15 | 8.45 | 12.01 | 17.95 | 30.49 | 47.63 | 70.29 | 104.40 | 142.84 | 202.98 | 271.19 | 322.15 | 0.317 |
| Aa-0 | 9.99 | 5.02 | 7.48 | 11.28 | 16.97 | 28.44 | 44.01 | 67.64 | 101.66 | 151.16 | 228.08 | 315.05 | 401.25 | 0.380 |
| Ak-1 | 11.10 | 6.17 | 8.61 | 13.27 | 20.02 | 30.66 | 46.68 | 68.69 | 101.16 | 150.81 | 224.16 | 309.58 | 405.89 | 0.369 |
| Akita | 11.37 | 5.62 | 8.09 | 11.89 | 19.31 | 30.83 | 48.34 | 72.75 | 108.67 | 164.28 | 246.29 | 335.16 | 445.82 | 0.362 |
| Alc-0 | 10.01 | 5.29 | 7.69 | 11.56 | 17.67 | 28.69 | 45.14 | 69.13 | 99.62 | 145.40 | 205.88 | 268.25 | 325.25 | 0.380 |
| ALL1-2 | 10.49 | 6.70 | 9.64 | 14.51 | 22.27 | 35.61 | 53.61 | 79.93 | 119.44 | 171.90 | 246.72 | 334.45 | 425.16 | 0.375 |
| Alst-1 | 11.79 | 7.08 | 10.60 | 16.05 | 24.74 | 38.73 | 58.49 | 88.35 | 132.07 | 192.07 | 259.33 | 338.21 | 419.78 | 0.396 |
| Amel-1 | 11.30 | 7.36 | 10.20 | 14.96 | 22.04 | 34.37 | 51.59 | 76.57 | 109.49 | 160.34 | 229.91 | 319.60 | 414.88 | 0.350 |
| An1 | 11.04 | 5.27 | 7.61 | 11.62 | 17.77 | 28.70 | 44.51 | 67.63 | 99.21 | 151.44 | 225.33 | 303.61 | 394.24 | 0.383 |
| An-2 | 11.24 | 5.22 | 7.76 | 12.30 | 19.07 | 29.77 | 45.25 | 66.88 | 101.08 | 151.17 | 216.49 | 284.17 | 356.11 | 0.417 |
| Ang-0 | 9.06 | 4.90 | 7.11 | 11.24 | 16.79 | 27.43 | 42.38 | 63.80 | 95.24 | 141.55 | 206.23 | 286.60 | 369.79 | 0.385 |
| Ang-0 | 13.91 | 7.53 | 10.79 | 16.15 | 25.45 | 40.38 | 61.82 | 93.26 | 143.33 | 214.79 | 312.06 | 415.34 | 505.67 | 0.373 |
| Ang-1 | 10.24 | 5.86 | 8.46 | 12.92 | 20.42 | 32.79 | 49.50 | 73.44 | 107.33 | 161.61 | 237.46 | 324.33 | 400.72 | 0.387 |
| Ann-1 | 13.76 | 7.16 | 10.47 | 15.81 | 24.63 | 40.50 | 64.89 | 100.70 | 152.38 | 222.35 | 308.13 | 395.03 | 480.73 | 0.384 |
| Arby-1 | 11.50 | 6.04 | 9.22 | 14.11 | 20.75 | 33.10 | 51.56 | 76.76 | 116.52 | 180.80 | 271.28 | 381.25 | 485.67 | 0.400 |
| Ba-1 | 11.15 | 5.80 | 8.38 | 11.91 | 18.69 | 31.54 | 49.71 | 77.16 | 115.65 | 179.06 | 266.43 | 360.13 | 438.29 | 0.351 |
| Ba1-2 | 14.02 | 7.00 | 10.27 | 16.01 | 25.15 | 40.27 | 62.24 | 95.86 | 144.93 | 221.63 | 330.13 | 446.89 | 576.60 | 0.401 |
| Baa-1 | 12.42 | 6.66 | 9.79 | 14.58 | 22.18 | 35.51 | 55.14 | 84.09 | 125.61 | 185.37 | 273.82 | 375.33 | 473.84 | 0.370 |
| Bay-0 | 13.06 | 6.39 | 9.26 | 14.54 | 22.67 | 36.53 | 56.21 | 85.77 | 129.48 | 192.84 | 273.70 | 371.45 | 460.68 | 0.403 |
| Bch-1 | 14.65 | 6.42 | 9.43 | 14.64 | 22.65 | 36.96 | 58.27 | 88.69 | 134.01 | 208.12 | 314.11 | 425.67 | 523.06 | 0.400 |
| Bd-0 | 8.98 | 4.98 | 7.08 | 10.39 | 15.98 | 25.59 | 40.04 | 60.51 | 89.23 | 129.99 | 193.31 | 279.09 | 371.14 | 0.358 |
| Be-0 | 9.85 | 6.43 | 9.30 | 13.62 | 20.49 | 32.85 | 49.85 | 72.86 | 106.91 | 166.00 | 248.15 | 352.26 | 461.89 | 0.360 |
| Be-1 | 11.63 | 5.76 | 8.40 | 12.79 | 19.93 | 32.22 | 49.98 | 76.54 | 115.61 | 179.06 | 269.42 | 377.53 | 487.99 | 0.387 |
| Belmonte4 | 8.80 | 6.45 | 9.20 | 13.99 | 20.62 | 31.34 | 46.35 | 68.46 | 99.83 | 145.81 | 212.19 | 297.05 | 388.05 | 0.370 |
| Benk-1 | 12.11 | 7.89 | 10.75 | 15.29 | 23.51 | 36.97 | 55.67 | 83.45 | 122.42 | 180.89 | 257.96 | 337.12 | 423.63 | 0.319 |
| Bg-2 | 10.13 | 6.13 | 8.44 | 12.16 | 18.19 | 28.74 | 43.27 | 63.78 | 95.63 | 147.84 | 220.59 | 315.89 | 434.89 | 0.336 |
| Bl-1 | 10.61 | 5.92 | 8.54 | 11.96 | 19.17 | 30.75 | 47.22 | 71.87 | 105.95 | 154.83 | 230.46 | 323.05 | 400.99 | 0.344 |
| Bla-1 | 10.82 | 6.51 | 9.34 | 14.08 | 21.00 | 33.36 | 50.93 | 77.16 | 112.51 | 166.10 | 232.22 | 302.24 | 361.37 | 0.368 |

| accession | DW20 | PLA07 | PLA08 | PLA09 | PLA10 | PLA11 | PLA12 | PLA13 | PLA14 | PLA15 | PLA16 | PLA17 | PLA18 | RGR07_09 |
|---|---|---|---|---|---|---|---|---|---|---|---|---|---|---|
| Bla-11 | 9.34 | 5.20 | 7.85 | 11.61 | 17.33 | 27.86 | 42.67 | 64.47 | 94.73 | 141.34 | 209.48 | 294.34 | 388.54 | 0.387 |
| Blh-1 | 10.20 | 5.46 | 8.28 | 12.68 | 19.44 | 31.59 | 49.81 | 74.89 | 112.64 | 167.48 | 240.12 | 315.81 | 398.32 | 0.403 |
| Blh-1 | 13.27 | 8.61 | 12.21 | 17.47 | 27.09 | 42.64 | 65.55 | 98.41 | 148.54 | 217.74 | 305.46 | 393.07 | 468.65 | 0.348 |
| Blh-2 | 10.47 | 6.73 | 9.58 | 13.73 | 20.94 | 33.37 | 52.32 | 78.61 | 116.59 | 176.30 | 261.45 | 349.82 | 442.51 | 0.347 |
| Boot-1 | 11.48 | 6.74 | 9.75 | 15.03 | 22.24 | 34.24 | 53.14 | 81.22 | 122.68 | 182.83 | 269.84 | 369.13 | 482.77 | 0.370 |
| Bor-1 | 10.48 | 6.43 | 9.37 | 14.19 | 21.42 | 32.99 | 49.40 | 72.79 | 104.41 | 151.51 | 227.31 | 321.65 | 434.71 | 0.385 |
| Bor-4 | 12.00 | 5.73 | 8.58 | 13.14 | 20.39 | 33.00 | 50.97 | 77.16 | 117.10 | 183.40 | 269.05 | 367.02 | 463.10 | 0.397 |
| Br-0 | 17.29 | 9.77 | 14.42 | 21.67 | 34.43 | 55.79 | 86.74 | 132.26 | 201.58 | 294.57 | 396.33 | 493.57 | 586.39 | 0.392 |
| Bs-1 | 9.99 | 5.94 | 8.53 | 12.29 | 18.55 | 29.06 | 44.41 | 65.94 | 94.41 | 143.66 | 221.45 | 322.28 | 427.14 | 0.351 |
| Bs-2 | 13.64 | 7.49 | 10.71 | 15.58 | 23.61 | 37.08 | 56.47 | 85.85 | 129.84 | 193.58 | 279.10 | 380.60 | 495.85 | 0.353 |
| Bsch-0 | 11.18 | 5.86 | 8.41 | 12.79 | 20.33 | 33.39 | 51.69 | 78.09 | 115.70 | 171.89 | 259.36 | 357.22 | 451.15 | 0.378 |
| Bsch-2 | 15.39 | 8.00 | 11.41 | 17.38 | 27.66 | 43.93 | 66.61 | 100.81 | 153.15 | 228.88 | 328.98 | 435.96 | 537.00 | 0.378 |
| Bu-0 | 10.28 | 6.27 | 9.19 | 13.75 | 20.90 | 33.85 | 53.48 | 80.29 | 118.78 | 172.12 | 241.53 | 318.73 | 399.65 | 0.382 |
| Bu-2 | 11.77 | 6.53 | 9.44 | 14.09 | 21.37 | 34.54 | 53.14 | 79.64 | 119.49 | 175.50 | 259.41 | 341.10 | 432.16 | 0.374 |
| Bur-0 | 10.91 | 8.15 | 10.88 | 15.02 | 22.76 | 35.22 | 52.37 | 75.83 | 108.48 | 157.22 | 233.53 | 321.83 | 424.88 | 0.291 |
| C24 | 10.06 | 6.76 | 9.90 | 14.57 | 21.44 | 33.88 | 51.07 | 74.28 | 109.59 | 163.50 | 239.81 | 327.59 | 438.59 | 0.377 |
| Ca-0 | 13.21 | 7.76 | 11.43 | 16.96 | 25.25 | 39.86 | 60.89 | 91.21 | 136.55 | 198.84 | 286.59 | 386.01 | 480.19 | 0.375 |
| Cal-0 | 15.85 | 10.15 | 13.88 | 19.77 | 30.85 | 47.98 | 71.90 | 105.14 | 152.81 | 227.46 | 325.49 | 439.79 | 551.47 | 0.326 |
| CAM-16 | 9.49 | 5.41 | 8.14 | 12.29 | 18.74 | 29.66 | 44.79 | 65.72 | 99.50 | 152.77 | 227.57 | 317.26 | 421.39 | 0.395 |
| CAM-61 | 11.08 | 5.77 | 8.43 | 12.74 | 19.09 | 29.28 | 44.61 | 67.14 | 99.35 | 148.60 | 214.97 | 291.21 | 372.50 | 0.375 |
| Can-0 | 11.66 | 7.08 | 10.22 | 16.10 | 24.65 | 38.26 | 58.39 | 87.54 | 133.37 | 202.85 | 290.81 | 386.44 | 473.45 | 0.401 |
| Cen-0 | 6.90 | 4.72 | 6.66 | 9.32 | 13.83 | 23.58 | 36.15 | 53.88 | 78.93 | 117.22 | 177.96 | 253.52 | 342.28 | 0.327 |
| Cha-0 | 12.48 | 7.64 | 10.99 | 16.35 | 25.41 | 40.35 | 62.30 | 94.65 | 139.90 | 208.36 | 300.59 | 394.14 | 458.36 | 0.372 |
| Chat-1 | 10.04 | 4.95 | 7.08 | 10.21 | 15.62 | 25.22 | 39.55 | 60.57 | 92.10 | 139.97 | 205.27 | 286.52 | 372.00 | 0.350 |
| Chi-0 | 12.82 | 6.37 | 9.38 | 14.58 | 22.30 | 35.47 | 55.89 | 85.13 | 129.14 | 195.26 | 288.70 | 395.02 | 498.83 | 0.390 |
| CIBC-17 | 11.88 | 6.90 | 9.93 | 14.33 | 22.41 | 35.78 | 54.58 | 82.93 | 125.67 | 187.37 | 270.45 | 359.87 | 470.55 | 0.354 |
| CIBC-2 | 12.14 | 7.09 | 10.14 | 15.04 | 23.59 | 38.96 | 60.63 | 90.99 | 133.48 | 192.87 | 267.87 | 351.81 | 444.32 | 0.370 |
| CIBC-4 | 11.37 | 7.33 | 10.72 | 15.39 | 24.13 | 38.03 | 57.89 | 86.26 | 128.41 | 192.10 | 274.59 | 360.43 | 449.89 | 0.363 |
| CIBC-5 | 7.66 | 5.22 | 7.68 | 11.82 | 17.76 | 28.34 | 43.25 | 64.38 | 93.06 | 135.37 | 197.65 | 266.68 | 332.54 | 0.379 |
| Cit-0 | 9.64 | 5.53 | 7.95 | 11.65 | 17.20 | 27.10 | 41.20 | 62.16 | 94.22 | 137.70 | 204.91 | 284.40 | 369.73 | 0.360 |

| accession | DW20 | PLA07 | PLA08 | PLA09 | PLA10 | PLA11 | PLA12 | PLA13 | PLA14 | PLA15 | PLA16 | PLA17 | PLA18 | RGR07_09 |
|---|---|---|---|---|---|---|---|---|---|---|---|---|---|---|
| Cl-0 | 14.11 | 7.97 | 11.76 | 17.70 | 27.39 | 42.84 | 64.14 | 97.63 | 150.65 | 225.99 | 319.54 | 421.50 | 539.82 | 0.392 |
| Cnt-1 | 10.96 | 7.34 | 10.52 | 15.61 | 22.70 | 34.53 | 51.83 | 77.35 | 115.85 | 172.46 | 252.38 | 346.17 | 456.09 | 0.364 |
| Co | 7.24 | 4.25 | 6.12 | 9.49 | 13.83 | 21.99 | 34.16 | 51.31 | 76.45 | 114.16 | 172.83 | 250.41 | 333.20 | 0.378 |
| Co-2 | 12.25 | 7.06 | 10.24 | 14.76 | 23.90 | 38.57 | 59.71 | 90.04 | 132.88 | 197.19 | 281.80 | 376.08 | 476.82 | 0.356 |
| Co-3 | 8.24 | 4.52 | 6.71 | 10.14 | 15.70 | 25.07 | 38.33 | 57.24 | 85.16 | 127.25 | 192.68 | 278.96 | 380.71 | 0.394 |
| Co-4 | 12.28 | 6.99 | 10.73 | 15.30 | 23.16 | 35.99 | 53.67 | 81.61 | 120.41 | 180.73 | 263.11 | 364.55 | 474.70 | 0.378 |
| Col-0 | 11.06 | 5.79 | 8.34 | 12.51 | 19.71 | 31.45 | 48.17 | 72.66 | 109.13 | 161.04 | 239.98 | 328.28 | 416.86 | 0.371 |
| Com-1 | 8.67 | 4.97 | 7.32 | 10.86 | 16.19 | 25.24 | 37.43 | 55.91 | 81.73 | 122.16 | 184.19 | 265.01 | 341.60 | 0.373 |
| CSHL-5 | 9.72 | 4.96 | 7.24 | 11.24 | 17.25 | 27.43 | 41.69 | 65.42 | 98.02 | 146.61 | 219.28 | 310.09 | 377.66 | 0.389 |
| Ct-1 | 12.96 | 6.20 | 9.62 | 14.72 | 22.74 | 36.02 | 55.92 | 85.81 | 128.06 | 182.69 | 255.53 | 338.94 | 411.97 | 0.420 |
| CUR-3 | 5.98 | 5.80 | 8.33 | 11.08 | 15.66 | 24.95 | 35.09 | 49.01 | 68.49 | 99.52 | 147.82 | 219.26 | 326.04 | 0.319 |
| Cvi-0 | 5.82 | 5.60 | 8.61 | 12.20 | 16.65 | 24.04 | 34.40 | 47.02 | 65.81 | 89.60 | 126.09 | 182.38 | 253.80 | 0.360 |
| Da(1)-12 | 12.35 | 6.25 | 9.12 | 13.98 | 22.44 | 36.18 | 55.62 | 83.74 | 122.49 | 179.50 | 265.87 | 355.55 | 438.76 | 0.390 |
| Da-0 | 11.56 | 5.97 | 8.87 | 13.26 | 20.08 | 32.84 | 51.75 | 80.99 | 120.97 | 176.01 | 254.57 | 350.22 | 453.53 | 0.375 |
| Db-0 | 8.86 | 4.46 | 6.63 | 10.31 | 15.46 | 25.45 | 40.34 | 61.66 | 93.47 | 139.22 | 212.14 | 300.91 | 401.94 | 0.399 |
| Db-1 | 11.74 | 6.14 | 9.06 | 14.10 | 22.07 | 34.90 | 54.13 | 81.04 | 121.75 | 176.84 | 255.65 | 350.01 | 441.32 | 0.398 |
| Di-1 | 10.37 | 6.81 | 9.94 | 14.47 | 21.19 | 33.09 | 49.83 | 72.14 | 109.68 | 167.68 | 246.36 | 336.96 | 449.95 | 0.365 |
| Dijon-M | 11.41 | 5.10 | 7.73 | 11.97 | 18.36 | 29.22 | 44.62 | 69.77 | 103.46 | 151.48 | 218.62 | 309.51 | 380.46 | 0.412 |
| Do-0 | 13.74 | 8.03 | 11.92 | 18.34 | 28.43 | 44.96 | 69.26 | 104.99 | 158.51 | 231.99 | 319.85 | 416.32 | 505.99 | 0.403 |
| Dr-0 | 14.12 | 6.96 | 9.69 | 14.77 | 23.81 | 38.84 | 60.28 | 91.63 | 137.97 | 206.08 | 305.51 | 414.20 | 496.68 | 0.361 |
| Dra-0 | 12.00 | 5.98 | 8.77 | 13.77 | 21.59 | 35.01 | 53.16 | 78.73 | 113.70 | 167.77 | 247.12 | 329.56 | 404.97 | 0.410 |
| Dra-2 | 9.75 | 5.12 | 7.58 | 11.03 | 17.63 | 27.93 | 42.36 | 62.26 | 93.89 | 142.58 | 212.47 | 301.53 | 388.14 | 0.367 |
| DraIV1-14 | 10.24 | 5.91 | 8.99 | 13.90 | 21.55 | 34.09 | 52.23 | 78.15 | 115.20 | 172.86 | 252.20 | 339.30 | 442.64 | 0.416 |
| DraIV1-5 | 11.39 | 6.35 | 9.54 | 13.77 | 22.42 | 36.39 | 56.84 | 85.61 | 125.69 | 182.93 | 263.41 | 352.64 | 443.68 | 0.366 |
| DraIV6-16 | 14.64 | 8.09 | 11.72 | 17.41 | 26.78 | 42.22 | 64.67 | 99.23 | 150.54 | 221.32 | 315.49 | 420.67 | 524.41 | 0.369 |
| DraIV6-35 | 14.41 | 7.05 | 10.28 | 16.06 | 26.45 | 42.01 | 62.62 | 94.05 | 143.28 | 213.10 | 294.44 | 394.01 | 492.33 | 0.388 |
| Duk | 12.46 | 6.45 | 9.71 | 15.02 | 23.40 | 37.03 | 57.18 | 86.70 | 129.75 | 197.99 | 288.27 | 397.76 | 518.89 | 0.405 |
| Durh-1 | 6.82 | 5.98 | 8.64 | 12.28 | 17.45 | 26.04 | 38.75 | 55.80 | 80.22 | 113.97 | 159.92 | 218.84 | 298.45 | 0.355 |
| Ede-1 | 10.81 | 5.95 | 8.72 | 13.22 | 19.98 | 31.05 | 46.91 | 70.84 | 103.79 | 156.75 | 236.82 | 325.81 | 421.36 | 0.385 |
| Eden-1 | 9.76 | 7.09 | 10.24 | 15.44 | 22.89 | 36.11 | 54.38 | 80.42 | 115.15 | 161.23 | 227.12 | 302.95 | 376.25 | 0.374 |

| accession | DW20 | PLA07 | PLA08 | PLA09 | PLA10 | PLA11 | PLA12 | PLA13 | PLA14 | PLA15 | PLA16 | PLA17 | PLA18 | RGR07_09 |
|---|---|---|---|---|---|---|---|---|---|---|---|---|---|---|
| Edi-0 | 9.77 | 6.41 | 9.40 | 13.82 | 22.23 | 35.42 | 54.01 | 80.76 | 121.84 | 177.24 | 256.42 | 340.09 | 425.30 | 0.367 |
| Ei-2 | 12.58 | 6.25 | 9.22 | 14.28 | 22.03 | 34.91 | 54.01 | 81.92 | 121.05 | 185.59 | 274.93 | 369.99 | 481.95 | 0.391 |
| Ei-4 | 13.47 | 6.58 | 9.80 | 15.65 | 23.33 | 37.17 | 56.77 | 85.16 | 130.16 | 193.56 | 277.21 | 369.83 | 469.04 | 0.402 |
| El-0 | 11.78 | 6.21 | 9.45 | 14.36 | 22.34 | 36.57 | 55.83 | 81.44 | 120.25 | 178.61 | 259.81 | 362.32 | 452.05 | 0.407 |
| En-1 | 13.57 | 6.48 | 10.14 | 15.32 | 23.55 | 36.87 | 56.14 | 87.29 | 129.63 | 187.30 | 265.94 | 354.80 | 429.68 | 0.414 |
| Enkheim-D | 10.76 | 5.37 | 8.20 | 12.33 | 18.46 | 30.01 | 47.27 | 72.45 | 107.80 | 164.62 | 244.20 | 331.78 | 423.79 | 0.400 |
| Ep-0 | 13.61 | 6.16 | 9.13 | 14.16 | 21.52 | 34.86 | 54.31 | 83.79 | 127.46 | 193.38 | 276.94 | 358.18 | 441.91 | 0.394 |
| Er-0 | 11.44 | 5.51 | 8.27 | 12.98 | 19.97 | 32.32 | 50.62 | 76.85 | 117.84 | 181.80 | 264.07 | 359.76 | 470.46 | 0.409 |
| Es-0 | 12.35 | 6.66 | 9.66 | 14.90 | 23.59 | 38.30 | 59.74 | 91.01 | 134.39 | 205.82 | 292.53 | 376.35 | 459.23 | 0.391 |
| Est-0 | 11.19 | 5.59 | 8.13 | 12.69 | 20.29 | 33.14 | 51.29 | 78.37 | 118.21 | 177.53 | 258.97 | 356.79 | 460.43 | 0.364 |
| Est-1 | 14.22 | 6.31 | 9.45 | 13.97 | 22.07 | 36.61 | 57.57 | 89.36 | 136.05 | 204.97 | 302.17 | 400.39 | 490.59 | 0.396 |
| Fei-0 | 12.24 | 6.14 | 9.04 | 13.95 | 20.61 | 32.89 | 50.94 | 77.44 | 117.44 | 178.25 | 256.06 | 337.44 | 427.05 | 0.377 |
| Fi-0 | 10.48 | 5.92 | 8.32 | 12.03 | 18.48 | 29.26 | 44.81 | 66.79 | 97.98 | 147.71 | 214.95 | 302.69 | 401.34 | 0.348 |
| Fi-1 | 12.45 | 6.70 | 10.01 | 15.07 | 23.34 | 37.43 | 57.83 | 87.82 | 131.85 | 197.82 | 288.02 | 387.17 | 484.37 | 0.392 |
| Fr-2 | 13.95 | 6.22 | 9.73 | 13.36 | 22.08 | 35.57 | 54.50 | 82.08 | 121.82 | 180.18 | 263.36 | 356.40 | 459.22 | 0.377 |
| Fr-4 | 11.84 | 5.94 | 8.86 | 13.55 | 20.96 | 33.79 | 53.61 | 81.30 | 122.85 | 182.08 | 260.87 | 345.60 | 443.19 | 0.396 |
| Ga-0 | 12.11 | 5.55 | 8.36 | 12.49 | 19.06 | 30.54 | 47.73 | 72.17 | 108.80 | 166.31 | 251.81 | 358.85 | 477.49 | 0.390 |
| Ga-2 | 10.00 | 4.88 | 7.07 | 10.88 | 16.43 | 26.70 | 42.24 | 65.76 | 98.51 | 149.59 | 224.57 | 315.74 | 405.92 | 0.383 |
| Gd-1 | 13.04 | 5.82 | 8.54 | 13.15 | 20.65 | 33.71 | 52.81 | 81.35 | 124.33 | 185.11 | 273.18 | 355.62 | 443.29 | 0.401 |
| Ge-1 | 11.25 | 5.30 | 8.11 | 12.45 | 19.04 | 29.98 | 45.00 | 69.92 | 106.30 | 156.92 | 230.63 | 319.37 | 413.59 | 0.400 |
| Ge-2 | 10.79 | 5.87 | 8.90 | 13.65 | 20.48 | 33.40 | 51.64 | 75.84 | 109.25 | 163.04 | 236.45 | 307.88 | 379.23 | 0.409 |
| Gel-1 | 10.79 | 5.56 | 8.41 | 12.34 | 19.50 | 31.17 | 48.52 | 74.13 | 110.83 | 166.82 | 252.36 | 353.88 | 452.16 | 0.381 |
| Gie-0 | 11.95 | 6.05 | 8.67 | 13.67 | 20.75 | 33.38 | 51.39 | 78.15 | 118.13 | 177.72 | 262.94 | 368.81 | 491.91 | 0.387 |
| Go-0 | 9.69 | 5.09 | 7.84 | 12.11 | 18.31 | 28.35 | 43.40 | 65.42 | 100.20 | 149.81 | 227.68 | 325.19 | 439.13 | 0.417 |
| Gö-2 | 10.70 | 5.47 | 7.78 | 11.99 | 18.06 | 28.78 | 45.03 | 68.61 | 102.11 | 154.47 | 228.94 | 324.89 | 425.15 | 0.370 |
| Golm-1 | 11.45 | 5.35 | 7.75 | 10.98 | 17.83 | 29.81 | 47.07 | 72.28 | 110.68 | 173.21 | 265.38 | 371.65 | 486.16 | 0.345 |
| GOT-7 | 11.57 | 7.64 | 10.63 | 14.94 | 22.11 | 34.92 | 53.08 | 79.74 | 114.55 | 160.67 | 232.18 | 307.54 | 373.20 | 0.331 |
| Gr | 8.48 | 5.12 | 7.32 | 11.31 | 16.47 | 26.22 | 40.52 | 61.64 | 90.50 | 136.69 | 204.68 | 291.91 | 388.75 | 0.369 |
| Gr-1 | 8.42 | 4.48 | 6.56 | 10.15 | 15.09 | 23.93 | 37.35 | 56.55 | 84.71 | 127.17 | 191.51 | 268.55 | 368.49 | 0.389 |
| Gr-5 | 12.10 | 6.05 | 8.66 | 11.94 | 20.16 | 31.14 | 48.24 | 72.56 | 112.94 | 174.09 | 263.99 | 371.58 | 478.24 | 0.319 |

| accession | DW20 | PLA07 | PLA08 | PLA09 | PLA10 | PLA11 | PLA12 | PLA13 | PLA14 | PLA15 | PLA16 | PLA17 | PLA18 | RGR07_09 |
|---|---|---|---|---|---|---|---|---|---|---|---|---|---|---|
| Gre-0 | 12.03 | 7.37 | 10.57 | 15.59 | 23.66 | 36.89 | 55.99 | 83.79 | 126.00 | 183.91 | 262.26 | 347.63 | 422.81 | 0.362 |
| Gu-0 | 11.58 | 5.86 | 8.18 | 10.80 | 18.43 | 30.42 | 47.31 | 70.99 | 105.87 | 160.01 | 235.58 | 317.87 | 409.33 | 0.294 |
| Gu-1 | 10.74 | 5.38 | 7.91 | 11.74 | 17.88 | 28.46 | 44.03 | 67.24 | 101.16 | 154.18 | 237.71 | 334.71 | 434.13 | 0.388 |
| Gy-0 | 8.51 | 6.38 | 8.79 | 13.02 | 19.68 | 30.74 | 45.82 | 66.67 | 96.72 | 141.80 | 202.85 | 273.42 | 368.61 | 0.343 |
| Ha-0 | 13.79 | 6.21 | 9.89 | 15.77 | 24.91 | 39.70 | 61.16 | 91.92 | 141.85 | 213.40 | 301.33 | 383.84 | 467.99 | 0.446 |
| Hau-0 | 13.37 | 6.72 | 9.93 | 14.84 | 22.34 | 34.90 | 52.63 | 79.50 | 118.42 | 176.41 | 258.24 | 342.15 | 422.22 | 0.385 |
| Hey-1 | 11.28 | 6.44 | 9.13 | 13.70 | 20.65 | 33.36 | 51.18 | 77.15 | 114.97 | 172.57 | 254.18 | 342.02 | 424.92 | 0.366 |
| Hh-0 | 9.10 | 4.47 | 6.56 | 10.40 | 15.25 | 24.43 | 37.97 | 58.29 | 87.76 | 130.90 | 195.27 | 270.27 | 344.62 | 0.394 |
| Hi-0 | 8.02 | 4.19 | 5.98 | 9.23 | 13.70 | 21.66 | 33.09 | 49.22 | 72.47 | 110.59 | 168.89 | 246.88 | 335.32 | 0.361 |
| Hl-3 | 11.88 | 5.66 | 8.61 | 13.38 | 20.72 | 33.57 | 52.47 | 80.15 | 119.54 | 178.43 | 262.07 | 356.13 | 458.08 | 0.416 |
| Hn-0 | 9.70 | 6.53 | 9.06 | 13.76 | 20.61 | 31.90 | 47.56 | 69.86 | 101.91 | 147.74 | 217.92 | 302.73 | 401.53 | 0.360 |
| Hod | 11.67 | 7.29 | 10.48 | 15.53 | 24.17 | 38.09 | 58.59 | 87.78 | 128.22 | 188.95 | 282.84 | 401.97 | 525.91 | 0.373 |
| HOG | 11.54 | 5.42 | 8.31 | 12.71 | 19.87 | 31.94 | 50.05 | 76.67 | 114.33 | 167.40 | 249.57 | 340.39 | 420.20 | 0.417 |
| Hoh-1 | 8.30 | 4.02 | 5.61 | 8.25 | 12.21 | 19.69 | 30.13 | 45.62 | 69.14 | 104.03 | 157.17 | 230.68 | 329.50 | 0.346 |
| Hovdala-2 | 11.30 | 5.62 | 8.04 | 11.74 | 18.25 | 30.80 | 48.51 | 73.13 | 109.34 | 161.94 | 232.21 | 316.22 | 393.84 | 0.343 |
| HR-10 | 10.37 | 5.46 | 8.11 | 12.48 | 18.50 | 28.83 | 44.11 | 66.58 | 99.12 | 150.49 | 221.56 | 311.61 | 408.71 | 0.396 |
| HR-5 | 9.68 | 5.15 | 7.53 | 11.58 | 17.43 | 27.70 | 42.36 | 64.59 | 97.63 | 146.55 | 218.39 | 299.44 | 385.99 | 0.384 |
| Hs-0 | 10.96 | 5.20 | 7.94 | 12.52 | 19.53 | 31.47 | 48.43 | 72.95 | 109.41 | 168.36 | 247.78 | 338.79 | 434.34 | 0.424 |
| HSm | 10.19 | 4.52 | 6.82 | 10.94 | 17.05 | 27.30 | 42.40 | 64.15 | 97.30 | 146.84 | 219.31 | 307.52 | 417.36 | 0.406 |
| In-0 | 12.25 | 6.12 | 8.99 | 13.08 | 20.34 | 32.77 | 51.44 | 78.57 | 118.04 | 178.71 | 272.40 | 384.14 | 494.94 | 0.364 |
| Is-1 | 12.81 | 5.20 | 7.90 | 12.36 | 19.51 | 32.33 | 50.17 | 75.91 | 115.27 | 175.46 | 265.55 | 368.18 | 480.96 | 0.421 |
| Je-0 | 9.43 | 5.76 | 8.09 | 12.01 | 18.48 | 30.47 | 47.56 | 72.96 | 108.07 | 158.06 | 225.49 | 301.37 | 376.14 | 0.350 |
| Je-54 | 8.25 | 4.67 | 6.75 | 10.38 | 15.86 | 25.81 | 40.50 | 61.70 | 89.73 | 131.66 | 195.09 | 279.55 | 382.27 | 0.382 |
| JEA | 6.95 | 6.39 | 9.20 | 13.73 | 19.47 | 29.70 | 43.57 | 60.46 | 82.38 | 113.54 | 161.94 | 222.44 | 287.62 | 0.358 |
| Jea | 7.45 | 6.41 | 9.36 | 13.98 | 20.34 | 31.48 | 47.31 | 67.59 | 91.54 | 126.52 | 178.44 | 245.03 | 317.02 | 0.368 |
| Jl-3 | 10.59 | 6.55 | 9.79 | 14.45 | 22.20 | 34.79 | 52.60 | 77.44 | 114.43 | 168.30 | 243.12 | 328.29 | 419.28 | 0.385 |
| Jm-1 | 10.02 | 5.28 | 7.83 | 12.93 | 19.47 | 30.01 | 45.38 | 69.27 | 104.08 | 157.38 | 234.30 | 325.09 | 425.42 | 0.408 |
| Kä-0 | 9.11 | 5.66 | 7.99 | 11.45 | 17.63 | 28.36 | 43.47 | 63.52 | 92.48 | 136.24 | 202.27 | 292.90 | 416.73 | 0.353 |
| Kas-2 | 13.22 | 7.80 | 11.57 | 17.52 | 26.63 | 41.61 | 63.01 | 94.21 | 138.59 | 194.14 | 268.59 | 355.79 | 444.40 | 0.393 |
| Kb-0 | 10.93 | 5.40 | 7.91 | 11.85 | 17.60 | 27.98 | 44.24 | 68.59 | 99.52 | 150.44 | 226.50 | 313.38 | 388.96 | 0.379 |

| accession | DW20 | PLA07 | PLA08 | PLA09 | PLA10 | PLA11 | PLA12 | PLA13 | PLA14 | PLA15 | PLA16 | PLA17 | PLA18 | RGR07_09 |
|---|---|---|---|---|---|---|---|---|---|---|---|---|---|---|
| KBS-Mac- | 12.68 | 7.94 | 11.39 | 16.93 | 24.91 | 38.33 | 56.97 | 84.62 | 126.82 | 190.79 | 280.02 | 379.95 | 480.99 | 0.366 |
| Kelsterbacl | 9.93 | 5.09 | 7.43 | 10.97 | 16.45 | 26.86 | 41.41 | 63.17 | 94.73 | 141.84 | 211.55 | 298.67 | 389.17 | 0.378 |
| Kelsterbacl | 15.54 | 6.50 | 9.87 | 15.72 | 24.41 | 40.48 | 63.32 | 96.88 | 149.61 | 229.61 | 345.93 | 460.57 | 575.68 | 0.420 |
| Kil-0 | 12.12 | 5.94 | 8.98 | 13.66 | 21.16 | 33.66 | 51.95 | 79.03 | 118.32 | 176.03 | 267.72 | 360.94 | 459.78 | 0.403 |
| Kin-0 | 10.36 | 5.77 | 8.63 | 13.04 | 19.23 | 30.06 | 45.98 | 68.37 | 103.28 | 160.12 | 240.14 | 336.87 | 439.71 | 0.396 |
| Kl-0 | 10.91 | 5.06 | 7.53 | 11.65 | 17.81 | 29.14 | 45.64 | 70.40 | 105.44 | 156.36 | 236.97 | 339.71 | 431.48 | 0.402 |
| Kl-5 | 10.59 | 5.13 | 8.03 | 12.63 | 18.24 | 29.42 | 46.12 | 71.06 | 102.48 | 150.76 | 223.09 | 303.38 | 390.55 | 0.420 |
| Kn-0 | 11.74 | 5.00 | 7.61 | 11.99 | 18.93 | 30.88 | 47.64 | 72.68 | 109.59 | 167.76 | 254.33 | 354.75 | 470.42 | 0.426 |
| KNO-11 | 13.01 | 6.88 | 9.68 | 13.66 | 22.65 | 37.35 | 56.84 | 85.93 | 127.86 | 190.84 | 278.13 | 364.67 | 450.00 | 0.330 |
| Kno-18 | 11.53 | 6.45 | 9.33 | 14.48 | 22.09 | 35.62 | 54.09 | 81.22 | 122.21 | 183.25 | 259.08 | 331.42 | 416.58 | 0.401 |
| Koln | 12.14 | 5.64 | 8.47 | 12.95 | 19.90 | 32.05 | 49.72 | 75.59 | 114.70 | 174.65 | 262.26 | 356.30 | 451.09 | 0.402 |
| Kondara | 12.94 | 6.84 | 10.53 | 16.53 | 24.13 | 37.84 | 57.17 | 84.69 | 122.56 | 186.01 | 269.87 | 359.69 | 451.58 | 0.416 |
| Kr-0 | 9.89 | 5.06 | 7.46 | 11.40 | 17.74 | 29.60 | 46.67 | 71.08 | 104.35 | 152.08 | 225.62 | 316.98 | 391.46 | 0.388 |
| Kro-0 | 10.87 | 5.00 | 6.67 | 9.87 | 16.34 | 27.48 | 43.60 | 66.20 | 100.17 | 151.47 | 229.62 | 333.65 | 439.41 | 0.329 |
| Krot-2 | 9.10 | 5.05 | 7.31 | 10.95 | 16.66 | 26.15 | 40.13 | 60.84 | 92.02 | 139.32 | 209.85 | 306.19 | 419.73 | 0.375 |
| Kz-1 | 11.05 | 6.44 | 9.89 | 15.43 | 24.23 | 38.08 | 56.62 | 81.24 | 120.16 | 174.91 | 272.43 | 383.22 | 490.92 | 0.430 |
| Kz-9 | 11.69 | 6.06 | 9.06 | 13.92 | 21.71 | 34.58 | 53.24 | 80.71 | 121.51 | 181.51 | 274.16 | 371.35 | 452.25 | 0.399 |
| LAC-3 | 11.44 | 6.36 | 9.18 | 13.65 | 20.81 | 33.34 | 51.72 | 78.84 | 115.65 | 173.94 | 255.25 | 341.26 | 418.13 | 0.370 |
| LAC-5 | 8.67 | 5.16 | 7.35 | 11.28 | 16.01 | 24.72 | 37.04 | 56.70 | 84.87 | 125.23 | 187.43 | 272.23 | 367.95 | 0.357 |
| Lan-0 | 11.36 | 7.15 | 10.41 | 15.50 | 24.16 | 38.54 | 59.11 | 89.78 | 134.72 | 201.61 | 296.40 | 401.29 | 512.99 | 0.374 |
| Laud-1 | 11.93 | 5.72 | 8.31 | 13.09 | 19.85 | 30.81 | 46.84 | 70.30 | 106.77 | 162.85 | 247.51 | 344.46 | 446.35 | 0.403 |
| Lc-0 | 8.99 | 4.76 | 6.86 | 10.56 | 15.31 | 24.40 | 37.90 | 57.99 | 88.62 | 133.69 | 199.33 | 280.92 | 374.05 | 0.374 |
| LDV-14 | 8.75 | 5.43 | 7.87 | 11.43 | 16.18 | 25.35 | 39.10 | 60.47 | 88.38 | 130.59 | 197.39 | 283.70 | 393.12 | 0.366 |
| LDV-25 | 11.20 | 6.82 | 9.78 | 14.46 | 21.64 | 33.40 | 50.88 | 76.69 | 115.22 | 172.96 | 253.91 | 343.53 | 428.22 | 0.368 |
| LDV-34 | 11.10 | 6.44 | 9.27 | 13.77 | 20.76 | 32.89 | 50.16 | 75.49 | 112.82 | 167.15 | 243.80 | 336.68 | 442.74 | 0.376 |
| LDV-58 | 10.23 | 5.88 | 8.80 | 13.04 | 19.31 | 29.73 | 45.23 | 67.96 | 102.26 | 154.12 | 228.27 | 313.02 | 409.32 | 0.385 |
| Ler-1 | 11.22 | 5.85 | 8.44 | 12.73 | 19.39 | 30.83 | 47.56 | 71.36 | 104.97 | 149.87 | 219.64 | 308.71 | 421.16 | 0.383 |
| Li-3 | 10.68 | 5.19 | 7.62 | 11.88 | 18.13 | 29.48 | 45.55 | 69.34 | 103.34 | 153.74 | 227.51 | 307.80 | 380.15 | 0.390 |
| Li-5:2 | 9.05 | 8.16 | 11.59 | 16.54 | 23.96 | 37.11 | 56.30 | 82.06 | 116.55 | 158.68 | 216.35 | 295.79 | 372.51 | 0.341 |
| Li-6 | 13.50 | 7.27 | 10.66 | 15.57 | 24.33 | 39.08 | 60.20 | 91.25 | 135.09 | 204.86 | 308.68 | 426.68 | 547.72 | 0.376 |

| accession | DW20 | PLA07 | PLA08 | PLA09 | PLA10 | PLA11 | PLA12 | PLA13 | PLA14 | PLA15 | PLA16 | PLA17 | PLA18 | RGR07_09 |
|---|---|---|---|---|---|---|---|---|---|---|---|---|---|---|
| Li-7 | 10.36 | 5.43 | 7.88 | 11.94 | 17.96 | 28.02 | 42.85 | 65.48 | 97.92 | 141.73 | 205.35 | 290.58 | 375.80 | 0.376 |
| Liarum | 10.51 | 5.83 | 8.48 | 12.98 | 19.69 | 31.49 | 48.47 | 73.00 | 109.24 | 160.10 | 236.43 | 319.81 | 411.60 | 0.390 |
| Limeport | 10.69 | 5.71 | 8.44 | 13.08 | 19.96 | 32.37 | 49.13 | 74.08 | 110.59 | 165.41 | 247.41 | 343.49 | 450.12 | 0.392 |
| Lip-0 | 14.53 | 6.25 | 9.62 | 15.36 | 23.96 | 38.09 | 58.46 | 87.51 | 133.06 | 200.06 | 293.12 | 389.21 | 493.39 | 0.433 |
| Lis-1 | 11.70 | 5.85 | 8.26 | 13.04 | 20.18 | 32.75 | 51.37 | 79.12 | 118.68 | 179.13 | 267.53 | 369.52 | 492.01 | 0.388 |
| LL-0 | 4.45 | 4.43 | 6.46 | 9.46 | 13.22 | 18.91 | 26.26 | 35.22 | 46.40 | 64.16 | 90.67 | 124.09 | 178.57 | 0.346 |
| Ll-OF-095 | 10.05 | 5.67 | 8.21 | 12.24 | 18.09 | 27.48 | 41.24 | 62.86 | 91.64 | 134.29 | 198.41 | 283.63 | 362.22 | 0.379 |
| Lm | 11.76 | 5.95 | 8.41 | 12.15 | 18.71 | 30.47 | 48.77 | 73.83 | 112.65 | 175.78 | 254.82 | 353.51 | 462.94 | 0.346 |
| Lm-2 | 11.68 | 5.99 | 8.40 | 12.09 | 18.60 | 30.90 | 49.18 | 74.57 | 112.47 | 176.39 | 262.75 | 364.58 | 482.14 | 0.343 |
| Lom1-1 | 13.13 | 7.79 | 11.93 | 18.05 | 26.32 | 42.03 | 64.72 | 97.33 | 143.55 | 212.05 | 298.01 | 384.31 | 462.44 | 0.397 |
| Löv-5 | 11.40 | 7.94 | 10.96 | 15.40 | 24.36 | 39.41 | 61.21 | 90.55 | 131.02 | 188.72 | 269.04 | 365.10 | 467.99 | 0.320 |
| Lp2-2 | 15.81 | 6.87 | 10.64 | 16.36 | 25.39 | 40.67 | 62.37 | 93.46 | 143.89 | 223.55 | 324.19 | 427.45 | 547.24 | 0.431 |
| Lp2-6 | 11.24 | 4.94 | 7.47 | 11.72 | 18.29 | 29.83 | 46.80 | 71.38 | 108.84 | 166.62 | 253.64 | 354.99 | 457.38 | 0.416 |
| Lu | 9.10 | 5.22 | 7.13 | 10.92 | 16.62 | 26.27 | 40.13 | 60.23 | 89.33 | 129.42 | 193.70 | 280.25 | 382.78 | 0.359 |
| Lz-0 | 14.34 | 7.67 | 11.35 | 17.20 | 26.74 | 43.82 | 67.43 | 102.17 | 150.40 | 220.70 | 303.90 | 384.70 | 448.24 | 0.397 |
| Map-42 | 11.18 | 7.02 | 10.16 | 15.28 | 22.91 | 35.65 | 54.30 | 81.15 | 119.66 | 176.01 | 256.60 | 351.86 | 445.84 | 0.374 |
| Mc-0 | 9.63 | 7.59 | 10.53 | 15.23 | 21.25 | 32.13 | 47.01 | 67.99 | 100.38 | 147.88 | 220.94 | 332.15 | 440.33 | 0.329 |
| Me-0 | 8.67 | 4.98 | 7.61 | 11.69 | 17.18 | 27.48 | 41.92 | 61.59 | 91.45 | 138.87 | 204.53 | 283.88 | 374.58 | 0.408 |
| Mh-0 | 8.69 | 6.16 | 8.77 | 13.41 | 19.93 | 30.75 | 45.37 | 64.69 | 95.83 | 141.24 | 204.83 | 285.49 | 376.87 | 0.372 |
| Mh-1 | 11.00 | 5.69 | 8.52 | 12.90 | 19.49 | 30.49 | 45.99 | 69.62 | 105.42 | 155.12 | 227.51 | 305.01 | 379.31 | 0.387 |
| MIB-15 | 10.04 | 6.14 | 8.82 | 12.74 | 19.79 | 31.64 | 47.63 | 71.09 | 105.24 | 155.01 | 227.27 | 316.02 | 401.19 | 0.352 |
| MIB-22 | 9.90 | 5.38 | 7.68 | 11.61 | 17.00 | 29.03 | 45.55 | 69.09 | 104.52 | 156.65 | 232.45 | 323.23 | 423.33 | 0.363 |
| MIB-28 | 10.57 | 5.57 | 8.13 | 12.29 | 19.16 | 30.41 | 46.85 | 71.49 | 107.49 | 162.08 | 240.58 | 330.68 | 445.80 | 0.381 |
| MIB-84 | 8.66 | 4.37 | 6.31 | 9.51 | 14.52 | 23.64 | 36.95 | 55.87 | 84.28 | 125.77 | 190.09 | 274.11 | 370.95 | 0.376 |
| MNF-Pot-4 | 8.05 | 4.48 | 6.35 | 10.01 | 14.70 | 23.70 | 36.78 | 56.15 | 82.90 | 126.99 | 193.11 | 277.07 | 374.63 | 0.380 |
| Mnz-0 | 12.26 | 5.96 | 9.16 | 13.94 | 20.86 | 33.06 | 50.75 | 75.62 | 111.12 | 167.35 | 248.27 | 345.21 | 439.58 | 0.415 |
| MOG-37 | 7.82 | 4.24 | 6.02 | 8.95 | 12.81 | 21.02 | 32.39 | 49.75 | 74.75 | 112.32 | 170.87 | 246.14 | 331.92 | 0.367 |
| Mrk-0 | 10.96 | 6.10 | 9.14 | 14.00 | 20.97 | 33.14 | 50.56 | 75.97 | 112.55 | 169.99 | 251.73 | 352.27 | 461.63 | 0.400 |
| Ms-0 | 13.50 | 6.09 | 9.11 | 14.16 | 21.51 | 36.01 | 59.05 | 93.39 | 138.55 | 207.59 | 300.52 | 405.54 | 513.69 | 0.401 |
| Mt-0 | 13.49 | 6.11 | 9.22 | 14.46 | 22.68 | 36.37 | 56.49 | 85.90 | 130.13 | 200.02 | 298.74 | 405.99 | 519.84 | 0.419 |

| accession | DW20 | PLA07 | PLA08 | PLA09 | PLA10 | PLA11 | PLA12 | PLA13 | PLA14 | PLA15 | PLA16 | PLA17 | PLA18 | RGR07_09 |
|---|---|---|---|---|---|---|---|---|---|---|---|---|---|---|
| Mz-0 | 9.23 | 6.50 | 9.62 | 14.71 | 22.18 | 34.47 | 51.09 | 73.70 | 103.49 | 147.00 | 207.24 | 279.29 | 360.84 | 0.392 |
| N13 | 10.40 | 4.83 | 6.90 | 10.51 | 16.39 | 27.53 | 44.47 | 70.76 | 107.12 | 160.87 | 243.78 | 345.78 | 469.66 | 0.384 |
| N4 | 15.64 | 9.01 | 13.01 | 19.80 | 30.49 | 49.15 | 77.15 | 118.65 | 173.94 | 251.48 | 364.05 | 489.78 | 604.10 | 0.387 |
| N7 | 13.12 | 6.59 | 9.93 | 15.10 | 22.39 | 35.59 | 55.42 | 86.34 | 128.58 | 190.05 | 275.94 | 370.42 | 454.14 | 0.415 |
| Na-1 | 9.23 | 4.24 | 6.16 | 9.15 | 13.49 | 23.39 | 36.94 | 56.43 | 85.08 | 127.20 | 189.50 | 271.91 | 364.61 | 0.359 |
| Nc-1 | 10.08 | 5.33 | 7.65 | 11.25 | 17.13 | 28.62 | 44.47 | 66.26 | 97.49 | 144.12 | 209.17 | 289.70 | 362.79 | 0.352 |
| Nd | 7.86 | 4.23 | 5.89 | 9.21 | 13.62 | 21.50 | 33.61 | 50.88 | 74.42 | 109.01 | 168.13 | 249.62 | 337.15 | 0.364 |
| Nd-1 | 8.96 | 4.56 | 6.52 | 9.66 | 15.01 | 23.92 | 37.15 | 55.67 | 82.54 | 123.74 | 188.59 | 265.94 | 357.63 | 0.358 |
| NFA-10 | 10.77 | 6.55 | 9.66 | 14.46 | 21.70 | 33.39 | 50.82 | 75.31 | 113.16 | 168.97 | 243.78 | 331.34 | 415.05 | 0.381 |
| NFA-8 | 14.85 | 8.43 | 12.17 | 18.19 | 29.17 | 46.70 | 71.64 | 107.30 | 160.07 | 239.45 | 330.07 | 409.28 | 478.35 | 0.373 |
| NFC-20 | 11.72 | 8.07 | 11.70 | 17.43 | 26.47 | 40.25 | 61.33 | 91.33 | 135.51 | 199.95 | 292.83 | 392.26 | 496.72 | 0.373 |
| No-0 | 13.95 | 7.50 | 10.72 | 16.33 | 25.15 | 41.55 | 64.54 | 99.76 | 147.09 | 216.44 | 301.63 | 387.66 | 458.22 | 0.370 |
| Nok-1 | 10.31 | 5.80 | 8.47 | 12.98 | 19.52 | 31.06 | 45.74 | 68.63 | 101.31 | 144.43 | 206.89 | 285.55 | 360.25 | 0.388 |
| Nok-2 | 12.86 | 6.41 | 9.44 | 14.42 | 22.09 | 35.44 | 54.96 | 85.02 | 127.16 | 188.23 | 282.98 | 387.01 | 486.30 | 0.396 |
| Nw-0 | 7.96 | 3.71 | 5.63 | 8.97 | 13.51 | 22.05 | 33.80 | 50.69 | 76.00 | 115.48 | 168.44 | 240.98 | 328.64 | 0.418 |
| Nw-2 | 11.27 | 4.80 | 6.91 | 11.04 | 16.48 | 26.73 | 41.92 | 65.67 | 101.13 | 153.31 | 229.74 | 318.28 | 406.77 | 0.391 |
| NW3 | 13.35 | 7.14 | 10.48 | 15.40 | 24.16 | 38.09 | 58.57 | 88.36 | 131.56 | 197.13 | 281.39 | 375.16 | 457.11 | 0.371 |
| Nz1 | 11.72 | 5.78 | 8.55 | 12.91 | 19.96 | 32.10 | 48.56 | 74.86 | 110.30 | 162.98 | 232.00 | 311.23 | 393.60 | 0.385 |
| Ob-1 | 9.17 | 4.37 | 6.48 | 10.43 | 15.65 | 25.15 | 38.41 | 58.32 | 87.78 | 134.60 | 206.09 | 300.66 | 386.96 | 0.406 |
| Old-1 | 11.75 | 5.90 | 8.92 | 14.33 | 21.01 | 34.10 | 53.01 | 81.73 | 122.51 | 179.90 | 260.66 | 363.86 | 461.47 | 0.418 |
| Or-0 | 9.80 | 5.70 | 8.46 | 12.83 | 18.60 | 29.16 | 43.95 | 65.15 | 97.95 | 150.76 | 230.70 | 330.14 | 448.65 | 0.387 |
| Ors-1 | 12.82 | 8.09 | 11.24 | 16.32 | 24.64 | 39.86 | 61.69 | 92.91 | 137.53 | 203.97 | 300.48 | 415.40 | 522.72 | 0.343 |
| Ors-2 | 11.49 | 7.53 | 11.10 | 16.18 | 24.07 | 38.45 | 58.71 | 88.13 | 127.41 | 186.79 | 274.76 | 390.71 | 503.39 | 0.373 |
| Ost-0 | 9.59 | 7.87 | 11.17 | 15.14 | 22.83 | 35.73 | 54.20 | 78.46 | 110.28 | 152.28 | 214.68 | 293.91 | 381.81 | 0.319 |
| Ove-0 | 13.36 | 8.06 | 11.61 | 17.37 | 26.28 | 41.17 | 62.70 | 93.58 | 136.47 | 198.99 | 281.05 | 371.71 | 467.21 | 0.376 |
| Ove-0 | 13.00 | 7.78 | 11.19 | 16.67 | 24.76 | 38.80 | 58.74 | 87.70 | 127.53 | 188.55 | 277.36 | 381.69 | 482.75 | 0.370 |
| Oy-0 | 13.18 | 6.28 | 9.65 | 15.14 | 22.67 | 36.18 | 57.22 | 89.20 | 136.39 | 209.23 | 310.36 | 421.43 | 552.56 | 0.414 |
| Oy-1 | 10.32 | 5.22 | 7.88 | 12.09 | 18.15 | 28.55 | 43.73 | 66.98 | 98.86 | 152.94 | 230.23 | 325.90 | 416.49 | 0.402 |
| Pa-1 | 10.57 | 5.67 | 8.62 | 13.04 | 19.35 | 30.64 | 46.72 | 70.26 | 105.72 | 161.23 | 233.63 | 316.45 | 408.10 | 0.394 |
| Pa-2 | 9.33 | 5.88 | 8.88 | 14.00 | 22.07 | 35.06 | 52.57 | 76.75 | 114.19 | 170.96 | 246.43 | 329.57 | 401.83 | 0.422 |

| accession | DW20 | PLA07 | PLA08 | PLA09 | PLA10 | PLA11 | PLA12 | PLA13 | PLA14 | PLA15 | PLA16 | PLA17 | PLA18 | RGR07_09 |
|---|---|---|---|---|---|---|---|---|---|---|---|---|---|---|
| PAR-3 | 9.76 | 5.93 | 8.55 | 12.98 | 19.25 | 29.23 | 42.33 | 61.05 | 89.06 | 130.10 | 188.55 | 261.40 | 330.77 | 0.379 |
| PAR-4 | 10.03 | 5.80 | 8.17 | 11.76 | 18.02 | 28.11 | 42.21 | 63.06 | 92.21 | 137.57 | 210.42 | 298.58 | 401.07 | 0.346 |
| Per-1 | 12.26 | 5.83 | 8.55 | 13.05 | 19.82 | 31.98 | 49.27 | 74.97 | 110.71 | 166.50 | 256.21 | 373.36 | 497.39 | 0.388 |
| Petergof | 8.80 | 4.99 | 7.07 | 10.05 | 15.25 | 25.00 | 37.32 | 55.26 | 82.28 | 123.46 | 185.44 | 268.72 | 364.03 | 0.335 |
| PHW-10 | 8.84 | 6.36 | 9.34 | 13.85 | 20.39 | 32.02 | 49.43 | 74.65 | 109.98 | 160.82 | 224.31 | 300.79 | 363.22 | 0.375 |
| PHW-13 | 8.79 | 5.29 | 7.65 | 11.36 | 16.61 | 25.67 | 38.24 | 56.79 | 83.61 | 126.03 | 193.78 | 278.35 | 371.90 | 0.371 |
| PHW-14 | 9.23 | 4.93 | 7.24 | 10.99 | 16.34 | 25.80 | 38.98 | 58.29 | 87.13 | 128.10 | 187.99 | 266.98 | 359.06 | 0.393 |
| PHW-20 | 6.84 | 4.74 | 6.88 | 10.38 | 14.98 | 22.99 | 34.33 | 50.76 | 73.76 | 107.65 | 157.62 | 217.93 | 275.82 | 0.369 |
| PHW-22 | 8.67 | 5.42 | 7.95 | 12.54 | 18.38 | 27.64 | 40.16 | 60.62 | 91.97 | 139.86 | 206.47 | 289.80 | 372.38 | 0.382 |
| PHW-26 | 11.47 | 6.65 | 9.60 | 14.33 | 22.24 | 35.99 | 57.00 | 87.12 | 127.49 | 185.44 | 266.90 | 342.45 | 406.27 | 0.369 |
| PHW-28 | 10.96 | 6.24 | 9.09 | 14.15 | 21.06 | 32.31 | 48.93 | 73.20 | 106.75 | 156.21 | 230.33 | 316.12 | 379.01 | 0.388 |
| PHW-31 | 6.63 | 5.40 | 7.64 | 11.11 | 15.91 | 24.57 | 37.01 | 54.92 | 79.58 | 115.28 | 172.54 | 257.12 | 363.58 | 0.338 |
| PHW-33 | 9.64 | 6.76 | 9.81 | 14.67 | 22.26 | 34.72 | 52.38 | 77.69 | 112.09 | 162.94 | 223.16 | 291.50 | 374.58 | 0.376 |
| PHW-35 | 7.83 | 6.77 | 9.85 | 14.40 | 22.44 | 35.27 | 51.71 | 72.45 | 97.76 | 137.00 | 185.57 | 246.69 | 330.48 | 0.367 |
| PHW-36 | 12.29 | 6.53 | 9.38 | 14.32 | 21.61 | 34.21 | 52.54 | 80.01 | 119.84 | 180.43 | 264.93 | 350.30 | 432.85 | 0.377 |
| PHW-37 | 9.79 | 5.69 | 8.09 | 12.36 | 18.93 | 30.40 | 46.36 | 70.45 | 100.84 | 151.23 | 225.26 | 315.36 | 408.66 | 0.374 |
| Pi-0 | 10.02 | 6.04 | 8.97 | 13.48 | 19.23 | 29.17 | 43.99 | 65.42 | 95.13 | 139.81 | 209.88 | 291.83 | 378.23 | 0.380 |
| Pla-0 | 11.49 | 6.84 | 10.14 | 15.25 | 22.66 | 35.85 | 54.17 | 81.34 | 119.97 | 177.40 | 256.71 | 331.46 | 406.85 | 0.385 |
| Pn-0 | 10.52 | 6.27 | 9.00 | 13.63 | 20.49 | 31.18 | 47.27 | 71.69 | 106.88 | 158.37 | 231.48 | 313.51 | 401.40 | 0.377 |
| Pna-10 | 10.69 | 6.79 | 9.82 | 14.56 | 21.26 | 32.34 | 48.71 | 72.74 | 106.33 | 157.92 | 231.54 | 320.56 | 411.41 | 0.364 |
| Pna-17 | 9.29 | 5.28 | 7.45 | 9.95 | 16.23 | 28.04 | 42.66 | 63.36 | 92.25 | 136.52 | 204.74 | 286.49 | 366.76 | 0.304 |
| Pna-17 | 9.13 | 5.27 | 7.50 | 9.41 | 16.96 | 27.72 | 41.82 | 62.05 | 89.61 | 130.26 | 195.73 | 274.69 | 359.13 | 0.270 |
| Po-0 | 13.36 | 6.95 | 10.13 | 15.37 | 23.30 | 37.09 | 57.09 | 86.16 | 127.02 | 191.11 | 284.33 | 383.79 | 487.65 | 0.385 |
| Pog-0 | 10.39 | 6.19 | 8.97 | 12.84 | 20.19 | 31.01 | 46.14 | 67.32 | 101.03 | 154.57 | 227.39 | 307.80 | 399.04 | 0.356 |
| Pr-0 | 8.78 | 4.63 | 6.82 | 10.71 | 16.98 | 27.82 | 42.51 | 64.32 | 97.52 | 149.50 | 214.65 | 286.25 | 364.51 | 0.418 |
| Pro-0 | 9.46 | 6.19 | 8.89 | 13.52 | 20.37 | 31.70 | 46.63 | 68.60 | 99.40 | 147.84 | 225.27 | 316.92 | 409.69 | 0.379 |
| Pt-0 | 11.19 | 5.91 | 8.55 | 12.75 | 19.46 | 31.81 | 49.07 | 75.01 | 114.73 | 176.73 | 267.21 | 370.82 | 484.32 | 0.371 |
| Pu2-23 | 8.69 | 4.91 | 7.36 | 11.03 | 17.74 | 29.44 | 45.76 | 68.27 | 103.32 | 155.58 | 219.31 | 291.22 | 379.34 | 0.386 |
| PU2-24 | 14.68 | 6.62 | 9.93 | 15.17 | 23.87 | 39.81 | 64.71 | 100.60 | 154.62 | 230.58 | 324.17 | 413.71 | 498.56 | 0.396 |
| Pu2-7 | 11.16 | 6.14 | 9.02 | 13.51 | 20.73 | 33.57 | 52.16 | 79.64 | 118.70 | 181.86 | 272.78 | 380.15 | 483.32 | 0.386 |

| accession | DW20 | PLA07 | PLA08 | PLA09 | PLA10 | PLA11 | PLA12 | PLA13 | PLA14 | PLA15 | PLA16 | PLA17 | PLA18 | RGR07_09 |
|---|---|---|---|---|---|---|---|---|---|---|---|---|---|---|
| Pyl-1 | 8.02 | 6.29 | 8.77 | 13.34 | 18.50 | 28.57 | 42.65 | 63.29 | 91.12 | 127.26 | 183.19 | 250.70 | 318.53 | 0.343 |
| Ra-0 | 9.68 | 5.22 | 7.83 | 11.94 | 17.54 | 27.46 | 41.82 | 63.31 | 94.82 | 142.17 | 213.19 | 302.30 | 385.95 | 0.394 |
| Rak-2 | 11.88 | 7.13 | 10.04 | 14.57 | 22.49 | 35.49 | 54.36 | 81.61 | 120.67 | 179.86 | 262.95 | 367.32 | 482.38 | 0.344 |
| Ren-1 | 9.09 | 5.01 | 7.46 | 11.27 | 16.97 | 26.60 | 40.15 | 60.62 | 91.22 | 136.76 | 199.18 | 278.42 | 366.74 | 0.390 |
| Ren-11 | 9.25 | 5.40 | 7.51 | 10.86 | 16.09 | 25.75 | 39.23 | 58.19 | 84.97 | 132.03 | 204.02 | 306.87 | 446.45 | 0.345 |
| Rhen-1 | 9.82 | 5.80 | 8.25 | 11.58 | 18.00 | 28.54 | 43.08 | 63.75 | 93.29 | 140.19 | 206.49 | 283.65 | 374.97 | 0.344 |
| Ri-0 | 11.19 | 6.41 | 9.71 | 14.55 | 21.76 | 33.84 | 51.36 | 76.03 | 116.26 | 176.59 | 254.08 | 342.54 | 448.06 | 0.394 |
| RLD-1 | 14.95 | 6.58 | 9.69 | 14.92 | 23.41 | 39.15 | 62.23 | 94.45 | 134.97 | 207.77 | 321.34 | 454.77 | 571.47 | 0.406 |
| RLD-2 | 16.11 | 6.96 | 10.16 | 15.85 | 25.42 | 42.25 | 67.09 | 101.89 | 148.20 | 223.06 | 336.93 | 468.69 | 602.13 | 0.401 |
| Rmx-A02 | 12.76 | 6.76 | 9.77 | 14.99 | 23.22 | 36.37 | 56.22 | 85.80 | 127.46 | 195.46 | 285.16 | 378.43 | 478.31 | 0.389 |
| Rmx-A180 | 13.00 | 6.22 | 9.16 | 13.74 | 21.53 | 34.80 | 54.32 | 83.31 | 126.29 | 193.76 | 288.08 | 398.11 | 495.57 | 0.383 |
| Rou-0 | 9.66 | 6.24 | 9.12 | 13.32 | 20.25 | 31.69 | 47.44 | 70.05 | 102.71 | 152.29 | 221.36 | 298.57 | 374.77 | 0.373 |
| RRS-10 | 9.28 | 5.39 | 7.72 | 11.37 | 16.73 | 26.86 | 40.95 | 61.51 | 90.28 | 134.31 | 201.98 | 285.74 | 361.25 | 0.354 |
| RRS-7 | 9.60 | 5.60 | 8.01 | 11.79 | 17.12 | 26.80 | 40.79 | 61.46 | 90.40 | 133.41 | 194.56 | 269.52 | 353.05 | 0.361 |
| Rsch-0 | 12.47 | 6.27 | 9.05 | 13.95 | 21.64 | 35.74 | 55.43 | 82.76 | 118.12 | 176.85 | 268.94 | 383.36 | 510.62 | 0.389 |
| Rsch-4 | 12.67 | 6.03 | 9.10 | 14.14 | 22.30 | 36.45 | 57.01 | 87.20 | 131.54 | 200.55 | 295.97 | 393.40 | 471.05 | 0.409 |
| Rubeszhno | 11.88 | 6.05 | 8.88 | 13.61 | 20.58 | 33.29 | 52.37 | 80.12 | 115.33 | 174.77 | 260.29 | 352.05 | 439.19 | 0.385 |
| S96 | 9.74 | 4.93 | 7.04 | 11.13 | 16.52 | 26.60 | 40.44 | 61.49 | 92.71 | 136.91 | 203.01 | 283.19 | 384.79 | 0.378 |
| Santa Clara | 6.60 | 4.23 | 6.19 | 9.11 | 13.44 | 20.03 | 30.73 | 47.38 | 71.75 | 101.15 | 138.16 | 196.98 | 288.66 | 0.368 |
| Sap-0 | 10.65 | 4.74 | 7.23 | 11.12 | 16.88 | 27.58 | 43.12 | 66.67 | 102.42 | 154.20 | 231.06 | 322.49 | 407.93 | 0.404 |
| Sapporo-0 | 10.15 | 5.78 | 8.43 | 12.85 | 19.28 | 30.75 | 46.96 | 70.27 | 100.64 | 154.22 | 227.47 | 308.35 | 396.92 | 0.380 |
| Sav-0 | 11.07 | 6.59 | 9.39 | 14.36 | 20.99 | 32.20 | 49.15 | 74.50 | 111.18 | 164.96 | 238.55 | 328.00 | 402.86 | 0.362 |
| Sav-0 | 11.91 | 5.22 | 8.15 | 13.40 | 19.54 | 30.91 | 47.55 | 72.60 | 108.26 | 158.39 | 227.84 | 319.72 | 411.54 | 0.416 |
| Se-0 | 11.82 | 6.79 | 10.30 | 15.24 | 22.87 | 37.01 | 56.12 | 83.73 | 121.50 | 180.96 | 260.24 | 346.82 | 430.22 | 0.392 |
| Sei-0 | 10.19 | 4.98 | 7.07 | 11.24 | 16.10 | 25.97 | 40.25 | 61.81 | 93.83 | 143.12 | 217.59 | 307.86 | 406.95 | 0.378 |
| Sg-1 | 9.67 | 4.89 | 7.32 | 11.36 | 17.14 | 27.17 | 40.81 | 61.36 | 90.92 | 134.62 | 198.68 | 275.97 | 351.37 | 0.401 |
| Sh-0 | 11.03 | 5.72 | 8.43 | 13.36 | 20.28 | 32.66 | 49.68 | 75.07 | 110.53 | 168.15 | 253.11 | 356.60 | 463.11 | 0.399 |
| Shahdara | 10.90 | 6.50 | 9.53 | 14.10 | 20.74 | 32.12 | 48.95 | 73.37 | 106.77 | 156.03 | 234.48 | 330.36 | 429.67 | 0.375 |
| Si-0 | 9.04 | 4.36 | 6.33 | 9.93 | 15.35 | 25.13 | 39.40 | 60.55 | 92.37 | 138.59 | 206.90 | 284.99 | 386.60 | 0.385 |
| SLSP-30 | 9.58 | 5.00 | 7.51 | 11.54 | 17.31 | 27.54 | 42.01 | 63.30 | 93.66 | 141.63 | 208.44 | 283.91 | 351.39 | 0.408 |

| accession | DW20 | PLA07 | PLA08 | PLA09 | PLA10 | PLA11 | PLA12 | PLA13 | PLA14 | PLA15 | PLA16 | PLA17 | PLA18 | RGR07_09 |
|---|---|---|---|---|---|---|---|---|---|---|---|---|---|---|
| Sorbo | 14.49 | 8.27 | 12.16 | 17.81 | 26.73 | 42.44 | 64.84 | 96.52 | 141.09 | 206.55 | 291.39 | 379.20 | 457.14 | 0.365 |
| Sp-0 | 12.13 | 5.27 | 7.78 | 11.91 | 18.77 | 31.39 | 50.72 | 79.44 | 121.21 | 184.95 | 292.14 | 416.67 | 554.98 | 0.400 |
| Sq-1 | 9.75 | 5.31 | 7.83 | 11.49 | 17.40 | 28.54 | 43.92 | 66.37 | 97.82 | 147.49 | 223.12 | 310.44 | 394.10 | 0.378 |
| Sq-8 | 12.85 | 6.62 | 9.75 | 14.72 | 22.70 | 36.73 | 56.47 | 86.24 | 130.19 | 196.39 | 289.49 | 396.62 | 494.78 | 0.387 |
| St-0 | 11.92 | 5.46 | 8.01 | 12.45 | 19.30 | 31.23 | 47.68 | 73.36 | 110.26 | 164.95 | 253.78 | 352.18 | 439.22 | 0.398 |
| Ste-0 | 11.97 | 5.87 | 8.62 | 12.69 | 20.82 | 34.65 | 54.05 | 83.05 | 124.01 | 184.27 | 270.85 | 365.03 | 462.81 | 0.368 |
| Ste-3 | 6.64 | 3.42 | 4.79 | 7.89 | 11.06 | 18.32 | 28.96 | 44.88 | 66.64 | 97.86 | 147.32 | 215.11 | 297.93 | 0.325 |
| Stw-0 | 12.29 | 5.92 | 8.88 | 13.56 | 20.71 | 33.08 | 50.89 | 76.53 | 115.52 | 172.50 | 252.61 | 341.01 | 427.85 | 0.403 |
| Ta-0 | 13.85 | 7.32 | 11.01 | 16.53 | 25.19 | 38.34 | 58.92 | 88.30 | 131.90 | 187.95 | 269.90 | 361.61 | 454.71 | 0.393 |
| TAMM-2 | 9.10 | 6.44 | 9.07 | 12.29 | 17.57 | 28.56 | 44.36 | 68.27 | 98.74 | 145.04 | 223.04 | 309.11 | 390.76 | 0.317 |
| Tamm-27 | 10.92 | 7.08 | 10.06 | 12.67 | 19.51 | 31.25 | 47.86 | 71.08 | 103.24 | 157.11 | 230.47 | 310.16 | 378.95 | 0.287 |
| TDr-1 | 11.90 | 5.79 | 8.56 | 12.96 | 20.75 | 34.43 | 54.40 | 84.46 | 126.14 | 190.16 | 276.13 | 366.96 | 475.55 | 0.388 |
| TDr-3 | 9.27 | 5.30 | 7.71 | 11.81 | 18.09 | 29.39 | 45.47 | 69.67 | 104.85 | 163.85 | 247.35 | 345.55 | 450.11 | 0.379 |
| Te-0 | 12.07 | 7.36 | 10.40 | 15.36 | 24.43 | 40.27 | 63.58 | 96.93 | 143.17 | 208.77 | 306.34 | 402.90 | 503.80 | 0.356 |
| Tha-1 | 9.94 | 4.95 | 7.33 | 12.03 | 18.15 | 28.00 | 42.16 | 63.44 | 94.91 | 140.83 | 215.31 | 308.12 | 392.75 | 0.397 |
| Ting-1 | 11.37 | 6.93 | 9.97 | 14.33 | 22.89 | 36.26 | 55.36 | 82.29 | 121.68 | 178.23 | 258.24 | 355.74 | 452.01 | 0.350 |
| Tiv-1 | 10.50 | 6.34 | 9.30 | 13.80 | 20.77 | 33.33 | 51.23 | 77.31 | 114.86 | 167.66 | 243.99 | 333.23 | 431.12 | 0.393 |
| Tol-0 | 12.47 | 6.98 | 10.01 | 14.31 | 21.86 | 34.32 | 51.71 | 77.81 | 113.24 | 164.92 | 240.72 | 324.10 | 398.34 | 0.351 |
| Tottarp-2 | 12.70 | 5.72 | 8.55 | 13.69 | 20.91 | 33.54 | 51.83 | 78.52 | 117.38 | 178.32 | 268.50 | 378.75 | 490.93 | 0.414 |
| TOU-A1-1 | 12.45 | 6.91 | 9.67 | 14.31 | 21.62 | 34.14 | 52.93 | 79.95 | 119.10 | 177.08 | 259.05 | 362.53 | 481.48 | 0.358 |
| TOU-A1-1 | 10.82 | 5.76 | 8.53 | 13.11 | 19.70 | 30.89 | 46.76 | 70.98 | 104.83 | 157.09 | 231.22 | 332.51 | 454.80 | 0.397 |
| TOU-A1-4 | 16.09 | 8.34 | 12.11 | 18.70 | 28.69 | 45.68 | 70.12 | 105.62 | 156.33 | 232.04 | 335.77 | 444.64 | 558.29 | 0.396 |
| TOU-A1-6 | 11.22 | 6.55 | 9.57 | 14.41 | 21.27 | 33.17 | 49.86 | 73.93 | 107.85 | 161.61 | 240.45 | 334.70 | 435.41 | 0.376 |
| TOU-A1-9 | 9.90 | 5.80 | 8.51 | 12.78 | 19.38 | 30.63 | 46.09 | 67.76 | 97.68 | 143.29 | 211.82 | 300.01 | 394.55 | 0.386 |
| TOU-C-3 | 11.90 | 7.22 | 10.79 | 15.71 | 23.33 | 35.39 | 53.19 | 78.95 | 115.21 | 171.30 | 248.63 | 341.72 | 439.76 | 0.387 |
| TOU-E-11 | 11.08 | 6.17 | 9.11 | 13.89 | 21.43 | 34.95 | 54.47 | 83.61 | 121.31 | 180.14 | 257.17 | 345.21 | 422.34 | 0.398 |
| TOU-H-13 | 10.83 | 5.87 | 8.58 | 12.87 | 19.19 | 30.41 | 46.25 | 70.55 | 106.02 | 159.26 | 242.14 | 338.35 | 437.84 | 0.384 |
| TOU-I-17 | 8.37 | 5.10 | 7.69 | 12.14 | 18.32 | 28.69 | 43.57 | 65.48 | 97.29 | 145.16 | 214.74 | 300.34 | 387.05 | 0.410 |
| TOU-I-2 | 12.14 | 8.55 | 12.61 | 19.00 | 29.16 | 45.75 | 68.11 | 98.76 | 139.97 | 203.14 | 286.29 | 390.45 | 502.29 | 0.392 |
| TOU-I-6 | 16.47 | 7.20 | 11.33 | 17.93 | 29.01 | 47.13 | 73.72 | 112.85 | 168.95 | 253.45 | 363.16 | 475.25 | 575.76 | 0.446 |

| accession | DW20 | PLA07 | PLA08 | PLA09 | PLA10 | PLA11 | PLA12 | PLA13 | PLA14 | PLA15 | PLA16 | PLA17 | PLA18 | RGR07_09 |
|---|---|---|---|---|---|---|---|---|---|---|---|---|---|---|
| Ts-1 | 10.70 | 6.02 | 8.71 | 12.98 | 19.87 | 31.44 | 47.31 | 70.12 | 102.00 | 147.78 | 219.48 | 313.44 | 403.17 | 0.374 |
| Ts-5 | 10.85 | 5.78 | 8.44 | 11.63 | 17.55 | 28.80 | 43.22 | 65.75 | 97.59 | 143.05 | 216.00 | 301.74 | 382.57 | 0.337 |
| Tscha-1 | 11.78 | 6.54 | 9.69 | 14.49 | 22.02 | 35.52 | 54.00 | 81.10 | 121.31 | 178.34 | 257.44 | 361.61 | 476.97 | 0.385 |
| Tsu-0 | 12.55 | 5.93 | 8.83 | 14.66 | 22.10 | 35.70 | 55.61 | 83.85 | 126.76 | 191.40 | 268.75 | 343.54 | 421.63 | 0.409 |
| Tsu-1 | 15.11 | 7.28 | 10.59 | 15.58 | 26.06 | 40.67 | 62.32 | 93.81 | 142.18 | 212.71 | 297.76 | 382.48 | 457.06 | 0.364 |
| Tu-0 | 13.83 | 6.71 | 9.96 | 15.18 | 25.09 | 39.90 | 61.52 | 92.87 | 137.69 | 206.68 | 298.97 | 398.62 | 483.17 | 0.396 |
| Tul-0 | 11.28 | 6.51 | 9.56 | 14.32 | 21.76 | 33.74 | 51.56 | 77.73 | 113.47 | 169.27 | 241.90 | 326.95 | 396.69 | 0.382 |
| Ty-0 | 11.34 | 6.80 | 9.60 | 14.19 | 21.12 | 32.98 | 50.15 | 75.15 | 108.93 | 161.13 | 236.51 | 319.93 | 415.46 | 0.356 |
| Udul1-34 | 8.93 | 4.60 | 6.63 | 9.94 | 15.47 | 25.68 | 40.53 | 61.55 | 91.91 | 139.60 | 209.18 | 297.98 | 412.90 | 0.389 |
| Uk-1 | 10.03 | 4.41 | 6.37 | 10.11 | 15.24 | 24.73 | 38.05 | 57.60 | 87.63 | 135.35 | 208.45 | 295.86 | 392.19 | 0.394 |
| Uk-2 | 9.65 | 4.98 | 7.30 | 11.64 | 17.74 | 27.91 | 41.93 | 62.69 | 93.92 | 142.44 | 214.41 | 307.85 | 407.75 | 0.405 |
| Uk-4 | 9.76 | 4.70 | 6.93 | 11.16 | 17.60 | 28.39 | 42.86 | 65.19 | 97.68 | 143.90 | 221.80 | 324.89 | 434.28 | 0.421 |
| UKID48 | 8.06 | 5.84 | 8.35 | 12.47 | 18.71 | 29.69 | 44.65 | 65.01 | 95.65 | 137.81 | 193.36 | 263.49 | 330.88 | 0.367 |
| UKNW06- | 8.76 | 5.03 | 7.24 | 11.42 | 16.79 | 27.00 | 41.77 | 62.61 | 93.16 | 140.02 | 210.97 | 298.68 | 395.04 | 0.387 |
| UKNW06- | 9.60 | 5.37 | 7.88 | 12.23 | 18.15 | 29.07 | 44.29 | 67.02 | 99.02 | 150.21 | 223.07 | 315.20 | 411.62 | 0.384 |
| UKNW06- | 10.54 | 6.77 | 9.11 | 13.38 | 19.66 | 30.78 | 45.89 | 67.98 | 102.24 | 153.59 | 225.40 | 322.08 | 414.74 | 0.327 |
| UKSE06-4 | 8.56 | 5.31 | 7.63 | 11.09 | 16.32 | 25.48 | 37.39 | 55.44 | 81.43 | 124.50 | 188.37 | 288.79 | 382.05 | 0.367 |
| UKSE06-4 | 11.56 | 6.42 | 9.19 | 13.42 | 20.32 | 32.43 | 50.49 | 76.33 | 112.66 | 166.37 | 244.21 | 343.38 | 435.30 | 0.356 |
| UKSE06-4 | 10.56 | 5.94 | 8.62 | 13.13 | 20.01 | 31.41 | 46.99 | 69.71 | 107.51 | 164.31 | 238.38 | 331.04 | 428.78 | 0.382 |
| UKSE06-5 | 7.39 | 4.68 | 6.84 | 10.08 | 14.95 | 24.09 | 36.81 | 54.58 | 79.70 | 119.26 | 176.44 | 253.22 | 350.23 | 0.366 |
| UKSE06-6 | 10.54 | 5.53 | 8.26 | 12.82 | 18.35 | 28.95 | 43.44 | 64.68 | 96.98 | 148.27 | 215.50 | 295.73 | 377.26 | 0.398 |
| Ull2-3 | 12.01 | 6.06 | 9.06 | 14.06 | 21.62 | 34.36 | 52.75 | 80.84 | 121.50 | 185.07 | 278.19 | 386.59 | 486.17 | 0.408 |
| Ull-2-5 | 9.86 | 5.94 | 8.41 | 12.62 | 19.33 | 30.88 | 47.40 | 72.10 | 107.29 | 158.28 | 233.14 | 320.50 | 424.68 | 0.359 |
| Uod-1 | 9.06 | 4.34 | 6.46 | 9.34 | 15.60 | 25.22 | 38.48 | 58.04 | 87.88 | 135.98 | 201.21 | 275.90 | 356.86 | 0.368 |
| Uod-7 | 9.24 | 5.30 | 7.56 | 11.05 | 15.91 | 25.44 | 39.32 | 60.52 | 90.80 | 135.94 | 202.59 | 287.84 | 379.26 | 0.359 |
| Utrecht | 9.16 | 4.89 | 7.30 | 11.27 | 16.60 | 26.40 | 40.91 | 63.32 | 95.68 | 146.00 | 214.29 | 290.06 | 380.30 | 0.385 |
| Van-0 | 11.15 | 6.36 | 9.15 | 13.65 | 21.91 | 35.25 | 54.07 | 82.47 | 123.26 | 185.08 | 274.61 | 385.08 | 500.05 | 0.374 |
| Var-2-1 | 6.40 | 5.48 | 7.30 | 10.45 | 14.76 | 22.69 | 33.04 | 48.18 | 69.83 | 100.57 | 142.95 | 197.70 | 253.82 | 0.285 |
| Ven-1 | 9.99 | 5.97 | 8.67 | 13.51 | 19.76 | 30.70 | 46.56 | 70.99 | 102.65 | 147.78 | 209.83 | 272.00 | 331.19 | 0.372 |
| Wa-1 | 11.36 | 5.78 | 8.44 | 12.81 | 19.79 | 31.98 | 49.46 | 77.01 | 114.78 | 169.13 | 245.26 | 342.64 | 447.06 | 0.392 |

| accession | DW20 | PLA07 | PLA08 | PLA09 | PLA10 | PLA11 | PLA12 | PLA13 | PLA14 | PLA15 | PLA16 | PLA17 | PLA18 | RGR07_09 |
|---|---|---|---|---|---|---|---|---|---|---|---|---|---|---|
| Wa-1 | 14.25 | 7.41 | 11.32 | 16.96 | 26.23 | 41.57 | 64.93 | 98.79 | 146.57 | 217.35 | 318.37 | 427.91 | 527.08 | 0.406 |
| Wag-3 | 10.86 | 5.92 | 8.22 | 12.76 | 19.05 | 31.25 | 49.48 | 75.26 | 111.11 | 165.36 | 243.63 | 331.37 | 420.69 | 0.360 |
| Wag-4 | 14.46 | 8.03 | 11.17 | 16.27 | 25.16 | 40.76 | 64.66 | 97.52 | 146.17 | 222.76 | 322.43 | 421.80 | 509.74 | 0.345 |
| Wag-5 | 10.78 | 6.33 | 9.15 | 13.58 | 20.34 | 32.56 | 50.45 | 74.71 | 109.45 | 165.05 | 241.24 | 325.64 | 415.63 | 0.368 |
| WAR | 12.09 | 6.41 | 9.45 | 14.31 | 21.77 | 34.80 | 53.01 | 80.11 | 119.55 | 177.66 | 260.29 | 352.18 | 457.74 | 0.388 |
| Wc-2 | 14.23 | 5.89 | 9.00 | 14.16 | 22.19 | 36.26 | 57.07 | 89.62 | 137.86 | 207.39 | 300.04 | 407.83 | 498.82 | 0.427 |
| Wei-(1) | 14.44 | 7.55 | 11.49 | 17.63 | 26.68 | 41.88 | 63.81 | 94.88 | 140.09 | 207.56 | 278.69 | 340.46 | 408.80 | 0.412 |
| Wei-0 | 11.43 | 5.42 | 8.23 | 12.61 | 18.89 | 30.01 | 45.61 | 68.20 | 102.37 | 155.64 | 229.48 | 307.75 | 385.56 | 0.411 |
| Wil | 9.98 | 4.79 | 6.72 | 10.73 | 17.19 | 28.58 | 45.33 | 69.43 | 103.51 | 154.28 | 231.85 | 331.88 | 441.55 | 0.394 |
| Wil-2 | 10.02 | 5.09 | 7.33 | 11.61 | 18.72 | 30.54 | 47.47 | 72.19 | 109.45 | 167.77 | 250.41 | 352.15 | 464.81 | 0.371 |
| Wl-0 | 12.02 | 6.37 | 9.48 | 14.36 | 21.81 | 34.99 | 55.16 | 84.12 | 127.01 | 195.55 | 295.45 | 403.91 | 503.19 | 0.398 |
| Ws | 10.61 | 5.04 | 7.17 | 10.98 | 16.25 | 26.02 | 41.26 | 64.80 | 98.32 | 146.80 | 221.20 | 321.06 | 416.53 | 0.379 |
| Ws-0 | 13.36 | 6.69 | 10.02 | 15.22 | 23.85 | 39.00 | 60.77 | 92.93 | 143.83 | 217.34 | 308.21 | 404.87 | 533.95 | 0.394 |
| Ws-2 | 11.41 | 5.18 | 7.44 | 11.27 | 16.96 | 27.22 | 42.94 | 66.87 | 101.58 | 152.47 | 232.70 | 339.98 | 447.85 | 0.375 |
| Ws-3 | 12.66 | 6.41 | 9.02 | 13.57 | 20.67 | 33.38 | 52.21 | 80.37 | 118.33 | 175.40 | 264.83 | 372.79 | 474.06 | 0.363 |
| Wt-3 | 13.04 | 6.48 | 9.35 | 14.70 | 23.57 | 38.54 | 59.48 | 88.41 | 133.84 | 205.76 | 284.30 | 374.13 | 466.81 | 0.400 |
| Wt-5 | 9.16 | 5.24 | 7.29 | 11.03 | 16.27 | 25.71 | 39.34 | 59.78 | 90.76 | 136.21 | 209.49 | 300.29 | 390.82 | 0.362 |
| Yo-0 | 11.72 | 6.65 | 9.89 | 15.31 | 23.14 | 36.39 | 55.28 | 83.52 | 122.91 | 185.99 | 267.29 | 357.49 | 443.91 | 0.398 |
| Zdr-1 | 10.54 | 5.12 | 7.85 | 12.17 | 18.77 | 29.60 | 45.31 | 69.46 | 105.60 | 162.12 | 246.32 | 350.71 | 476.95 | 0.417 |
| Zdr-6 | 10.93 | 5.90 | 8.40 | 12.88 | 20.18 | 32.37 | 48.43 | 72.08 | 105.64 | 153.43 | 219.76 | 298.79 | 394.54 | 0.373 |
| Zdrl2-24 | 12.84 | 7.19 | 10.81 | 16.65 | 25.66 | 40.43 | 61.80 | 92.09 | 134.42 | 198.62 | 291.06 | 388.42 | 479.49 | 0.409 |
| Zdrl2-25 | 11.85 | 5.49 | 8.40 | 13.49 | 21.05 | 34.05 | 51.81 | 78.96 | 119.39 | 181.64 | 273.73 | 378.34 | 489.48 | 0.428 |
| Zü-1 | 13.73 | 7.95 | 11.69 | 17.53 | 26.03 | 40.87 | 63.19 | 94.17 | 138.51 | 204.00 | 290.37 | 385.69 | 486.33 | 0.383 |

| accession | RGR08_10 | RGR09_11 | RGR10_12 | RGR11_13 | RGR12_14 | RGR13_15 | RGR14_16 | RGR15_17 | RGR16_18 | SA | SL | SW |
|---|---|---|---|---|---|---|---|---|---|---|---|---|
| $H^2$ | 0.7997 | 0.8173 | 0.8100 | 0.8359 | 0.8304 | 0.8179 | 0.7191 | 0.6622 | 0.6722 | 0.933 | 0.895 | 0.905 |
| 11PNA4 | 0.402 | 0.441 | 0.451 | 0.412 | 0.387 | 0.377 | 0.360 | 0.323 | 0.268 | 0.148 | 0.535 | 0.393 |
| 328PNA | 0.363 | 0.471 | 0.498 | 0.419 | 0.393 | 0.362 | 0.344 | 0.326 | 0.229 | 0.138 | 0.512 | 0.380 |
| Aa-0 | 0.399 | 0.469 | 0.478 | 0.434 | 0.414 | 0.397 | 0.399 | 0.368 | 0.290 | 0.131 | 0.532 | 0.350 |
| Ak-1 | 0.415 | 0.428 | 0.427 | 0.402 | 0.386 | 0.393 | 0.402 | 0.369 | 0.310 | 0.130 | 0.526 | 0.350 |
| Akita | 0.425 | 0.478 | 0.456 | 0.429 | 0.403 | 0.403 | 0.407 | 0.363 | 0.311 | 0.125 | 0.529 | 0.341 |
| Alc-0 | 0.405 | 0.450 | 0.466 | 0.440 | 0.393 | 0.370 | 0.366 | 0.314 | 0.233 | 0.106 | 0.455 | 0.336 |
| ALL1-2 | 0.411 | 0.454 | 0.443 | 0.407 | 0.400 | 0.386 | 0.374 | 0.343 | 0.284 | 0.120 | 0.495 | 0.347 |
| Alst-1 | 0.415 | 0.438 | 0.427 | 0.408 | 0.402 | 0.388 | 0.356 | 0.307 | 0.257 | 0.130 | 0.510 | 0.363 |
| Amel-1 | 0.373 | 0.405 | 0.414 | 0.390 | 0.369 | 0.368 | 0.374 | 0.359 | 0.320 | 0.149 | 0.539 | 0.391 |
| An1 | 0.420 | 0.457 | 0.460 | 0.428 | 0.400 | 0.401 | 0.414 | 0.366 | 0.295 | 0.138 | 0.529 | 0.370 |
| An-2 | 0.444 | 0.452 | 0.438 | 0.406 | 0.400 | 0.411 | 0.391 | 0.324 | 0.255 | 0.128 | 0.518 | 0.354 |
| Ang-0 | 0.426 | 0.458 | 0.464 | 0.423 | 0.401 | 0.396 | 0.390 | 0.363 | 0.307 | 0.107 | 0.484 | 0.322 |
| Ang-0 | 0.425 | 0.464 | 0.442 | 0.415 | 0.412 | 0.412 | 0.396 | 0.345 | 0.262 | 0.125 | 0.520 | 0.341 |
| Ang-1 | 0.437 | 0.468 | 0.444 | 0.402 | 0.389 | 0.397 | 0.399 | 0.353 | 0.265 | 0.130 | 0.495 | 0.367 |
| Ann-1 | 0.416 | 0.463 | 0.479 | 0.453 | 0.425 | 0.400 | 0.369 | 0.312 | 0.245 | 0.135 | 0.540 | 0.355 |
| Arby-1 | 0.400 | 0.441 | 0.459 | 0.421 | 0.400 | 0.421 | 0.424 | 0.383 | 0.307 | 0.112 | 0.498 | 0.321 |
| Ba-1 | 0.397 | 0.488 | 0.487 | 0.444 | 0.418 | 0.417 | 0.416 | 0.360 | 0.265 | 0.110 | 0.482 | 0.325 |
| Ba1-2 | 0.442 | 0.463 | 0.452 | 0.431 | 0.419 | 0.415 | 0.414 | 0.365 | 0.296 | 0.117 | 0.497 | 0.340 |
| Baa-1 | 0.398 | 0.442 | 0.450 | 0.428 | 0.405 | 0.389 | 0.385 | 0.364 | 0.295 | 0.148 | 0.554 | 0.375 |
| Bay-0 | 0.448 | 0.466 | 0.460 | 0.428 | 0.416 | 0.408 | 0.382 | 0.336 | 0.263 | 0.125 | 0.518 | 0.345 |
| Bch-1 | 0.438 | 0.470 | 0.470 | 0.438 | 0.419 | 0.424 | 0.425 | 0.369 | 0.270 | 0.136 | 0.527 | 0.371 |
| Bd-0 | 0.402 | 0.455 | 0.464 | 0.436 | 0.405 | 0.384 | 0.389 | 0.388 | 0.330 | 0.121 | 0.495 | 0.355 |
| Be-0 | 0.390 | 0.443 | 0.440 | 0.395 | 0.372 | 0.401 | 0.422 | 0.386 | 0.324 | 0.120 | 0.488 | 0.349 |
| Be-1 | 0.426 | 0.466 | 0.460 | 0.433 | 0.417 | 0.418 | 0.422 | 0.385 | 0.315 | 0.119 | 0.491 | 0.344 |
| Belmonte4 | 0.394 | 0.416 | 0.407 | 0.387 | 0.375 | 0.370 | 0.377 | 0.365 | 0.321 | 0.178 | 0.566 | 0.438 |
| Benk-1 | 0.384 | 0.446 | 0.437 | 0.410 | 0.392 | 0.387 | 0.377 | 0.326 | 0.261 | 0.226 | 0.648 | 0.478 |
| Bg-2 | 0.378 | 0.431 | 0.435 | 0.401 | 0.390 | 0.410 | 0.419 | 0.393 | 0.354 | 0.169 | 0.556 | 0.427 |
| Bl-1 | 0.403 | 0.493 | 0.463 | 0.426 | 0.406 | 0.390 | 0.396 | 0.372 | 0.280 | 0.163 | 0.553 | 0.411 |
| Bla-1 | 0.399 | 0.443 | 0.444 | 0.419 | 0.392 | 0.382 | 0.373 | 0.319 | 0.232 | 0.192 | 0.601 | 0.445 |

| accession | RGR08_10 | RGR09_11 | RGR10_12 | RGR11_13 | RGR12_14 | RGR13_15 | RGR14_16 | RGR15_17 | RGR16_18 | SA | SL | SW |
|---|---|---|---|---|---|---|---|---|---|---|---|---|
| Bla-11 | 0.390 | 0.430 | 0.442 | 0.419 | 0.399 | 0.391 | 0.399 | 0.379 | 0.327 | 0.188 | 0.591 | 0.446 |
| Blh-1 | 0.418 | 0.463 | 0.469 | 0.431 | 0.402 | 0.397 | 0.382 | 0.331 | 0.272 | 0.167 | 0.558 | 0.419 |
| Blh-1 | 0.398 | 0.453 | 0.445 | 0.421 | 0.412 | 0.403 | 0.370 | 0.304 | 0.222 | 0.183 | 0.600 | 0.425 |
| Blh-2 | 0.393 | 0.451 | 0.460 | 0.431 | 0.401 | 0.403 | 0.407 | 0.351 | 0.272 | 0.194 | 0.616 | 0.434 |
| Boot-1 | 0.410 | 0.434 | 0.440 | 0.431 | 0.419 | 0.409 | 0.399 | 0.365 | 0.310 | 0.171 | 0.572 | 0.421 |
| Bor-1 | 0.410 | 0.430 | 0.420 | 0.397 | 0.375 | 0.366 | 0.383 | 0.374 | 0.329 | 0.145 | 0.521 | 0.394 |
| Bor-4 | 0.420 | 0.463 | 0.460 | 0.422 | 0.407 | 0.418 | 0.412 | 0.362 | 0.295 | 0.154 | 0.540 | 0.405 |
| Br-0 | 0.429 | 0.476 | 0.467 | 0.433 | 0.420 | 0.412 | 0.362 | 0.281 | 0.216 | 0.193 | 0.613 | 0.427 |
| Bs-1 | 0.382 | 0.435 | 0.435 | 0.409 | 0.379 | 0.388 | 0.426 | 0.405 | 0.333 | 0.190 | 0.600 | 0.443 |
| Bs-2 | 0.391 | 0.438 | 0.433 | 0.413 | 0.410 | 0.406 | 0.391 | 0.351 | 0.296 | 0.191 | 0.612 | 0.435 |
| Bsch-0 | 0.438 | 0.481 | 0.467 | 0.424 | 0.399 | 0.394 | 0.405 | 0.370 | 0.281 | 0.161 | 0.548 | 0.406 |
| Bsch-2 | 0.437 | 0.469 | 0.445 | 0.417 | 0.412 | 0.409 | 0.395 | 0.343 | 0.260 | 0.191 | 0.607 | 0.442 |
| Bu-0 | 0.405 | 0.450 | 0.468 | 0.430 | 0.395 | 0.374 | 0.345 | 0.303 | 0.256 | 0.181 | 0.599 | 0.426 |
| Bu-2 | 0.403 | 0.450 | 0.456 | 0.421 | 0.403 | 0.391 | 0.387 | 0.338 | 0.262 | 0.192 | 0.589 | 0.452 |
| Bur-0 | 0.355 | 0.423 | 0.417 | 0.382 | 0.363 | 0.358 | 0.375 | 0.364 | 0.319 | 0.237 | 0.676 | 0.478 |
| C24 | 0.383 | 0.423 | 0.437 | 0.393 | 0.374 | 0.386 | 0.392 | 0.357 | 0.310 | 0.175 | 0.564 | 0.425 |
| Ca-0 | 0.397 | 0.440 | 0.443 | 0.413 | 0.400 | 0.388 | 0.376 | 0.347 | 0.276 | 0.165 | 0.552 | 0.420 |
| Cal-0 | 0.396 | 0.447 | 0.423 | 0.393 | 0.379 | 0.387 | 0.383 | 0.338 | 0.271 | 0.209 | 0.637 | 0.454 |
| CAM-16 | 0.409 | 0.449 | 0.435 | 0.404 | 0.400 | 0.417 | 0.414 | 0.375 | 0.320 | 0.164 | 0.555 | 0.416 |
| CAM-61 | 0.405 | 0.424 | 0.427 | 0.415 | 0.393 | 0.391 | 0.392 | 0.349 | 0.279 | 0.217 | 0.610 | 0.496 |
| Can-0 | 0.436 | 0.441 | 0.431 | 0.408 | 0.409 | 0.418 | 0.392 | 0.334 | 0.255 | 0.217 | 0.631 | 0.480 |
| Cen-0 | 0.364 | 0.474 | 0.486 | 0.415 | 0.388 | 0.384 | 0.405 | 0.394 | 0.342 | 0.184 | 0.584 | 0.449 |
| Cha-0 | 0.415 | 0.458 | 0.453 | 0.426 | 0.402 | 0.393 | 0.389 | 0.337 | 0.232 | 0.209 | 0.602 | 0.470 |
| Chat-1 | 0.388 | 0.456 | 0.466 | 0.438 | 0.419 | 0.414 | 0.399 | 0.359 | 0.302 | 0.176 | 0.557 | 0.443 |
| Chi-0 | 0.421 | 0.467 | 0.468 | 0.436 | 0.416 | 0.414 | 0.411 | 0.371 | 0.297 | 0.178 | 0.569 | 0.437 |
| CIBC-17 | 0.397 | 0.456 | 0.444 | 0.417 | 0.411 | 0.403 | 0.389 | 0.341 | 0.294 | 0.208 | 0.601 | 0.481 |
| CIBC-2 | 0.409 | 0.470 | 0.473 | 0.428 | 0.391 | 0.371 | 0.356 | 0.315 | 0.266 | 0.218 | 0.615 | 0.488 |
| CIBC-4 | 0.398 | 0.455 | 0.438 | 0.408 | 0.392 | 0.396 | 0.384 | 0.328 | 0.261 | 0.220 | 0.650 | 0.469 |
| CIBC-5 | 0.405 | 0.442 | 0.444 | 0.409 | 0.376 | 0.369 | 0.379 | 0.350 | 0.289 | 0.187 | 0.576 | 0.453 |
| Cit-0 | 0.376 | 0.419 | 0.435 | 0.414 | 0.409 | 0.397 | 0.388 | 0.369 | 0.307 | 0.202 | 0.588 | 0.486 |

| accession | RGR08_10 | RGR09_11 | RGR10_12 | RGR11_13 | RGR12_14 | RGR13_15 | RGR14_16 | RGR15_17 | RGR16_18 | SA | SL | SW |
|---|---|---|---|---|---|---|---|---|---|---|---|---|
| Cl-0 | 0.419 | 0.453 | 0.433 | 0.413 | 0.423 | 0.419 | 0.386 | 0.324 | 0.275 | 0.200 | 0.612 | 0.454 |
| Cnt-1 | 0.378 | 0.392 | 0.401 | 0.396 | 0.397 | 0.398 | 0.387 | 0.353 | 0.315 | 0.225 | 0.628 | 0.490 |
| Co | 0.400 | 0.439 | 0.455 | 0.421 | 0.400 | 0.397 | 0.408 | 0.398 | 0.333 | 0.182 | 0.567 | 0.446 |
| Co-2 | 0.414 | 0.481 | 0.456 | 0.424 | 0.400 | 0.397 | 0.382 | 0.333 | 0.278 | 0.194 | 0.600 | 0.442 |
| Co-3 | 0.417 | 0.456 | 0.446 | 0.411 | 0.395 | 0.394 | 0.405 | 0.391 | 0.341 | 0.161 | 0.546 | 0.418 |
| Co-4 | 0.389 | 0.438 | 0.415 | 0.401 | 0.399 | 0.396 | 0.399 | 0.369 | 0.314 | 0.217 | 0.651 | 0.468 |
| Col-0 | 0.420 | 0.466 | 0.451 | 0.420 | 0.405 | 0.393 | 0.393 | 0.367 | 0.300 | 0.215 | 0.625 | 0.485 |
| Com-1 | 0.393 | 0.428 | 0.419 | 0.395 | 0.385 | 0.384 | 0.400 | 0.386 | 0.306 | 0.196 | 0.590 | 0.467 |
| CSHL-5 | 0.422 | 0.453 | 0.443 | 0.432 | 0.422 | 0.399 | 0.403 | 0.389 | 0.280 | 0.190 | 0.574 | 0.460 |
| Ct-1 | 0.425 | 0.451 | 0.446 | 0.426 | 0.409 | 0.381 | 0.352 | 0.316 | 0.242 | 0.214 | 0.624 | 0.474 |
| CUR-3 | 0.308 | 0.414 | 0.424 | 0.345 | 0.333 | 0.343 | 0.377 | 0.394 | 0.392 | 0.181 | 0.565 | 0.448 |
| Cvi-0 | 0.318 | 0.345 | 0.358 | 0.330 | 0.318 | 0.309 | 0.308 | 0.348 | 0.341 | 0.220 | 0.636 | 0.478 |
| Da(1)-12 | 0.441 | 0.475 | 0.455 | 0.416 | 0.391 | 0.381 | 0.388 | 0.350 | 0.259 | 0.229 | 0.643 | 0.494 |
| Da-0 | 0.405 | 0.469 | 0.475 | 0.454 | 0.427 | 0.395 | 0.380 | 0.348 | 0.297 | 0.178 | 0.566 | 0.441 |
| Db-0 | 0.419 | 0.464 | 0.480 | 0.443 | 0.419 | 0.404 | 0.401 | 0.381 | 0.332 | 0.155 | 0.546 | 0.401 |
| Db-1 | 0.431 | 0.463 | 0.461 | 0.426 | 0.402 | 0.392 | 0.380 | 0.354 | 0.292 | 0.181 | 0.588 | 0.435 |
| Di-1 | 0.373 | 0.419 | 0.428 | 0.391 | 0.390 | 0.418 | 0.409 | 0.355 | 0.307 | 0.207 | 0.606 | 0.470 |
| Dijon-M | 0.429 | 0.454 | 0.446 | 0.436 | 0.423 | 0.394 | 0.379 | 0.364 | 0.280 | 0.215 | 0.620 | 0.483 |
| Do-0 | 0.426 | 0.452 | 0.448 | 0.424 | 0.412 | 0.399 | 0.371 | 0.321 | 0.250 | 0.190 | 0.616 | 0.430 |
| Dr-0 | 0.438 | 0.488 | 0.470 | 0.431 | 0.416 | 0.408 | 0.405 | 0.360 | 0.259 | 0.195 | 0.611 | 0.447 |
| Dra-0 | 0.445 | 0.471 | 0.456 | 0.405 | 0.381 | 0.381 | 0.392 | 0.343 | 0.252 | 0.170 | 0.584 | 0.409 |
| Dra-2 | 0.411 | 0.472 | 0.441 | 0.402 | 0.390 | 0.406 | 0.407 | 0.377 | 0.312 | 0.196 | 0.617 | 0.442 |
| DraIV1-14 | 0.433 | 0.454 | 0.443 | 0.416 | 0.396 | 0.393 | 0.392 | 0.347 | 0.295 | 0.160 | 0.547 | 0.413 |
| DraIV1-5 | 0.423 | 0.499 | 0.465 | 0.430 | 0.398 | 0.381 | 0.373 | 0.331 | 0.264 | 0.165 | 0.579 | 0.400 |
| DraIV6-16 | 0.406 | 0.447 | 0.440 | 0.424 | 0.418 | 0.404 | 0.382 | 0.338 | 0.269 | 0.181 | 0.572 | 0.443 |
| DraIV6-35 | 0.462 | 0.492 | 0.439 | 0.409 | 0.412 | 0.413 | 0.382 | 0.331 | 0.273 | 0.181 | 0.575 | 0.439 |
| Duk | 0.431 | 0.456 | 0.445 | 0.425 | 0.405 | 0.406 | 0.407 | 0.366 | 0.314 | 0.168 | 0.557 | 0.419 |
| Durh-1 | 0.354 | 0.383 | 0.398 | 0.375 | 0.354 | 0.351 | 0.347 | 0.332 | 0.313 | 0.155 | 0.540 | 0.409 |
| Ede-1 | 0.407 | 0.430 | 0.432 | 0.416 | 0.396 | 0.396 | 0.413 | 0.371 | 0.306 | 0.142 | 0.528 | 0.380 |
| Eden-1 | 0.393 | 0.423 | 0.438 | 0.407 | 0.378 | 0.352 | 0.344 | 0.322 | 0.267 | 0.235 | 0.647 | 0.489 |

| accession | RGR08_10 | RGR09_11 | RGR10_12 | RGR11_13 | RGR12_14 | RGR13_15 | RGR14_16 | RGR15_17 | RGR16_18 | SA | SL | SW |
|---|---|---|---|---|---|---|---|---|---|---|---|---|
| Edi-0 | 0.417 | 0.470 | 0.441 | 0.405 | 0.395 | 0.383 | 0.368 | 0.335 | 0.272 | 0.190 | 0.574 | 0.459 |
| Ei-2 | 0.424 | 0.456 | 0.456 | 0.431 | 0.404 | 0.406 | 0.417 | 0.361 | 0.296 | 0.210 | 0.606 | 0.481 |
| Ei-4 | 0.426 | 0.454 | 0.451 | 0.419 | 0.410 | 0.403 | 0.383 | 0.345 | 0.290 | 0.161 | 0.562 | 0.402 |
| El-0 | 0.425 | 0.470 | 0.460 | 0.408 | 0.388 | 0.390 | 0.385 | 0.358 | 0.281 | 0.175 | 0.574 | 0.424 |
| En-1 | 0.413 | 0.441 | 0.429 | 0.425 | 0.417 | 0.388 | 0.371 | 0.334 | 0.246 | 0.189 | 0.622 | 0.421 |
| Enkheim-D | 0.400 | 0.447 | 0.468 | 0.441 | 0.414 | 0.411 | 0.411 | 0.352 | 0.280 | 0.196 | 0.599 | 0.461 |
| Ep-0 | 0.424 | 0.462 | 0.462 | 0.439 | 0.425 | 0.416 | 0.395 | 0.327 | 0.252 | 0.194 | 0.595 | 0.456 |
| Er-0 | 0.429 | 0.463 | 0.464 | 0.436 | 0.422 | 0.426 | 0.408 | 0.356 | 0.306 | 0.176 | 0.573 | 0.433 |
| Es-0 | 0.439 | 0.471 | 0.463 | 0.433 | 0.404 | 0.408 | 0.396 | 0.315 | 0.238 | 0.213 | 0.628 | 0.474 |
| Est-0 | 0.456 | 0.513 | 0.467 | 0.431 | 0.411 | 0.403 | 0.397 | 0.365 | 0.309 | 0.223 | 0.623 | 0.494 |
| Est-1 | 0.421 | 0.482 | 0.481 | 0.447 | 0.431 | 0.415 | 0.402 | 0.342 | 0.246 | 0.217 | 0.644 | 0.469 |
| Fei-0 | 0.406 | 0.451 | 0.450 | 0.425 | 0.411 | 0.412 | 0.396 | 0.338 | 0.276 | 0.185 | 0.574 | 0.447 |
| Fi-0 | 0.394 | 0.449 | 0.443 | 0.413 | 0.387 | 0.390 | 0.393 | 0.364 | 0.321 | 0.206 | 0.600 | 0.474 |
| Fi-1 | 0.415 | 0.456 | 0.455 | 0.425 | 0.409 | 0.407 | 0.404 | 0.356 | 0.276 | 0.189 | 0.589 | 0.447 |
| Fr-2 | 0.404 | 0.496 | 0.454 | 0.420 | 0.403 | 0.390 | 0.386 | 0.351 | 0.288 | 0.218 | 0.645 | 0.475 |
| Fr-4 | 0.420 | 0.456 | 0.466 | 0.435 | 0.409 | 0.403 | 0.380 | 0.329 | 0.281 | 0.184 | 0.574 | 0.451 |
| Ga-0 | 0.406 | 0.451 | 0.457 | 0.429 | 0.408 | 0.410 | 0.413 | 0.390 | 0.338 | 0.192 | 0.607 | 0.442 |
| Ga-2 | 0.413 | 0.461 | 0.481 | 0.451 | 0.422 | 0.408 | 0.409 | 0.375 | 0.301 | 0.183 | 0.572 | 0.447 |
| Gd-1 | 0.433 | 0.471 | 0.472 | 0.439 | 0.423 | 0.413 | 0.399 | 0.337 | 0.262 | 0.195 | 0.618 | 0.441 |
| Ge-1 | 0.415 | 0.449 | 0.441 | 0.426 | 0.422 | 0.406 | 0.393 | 0.363 | 0.302 | 0.178 | 0.564 | 0.440 |
| Ge-2 | 0.408 | 0.449 | 0.463 | 0.412 | 0.377 | 0.387 | 0.392 | 0.328 | 0.248 | 0.191 | 0.584 | 0.455 |
| Gel-1 | 0.416 | 0.473 | 0.459 | 0.434 | 0.414 | 0.405 | 0.405 | 0.376 | 0.302 | 0.174 | 0.560 | 0.433 |
| Gie-0 | 0.429 | 0.460 | 0.457 | 0.429 | 0.413 | 0.405 | 0.396 | 0.375 | 0.329 | 0.184 | 0.580 | 0.442 |
| Go-0 | 0.419 | 0.432 | 0.430 | 0.416 | 0.413 | 0.412 | 0.412 | 0.388 | 0.342 | 0.218 | 0.606 | 0.502 |
| Gö-2 | 0.419 | 0.455 | 0.459 | 0.437 | 0.412 | 0.406 | 0.403 | 0.373 | 0.313 | 0.180 | 0.559 | 0.455 |
| Golm-1 | 0.415 | 0.510 | 0.489 | 0.444 | 0.426 | 0.436 | 0.437 | 0.387 | 0.311 | 0.176 | 0.562 | 0.440 |
| GOT-7 | 0.368 | 0.428 | 0.435 | 0.413 | 0.392 | 0.362 | 0.356 | 0.332 | 0.256 | 0.215 | 0.643 | 0.463 |
| Gr | 0.402 | 0.444 | 0.451 | 0.430 | 0.399 | 0.392 | 0.411 | 0.386 | 0.329 | 0.148 | 0.517 | 0.401 |
| Gr-1 | 0.413 | 0.441 | 0.451 | 0.429 | 0.405 | 0.399 | 0.408 | 0.385 | 0.343 | 0.151 | 0.510 | 0.410 |
| Gr-5 | 0.409 | 0.491 | 0.445 | 0.422 | 0.417 | 0.428 | 0.421 | 0.385 | 0.316 | 0.174 | 0.569 | 0.427 |

| accession | RGR08_10 | RGR09_11 | RGR10_12 | RGR11_13 | RGR12_14 | RGR13_15 | RGR14_16 | RGR15_17 | RGR16_18 | SA | SL | SW |
|---|---|---|---|---|---|---|---|---|---|---|---|---|
| Gre-0 | 0.394 | 0.436 | 0.428 | 0.406 | 0.400 | 0.393 | 0.381 | 0.335 | 0.250 | 0.217 | 0.622 | 0.486 |
| Gu-0 | 0.396 | 0.522 | 0.476 | 0.421 | 0.398 | 0.404 | 0.402 | 0.351 | 0.288 | 0.185 | 0.598 | 0.433 |
| Gu-1 | 0.406 | 0.442 | 0.447 | 0.427 | 0.410 | 0.409 | 0.424 | 0.394 | 0.314 | 0.169 | 0.586 | 0.407 |
| Gy-0 | 0.396 | 0.437 | 0.425 | 0.385 | 0.366 | 0.370 | 0.370 | 0.339 | 0.311 | 0.177 | 0.575 | 0.432 |
| Ha-0 | 0.449 | 0.467 | 0.453 | 0.420 | 0.415 | 0.417 | 0.388 | 0.321 | 0.247 | 0.211 | 0.618 | 0.479 |
| Hau-0 | 0.398 | 0.431 | 0.430 | 0.410 | 0.401 | 0.397 | 0.396 | 0.343 | 0.256 | 0.195 | 0.589 | 0.463 |
| Hey-1 | 0.399 | 0.447 | 0.456 | 0.417 | 0.400 | 0.399 | 0.400 | 0.353 | 0.270 | 0.198 | 0.608 | 0.454 |
| Hh-0 | 0.405 | 0.445 | 0.463 | 0.432 | 0.416 | 0.404 | 0.396 | 0.368 | 0.301 | 0.143 | 0.514 | 0.396 |
| Hi-0 | 0.412 | 0.454 | 0.447 | 0.414 | 0.395 | 0.407 | 0.424 | 0.407 | 0.342 | 0.188 | 0.576 | 0.452 |
| Hl-3 | 0.431 | 0.460 | 0.462 | 0.432 | 0.412 | 0.405 | 0.400 | 0.354 | 0.289 | 0.152 | 0.540 | 0.396 |
| Hn-0 | 0.407 | 0.430 | 0.421 | 0.394 | 0.381 | 0.373 | 0.379 | 0.363 | 0.313 | 0.184 | 0.575 | 0.452 |
| Hod | 0.413 | 0.444 | 0.437 | 0.415 | 0.387 | 0.375 | 0.394 | 0.387 | 0.332 | 0.164 | 0.545 | 0.423 |
| HOG | 0.432 | 0.464 | 0.461 | 0.439 | 0.414 | 0.391 | 0.389 | 0.356 | 0.266 | 0.146 | 0.516 | 0.401 |
| Hoh-1 | 0.380 | 0.462 | 0.459 | 0.424 | 0.415 | 0.411 | 0.410 | 0.402 | 0.385 | 0.153 | 0.537 | 0.402 |
| Hovdala-2 | 0.384 | 0.486 | 0.491 | 0.430 | 0.399 | 0.389 | 0.377 | 0.354 | 0.294 | 0.156 | 0.537 | 0.411 |
| HR-10 | 0.410 | 0.428 | 0.432 | 0.414 | 0.397 | 0.402 | 0.407 | 0.376 | 0.315 | 0.159 | 0.547 | 0.409 |
| HR-5 | 0.411 | 0.434 | 0.438 | 0.417 | 0.409 | 0.404 | 0.402 | 0.369 | 0.306 | 0.149 | 0.517 | 0.409 |
| Hs-0 | 0.443 | 0.465 | 0.456 | 0.422 | 0.407 | 0.417 | 0.412 | 0.361 | 0.293 | 0.180 | 0.592 | 0.430 |
| HSm | 0.450 | 0.478 | 0.462 | 0.424 | 0.412 | 0.416 | 0.417 | 0.389 | 0.338 | 0.162 | 0.544 | 0.419 |
| In-0 | 0.404 | 0.463 | 0.464 | 0.437 | 0.415 | 0.410 | 0.419 | 0.390 | 0.312 | 0.191 | 0.594 | 0.454 |
| Is-1 | 0.447 | 0.489 | 0.480 | 0.431 | 0.414 | 0.414 | 0.421 | 0.383 | 0.310 | 0.146 | 0.523 | 0.384 |
| Je-0 | 0.404 | 0.466 | 0.473 | 0.438 | 0.410 | 0.388 | 0.375 | 0.337 | 0.274 | 0.179 | 0.593 | 0.421 |
| Je-54 | 0.421 | 0.475 | 0.478 | 0.441 | 0.402 | 0.381 | 0.388 | 0.380 | 0.352 | 0.143 | 0.524 | 0.386 |
| JEA | 0.372 | 0.405 | 0.412 | 0.362 | 0.324 | 0.317 | 0.335 | 0.337 | 0.299 | 0.247 | 0.653 | 0.517 |
| Jea | 0.381 | 0.413 | 0.423 | 0.378 | 0.326 | 0.316 | 0.338 | 0.340 | 0.306 | 0.276 | 0.720 | 0.532 |
| Jl-3 | 0.405 | 0.440 | 0.429 | 0.398 | 0.383 | 0.385 | 0.384 | 0.346 | 0.290 | 0.219 | 0.626 | 0.486 |
| Jm-1 | 0.443 | 0.450 | 0.440 | 0.423 | 0.412 | 0.406 | 0.404 | 0.372 | 0.318 | 0.204 | 0.593 | 0.473 |
| Kä-0 | 0.393 | 0.463 | 0.454 | 0.405 | 0.380 | 0.380 | 0.390 | 0.386 | 0.367 | 0.229 | 0.664 | 0.486 |
| Kas-2 | 0.413 | 0.429 | 0.426 | 0.404 | 0.392 | 0.368 | 0.335 | 0.307 | 0.263 | 0.301 | 0.697 | 0.584 |
| Kb-0 | 0.399 | 0.438 | 0.461 | 0.445 | 0.407 | 0.395 | 0.409 | 0.371 | 0.281 | 0.217 | 0.609 | 0.494 |

| accession | RGR08_10 | RGR09_11 | RGR10_12 | RGR11_13 | RGR12_14 | RGR13_15 | RGR14_16 | RGR15_17 | RGR16_18 | SA | SL | SW |
|---|---|---|---|---|---|---|---|---|---|---|---|---|
| KBS-Mac- | 0.385 | 0.410 | 0.410 | 0.391 | 0.394 | 0.402 | 0.404 | 0.360 | 0.287 | 0.263 | 0.673 | 0.537 |
| Kelsterbacl | 0.401 | 0.450 | 0.462 | 0.431 | 0.413 | 0.406 | 0.407 | 0.378 | 0.306 | 0.160 | 0.537 | 0.423 |
| Kelsterbacl | 0.452 | 0.493 | 0.483 | 0.439 | 0.430 | 0.432 | 0.425 | 0.361 | 0.266 | 0.223 | 0.636 | 0.497 |
| Kil-0 | 0.422 | 0.464 | 0.460 | 0.432 | 0.416 | 0.403 | 0.410 | 0.367 | 0.279 | 0.215 | 0.606 | 0.485 |
| Kin-0 | 0.399 | 0.425 | 0.435 | 0.408 | 0.396 | 0.414 | 0.418 | 0.377 | 0.316 | 0.173 | 0.543 | 0.448 |
| Kl-0 | 0.422 | 0.465 | 0.472 | 0.441 | 0.419 | 0.401 | 0.407 | 0.388 | 0.302 | 0.210 | 0.595 | 0.486 |
| Kl-5 | 0.404 | 0.442 | 0.460 | 0.445 | 0.409 | 0.388 | 0.397 | 0.362 | 0.297 | 0.235 | 0.637 | 0.507 |
| Kn-0 | 0.454 | 0.482 | 0.465 | 0.430 | 0.414 | 0.415 | 0.420 | 0.377 | 0.314 | 0.179 | 0.558 | 0.445 |
| KNO-11 | 0.419 | 0.511 | 0.466 | 0.414 | 0.401 | 0.400 | 0.395 | 0.332 | 0.247 | 0.254 | 0.646 | 0.539 |
| Kno-18 | 0.439 | 0.460 | 0.447 | 0.410 | 0.406 | 0.406 | 0.377 | 0.305 | 0.243 | 0.234 | 0.647 | 0.503 |
| Koln | 0.422 | 0.459 | 0.458 | 0.426 | 0.414 | 0.417 | 0.416 | 0.369 | 0.286 | 0.204 | 0.591 | 0.480 |
| Kondara | 0.416 | 0.442 | 0.440 | 0.410 | 0.389 | 0.401 | 0.399 | 0.334 | 0.260 | 0.199 | 0.593 | 0.466 |
| Kr-0 | 0.427 | 0.484 | 0.485 | 0.443 | 0.407 | 0.384 | 0.387 | 0.374 | 0.280 | 0.227 | 0.593 | 0.525 |
| Kro-0 | 0.446 | 0.523 | 0.498 | 0.443 | 0.418 | 0.414 | 0.413 | 0.396 | 0.326 | 0.160 | 0.533 | 0.421 |
| Krot-2 | 0.406 | 0.445 | 0.441 | 0.419 | 0.410 | 0.412 | 0.417 | 0.399 | 0.355 | 0.179 | 0.564 | 0.434 |
| Kz-1 | 0.450 | 0.464 | 0.432 | 0.378 | 0.366 | 0.374 | 0.407 | 0.394 | 0.303 | 0.185 | 0.591 | 0.432 |
| Kz-9 | 0.428 | 0.453 | 0.445 | 0.419 | 0.406 | 0.400 | 0.404 | 0.368 | 0.277 | 0.177 | 0.553 | 0.443 |
| LAC-3 | 0.406 | 0.449 | 0.456 | 0.430 | 0.401 | 0.400 | 0.405 | 0.349 | 0.257 | 0.180 | 0.564 | 0.443 |
| LAC-5 | 0.381 | 0.416 | 0.422 | 0.413 | 0.409 | 0.393 | 0.394 | 0.395 | 0.355 | 0.182 | 0.569 | 0.445 |
| Lan-0 | 0.417 | 0.463 | 0.450 | 0.424 | 0.409 | 0.404 | 0.397 | 0.350 | 0.284 | 0.197 | 0.610 | 0.449 |
| Laud-1 | 0.433 | 0.441 | 0.434 | 0.411 | 0.408 | 0.419 | 0.419 | 0.375 | 0.302 | 0.172 | 0.552 | 0.434 |
| Lc-0 | 0.383 | 0.433 | 0.465 | 0.435 | 0.423 | 0.411 | 0.403 | 0.391 | 0.349 | 0.154 | 0.541 | 0.402 |
| LDV-14 | 0.358 | 0.403 | 0.442 | 0.436 | 0.408 | 0.384 | 0.405 | 0.393 | 0.349 | 0.212 | 0.606 | 0.466 |
| LDV-25 | 0.389 | 0.432 | 0.438 | 0.416 | 0.407 | 0.408 | 0.401 | 0.354 | 0.281 | 0.203 | 0.588 | 0.481 |
| LDV-34 | 0.400 | 0.433 | 0.438 | 0.411 | 0.402 | 0.396 | 0.392 | 0.363 | 0.312 | 0.183 | 0.576 | 0.446 |
| LDV-58 | 0.387 | 0.415 | 0.425 | 0.407 | 0.400 | 0.406 | 0.403 | 0.363 | 0.303 | 0.161 | 0.543 | 0.416 |
| Ler-1 | 0.414 | 0.447 | 0.449 | 0.418 | 0.394 | 0.369 | 0.367 | 0.365 | 0.334 | 0.163 | 0.534 | 0.431 |
| Li-3 | 0.428 | 0.469 | 0.458 | 0.429 | 0.409 | 0.402 | 0.401 | 0.354 | 0.257 | 0.149 | 0.538 | 0.395 |
| Li-5:2 | 0.355 | 0.399 | 0.422 | 0.393 | 0.358 | 0.328 | 0.308 | 0.310 | 0.277 | 0.228 | 0.676 | 0.470 |
| Li-6 | 0.410 | 0.460 | 0.455 | 0.425 | 0.401 | 0.400 | 0.407 | 0.371 | 0.296 | 0.177 | 0.565 | 0.445 |

| accession | RGR08_10 | RGR09_11 | RGR10_12 | RGR11_13 | RGR12_14 | RGR13_15 | RGR14_16 | RGR15_17 | RGR16_18 | SA | SL | SW |
|---|---|---|---|---|---|---|---|---|---|---|---|---|
| Li-7 | 0.410 | 0.441 | 0.438 | 0.424 | 0.414 | 0.392 | 0.375 | 0.362 | 0.300 | 0.176 | 0.558 | 0.440 |
| Liarum | 0.414 | 0.441 | 0.448 | 0.417 | 0.400 | 0.390 | 0.389 | 0.357 | 0.292 | 0.171 | 0.565 | 0.428 |
| Limeport | 0.426 | 0.467 | 0.453 | 0.416 | 0.407 | 0.402 | 0.405 | 0.374 | 0.314 | 0.158 | 0.571 | 0.390 |
| Lip-0 | 0.451 | 0.466 | 0.446 | 0.415 | 0.410 | 0.411 | 0.396 | 0.342 | 0.271 | 0.159 | 0.558 | 0.394 |
| Lis-1 | 0.437 | 0.460 | 0.466 | 0.440 | 0.416 | 0.404 | 0.405 | 0.372 | 0.323 | 0.237 | 0.642 | 0.519 |
| LL-0 | 0.340 | 0.336 | 0.316 | 0.274 | 0.227 | 0.248 | 0.321 | 0.369 | 0.413 | 0.154 | 0.514 | 0.425 |
| Ll-OF-095 | 0.392 | 0.405 | 0.410 | 0.412 | 0.396 | 0.378 | 0.389 | 0.378 | 0.310 | 0.207 | 0.594 | 0.492 |
| Lm | 0.392 | 0.467 | 0.479 | 0.440 | 0.413 | 0.429 | 0.420 | 0.361 | 0.313 | 0.166 | 0.546 | 0.420 |
| Lm-2 | 0.396 | 0.473 | 0.484 | 0.439 | 0.410 | 0.427 | 0.427 | 0.372 | 0.313 | 0.187 | 0.584 | 0.448 |
| Lom1-1 | 0.405 | 0.444 | 0.450 | 0.420 | 0.397 | 0.390 | 0.375 | 0.313 | 0.228 | 0.168 | 0.560 | 0.423 |
| Löv-5 | 0.388 | 0.469 | 0.465 | 0.422 | 0.388 | 0.374 | 0.369 | 0.341 | 0.290 | 0.204 | 0.645 | 0.437 |
| Lp2-2 | 0.437 | 0.457 | 0.452 | 0.416 | 0.415 | 0.437 | 0.414 | 0.337 | 0.273 | 0.207 | 0.611 | 0.477 |
| Lp2-6 | 0.435 | 0.476 | 0.477 | 0.436 | 0.419 | 0.422 | 0.429 | 0.388 | 0.303 | 0.163 | 0.509 | 0.421 |
| Lu | 0.421 | 0.448 | 0.442 | 0.414 | 0.399 | 0.381 | 0.386 | 0.387 | 0.342 | 0.218 | 0.535 | 0.470 |
| Lz-0 | 0.427 | 0.471 | 0.464 | 0.426 | 0.402 | 0.386 | 0.358 | 0.291 | 0.215 | 0.190 | 0.579 | 0.435 |
| Map-42 | 0.401 | 0.422 | 0.426 | 0.406 | 0.388 | 0.381 | 0.389 | 0.368 | 0.295 | 0.184 | 0.518 | 0.350 |
| Mc-0 | 0.345 | 0.385 | 0.397 | 0.376 | 0.376 | 0.385 | 0.394 | 0.406 | 0.355 | 0.207 | 0.448 | 0.398 |
| Me-0 | 0.408 | 0.438 | 0.447 | 0.405 | 0.388 | 0.402 | 0.401 | 0.364 | 0.309 | 0.201 | 0.576 | 0.482 |
| Mh-0 | 0.406 | 0.424 | 0.418 | 0.376 | 0.371 | 0.387 | 0.382 | 0.357 | 0.304 | 0.212 | 0.594 | 0.466 |
| Mh-1 | 0.404 | 0.441 | 0.431 | 0.410 | 0.412 | 0.405 | 0.395 | 0.347 | 0.258 | 0.193 | 0.548 | 0.434 |
| MIB-15 | 0.399 | 0.461 | 0.442 | 0.406 | 0.395 | 0.388 | 0.386 | 0.365 | 0.299 | 0.202 | 0.506 | 0.392 |
| MIB-22 | 0.381 | 0.468 | 0.504 | 0.438 | 0.415 | 0.406 | 0.402 | 0.377 | 0.319 | 0.145 | 0.524 | 0.391 |
| MIB-28 | 0.422 | 0.455 | 0.448 | 0.424 | 0.411 | 0.406 | 0.408 | 0.373 | 0.328 | 0.159 | 0.548 | 0.404 |
| MIB-84 | 0.415 | 0.463 | 0.468 | 0.432 | 0.409 | 0.402 | 0.405 | 0.397 | 0.355 | 0.168 | 0.487 | 0.366 |
| MNF-Pot- | 0.423 | 0.451 | 0.463 | 0.434 | 0.407 | 0.404 | 0.420 | 0.398 | 0.343 | 0.148 | 0.499 | 0.380 |
| Mnz-0 | 0.409 | 0.434 | 0.442 | 0.410 | 0.388 | 0.397 | 0.405 | 0.366 | 0.288 | 0.197 | 0.566 | 0.457 |
| MOG-37 | 0.380 | 0.433 | 0.456 | 0.440 | 0.421 | 0.406 | 0.414 | 0.401 | 0.333 | 0.177 | 0.574 | 0.423 |
| Mrk-0 | 0.417 | 0.446 | 0.443 | 0.415 | 0.400 | 0.405 | 0.407 | 0.370 | 0.306 | 0.179 | 0.577 | 0.400 |
| Ms-0 | 0.427 | 0.482 | 0.501 | 0.476 | 0.428 | 0.401 | 0.395 | 0.353 | 0.289 | 0.210 | 0.591 | 0.460 |
| Mt-0 | 0.446 | 0.464 | 0.455 | 0.428 | 0.414 | 0.421 | 0.419 | 0.363 | 0.288 | 0.214 | 0.578 | 0.434 |

| accession | RGR08_10 | RGR09_11 | RGR10_12 | RGR11_13 | RGR12_14 | RGR13_15 | RGR14_16 | RGR15_17 | RGR16_18 | SA | SL | SW |
|---|---|---|---|---|---|---|---|---|---|---|---|---|
| Mz-0 | 0.409 | 0.429 | 0.418 | 0.377 | 0.344 | 0.341 | 0.352 | 0.331 | 0.287 | 0.230 | 0.579 | 0.473 |
| N13 | 0.431 | 0.480 | 0.497 | 0.471 | 0.436 | 0.406 | 0.410 | 0.390 | 0.337 | 0.169 | 0.534 | 0.422 |
| N4 | 0.424 | 0.457 | 0.463 | 0.441 | 0.407 | 0.378 | 0.375 | 0.342 | 0.264 | 0.206 | 0.603 | 0.461 |
| N7 | 0.408 | 0.435 | 0.453 | 0.445 | 0.422 | 0.400 | 0.391 | 0.345 | 0.261 | 0.177 | 0.546 | 0.437 |
| Na-1 | 0.391 | 0.490 | 0.504 | 0.442 | 0.417 | 0.405 | 0.401 | 0.387 | 0.337 | 0.172 | 0.524 | 0.425 |
| Nc-1 | 0.390 | 0.483 | 0.485 | 0.417 | 0.385 | 0.387 | 0.388 | 0.359 | 0.279 | 0.258 | 0.672 | 0.521 |
| Nd | 0.409 | 0.442 | 0.458 | 0.430 | 0.400 | 0.387 | 0.406 | 0.405 | 0.343 | 0.225 | 0.625 | 0.498 |
| Nd-1 | 0.415 | 0.466 | 0.456 | 0.422 | 0.397 | 0.395 | 0.408 | 0.389 | 0.333 | 0.242 | 0.643 | 0.524 |
| NFA-10 | 0.400 | 0.427 | 0.428 | 0.408 | 0.398 | 0.409 | 0.396 | 0.349 | 0.275 | 0.214 | 0.613 | 0.487 |
| NFA-8 | 0.436 | 0.478 | 0.448 | 0.418 | 0.403 | 0.400 | 0.370 | 0.281 | 0.195 | 0.243 | 0.635 | 0.515 |
| NFC-20 | 0.401 | 0.419 | 0.419 | 0.409 | 0.392 | 0.390 | 0.391 | 0.352 | 0.290 | 0.234 | 0.644 | 0.493 |
| No-0 | 0.410 | 0.463 | 0.482 | 0.450 | 0.413 | 0.395 | 0.379 | 0.315 | 0.225 | 0.234 | 0.647 | 0.524 |
| Nok-1 | 0.412 | 0.439 | 0.430 | 0.399 | 0.395 | 0.372 | 0.359 | 0.349 | 0.288 | 0.309 | 0.737 | 0.572 |
| Nok-2 | 0.422 | 0.452 | 0.454 | 0.435 | 0.417 | 0.395 | 0.394 | 0.362 | 0.280 | 0.261 | 0.657 | 0.554 |
| Nw-0 | 0.434 | 0.471 | 0.461 | 0.414 | 0.397 | 0.408 | 0.421 | 0.385 | 0.336 | 0.183 | 0.555 | 0.459 |
| Nw-2 | 0.429 | 0.454 | 0.466 | 0.447 | 0.436 | 0.428 | 0.420 | 0.380 | 0.296 | 0.169 | 0.549 | 0.421 |
| NW3 | 0.409 | 0.455 | 0.441 | 0.420 | 0.405 | 0.401 | 0.387 | 0.335 | 0.251 | 0.243 | 0.657 | 0.505 |
| Nz1 | 0.416 | 0.462 | 0.451 | 0.424 | 0.413 | 0.397 | 0.382 | 0.344 | 0.284 | 0.212 | 0.600 | 0.472 |
| Ob-1 | 0.436 | 0.458 | 0.449 | 0.418 | 0.409 | 0.414 | 0.420 | 0.405 | 0.324 | 0.171 | 0.530 | 0.450 |
| Old-1 | 0.432 | 0.459 | 0.467 | 0.439 | 0.418 | 0.397 | 0.383 | 0.361 | 0.292 | 0.218 | 0.613 | 0.494 |
| Or-0 | 0.397 | 0.429 | 0.434 | 0.405 | 0.398 | 0.417 | 0.435 | 0.403 | 0.342 | 0.204 | 0.574 | 0.489 |
| Ors-1 | 0.381 | 0.449 | 0.461 | 0.421 | 0.399 | 0.389 | 0.391 | 0.364 | 0.295 | 0.251 | 0.658 | 0.519 |
| Ors-2 | 0.383 | 0.435 | 0.447 | 0.414 | 0.387 | 0.374 | 0.383 | 0.371 | 0.312 | 0.225 | 0.630 | 0.503 |
| Ost-0 | 0.343 | 0.424 | 0.443 | 0.400 | 0.356 | 0.333 | 0.337 | 0.334 | 0.299 | 0.301 | 0.728 | 0.565 |
| Ove-0 | 0.406 | 0.435 | 0.437 | 0.411 | 0.390 | 0.378 | 0.367 | 0.325 | 0.267 | 0.242 | 0.642 | 0.527 |
| Ove-0 | 0.398 | 0.428 | 0.433 | 0.410 | 0.389 | 0.387 | 0.390 | 0.359 | 0.289 | 0.225 | 0.621 | 0.500 |
| Oy-0 | 0.429 | 0.452 | 0.462 | 0.449 | 0.430 | 0.422 | 0.413 | 0.365 | 0.307 | 0.230 | 0.627 | 0.499 |
| Oy-1 | 0.421 | 0.445 | 0.441 | 0.426 | 0.408 | 0.413 | 0.423 | 0.380 | 0.301 | 0.199 | 0.600 | 0.451 |
| Pa-1 | 0.395 | 0.425 | 0.430 | 0.403 | 0.394 | 0.405 | 0.403 | 0.358 | 0.296 | 0.208 | 0.606 | 0.480 |
| Pa-2 | 0.451 | 0.464 | 0.434 | 0.390 | 0.380 | 0.392 | 0.388 | 0.337 | 0.260 | 0.180 | 0.557 | 0.449 |

| accession | RGR08_10 | RGR09_11 | RGR10_12 | RGR11_13 | RGR12_14 | RGR13_15 | RGR14_16 | RGR15_17 | RGR16_18 | SA | SL | SW |
|---|---|---|---|---|---|---|---|---|---|---|---|---|
| PAR-3 | 0.404 | 0.415 | 0.397 | 0.369 | 0.371 | 0.373 | 0.374 | 0.356 | 0.281 | 0.203 | 0.620 | 0.454 |
| PAR-4 | 0.389 | 0.430 | 0.424 | 0.402 | 0.386 | 0.387 | 0.410 | 0.395 | 0.331 | 0.218 | 0.621 | 0.481 |
| Per-1 | 0.415 | 0.452 | 0.455 | 0.428 | 0.406 | 0.399 | 0.418 | 0.404 | 0.340 | 0.190 | 0.595 | 0.446 |
| Petergof | 0.369 | 0.464 | 0.463 | 0.399 | 0.391 | 0.394 | 0.399 | 0.388 | 0.347 | 0.161 | 0.550 | 0.409 |
| PHW-10 | 0.385 | 0.425 | 0.443 | 0.423 | 0.398 | 0.384 | 0.364 | 0.326 | 0.247 | 0.190 | 0.585 | 0.452 |
| PHW-13 | 0.388 | 0.419 | 0.423 | 0.400 | 0.388 | 0.394 | 0.415 | 0.394 | 0.329 | 0.185 | 0.565 | 0.459 |
| PHW-14 | 0.401 | 0.427 | 0.433 | 0.403 | 0.394 | 0.389 | 0.385 | 0.376 | 0.331 | 0.191 | 0.571 | 0.463 |
| PHW-20 | 0.390 | 0.418 | 0.422 | 0.401 | 0.382 | 0.375 | 0.377 | 0.350 | 0.269 | 0.175 | 0.560 | 0.442 |
| PHW-22 | 0.389 | 0.426 | 0.432 | 0.412 | 0.414 | 0.426 | 0.418 | 0.381 | 0.306 | 0.191 | 0.581 | 0.455 |
| PHW-26 | 0.411 | 0.460 | 0.466 | 0.437 | 0.401 | 0.375 | 0.368 | 0.319 | 0.232 | 0.190 | 0.609 | 0.437 |
| PHW-28 | 0.412 | 0.421 | 0.419 | 0.398 | 0.379 | 0.377 | 0.386 | 0.356 | 0.243 | 0.203 | 0.601 | 0.469 |
| PHW-31 | 0.358 | 0.408 | 0.423 | 0.401 | 0.381 | 0.366 | 0.375 | 0.390 | 0.374 | 0.192 | 0.585 | 0.454 |
| PHW-33 | 0.405 | 0.433 | 0.430 | 0.399 | 0.375 | 0.370 | 0.350 | 0.300 | 0.267 | 0.210 | 0.621 | 0.464 |
| PHW-35 | 0.410 | 0.453 | 0.419 | 0.356 | 0.312 | 0.310 | 0.309 | 0.289 | 0.286 | 0.196 | 0.603 | 0.456 |
| PHW-36 | 0.409 | 0.441 | 0.448 | 0.424 | 0.407 | 0.406 | 0.406 | 0.347 | 0.258 | 0.189 | 0.579 | 0.457 |
| PHW-37 | 0.422 | 0.455 | 0.452 | 0.424 | 0.390 | 0.381 | 0.401 | 0.375 | 0.309 | 0.157 | 0.548 | 0.409 |
| Pi-0 | 0.382 | 0.404 | 0.415 | 0.405 | 0.389 | 0.383 | 0.396 | 0.369 | 0.303 | 0.198 | 0.591 | 0.465 |
| Pla-0 | 0.400 | 0.433 | 0.437 | 0.410 | 0.393 | 0.395 | 0.390 | 0.323 | 0.239 | 0.218 | 0.624 | 0.482 |
| Pn-0 | 0.406 | 0.420 | 0.422 | 0.414 | 0.400 | 0.397 | 0.400 | 0.356 | 0.285 | 0.202 | 0.612 | 0.466 |
| Pna-10 | 0.378 | 0.405 | 0.412 | 0.400 | 0.386 | 0.388 | 0.391 | 0.357 | 0.296 | 0.182 | 0.584 | 0.437 |
| Pna-17 | 0.370 | 0.521 | 0.495 | 0.407 | 0.383 | 0.381 | 0.396 | 0.374 | 0.300 | 0.167 | 0.557 | 0.417 |
| Pna-17 | 0.405 | 0.557 | 0.466 | 0.408 | 0.389 | 0.374 | 0.391 | 0.377 | 0.314 | 0.163 | 0.552 | 0.416 |
| Po-0 | 0.416 | 0.453 | 0.450 | 0.419 | 0.400 | 0.400 | 0.404 | 0.353 | 0.276 | 0.186 | 0.580 | 0.444 |
| Pog-0 | 0.403 | 0.441 | 0.413 | 0.387 | 0.386 | 0.411 | 0.405 | 0.350 | 0.289 | 0.184 | 0.600 | 0.428 |
| Pr-0 | 0.463 | 0.483 | 0.453 | 0.419 | 0.408 | 0.417 | 0.399 | 0.337 | 0.282 | 0.148 | 0.529 | 0.391 |
| Pro-0 | 0.411 | 0.432 | 0.417 | 0.388 | 0.378 | 0.381 | 0.405 | 0.385 | 0.312 | 0.195 | 0.569 | 0.474 |
| Pt-0 | 0.408 | 0.462 | 0.468 | 0.430 | 0.425 | 0.427 | 0.421 | 0.378 | 0.314 | 0.165 | 0.560 | 0.414 |
| Pu2-23 | 0.439 | 0.499 | 0.482 | 0.419 | 0.403 | 0.409 | 0.383 | 0.327 | 0.294 | 0.146 | 0.546 | 0.380 |
| PU2-24 | 0.429 | 0.492 | 0.498 | 0.460 | 0.430 | 0.415 | 0.384 | 0.311 | 0.230 | 0.182 | 0.581 | 0.441 |
| Pu2-7 | 0.416 | 0.461 | 0.462 | 0.434 | 0.412 | 0.412 | 0.418 | 0.375 | 0.296 | 0.173 | 0.565 | 0.424 |

| accession | RGR08_10 | RGR09_11 | RGR10_12 | RGR11_13 | RGR12_14 | RGR13_15 | RGR14_16 | RGR15_17 | RGR16_18 | SA | SL | SW |
|---|---|---|---|---|---|---|---|---|---|---|---|---|
| Pyl-1 | 0.376 | 0.392 | 0.412 | 0.396 | 0.378 | 0.352 | 0.351 | 0.350 | 0.290 | 0.231 | 0.676 | 0.472 |
| Ra-0 | 0.399 | 0.419 | 0.433 | 0.417 | 0.403 | 0.399 | 0.401 | 0.382 | 0.309 | 0.149 | 0.520 | 0.403 |
| Rak-2 | 0.403 | 0.452 | 0.440 | 0.419 | 0.401 | 0.394 | 0.391 | 0.365 | 0.314 | 0.178 | 0.553 | 0.448 |
| Ren-1 | 0.405 | 0.425 | 0.429 | 0.409 | 0.410 | 0.403 | 0.391 | 0.370 | 0.319 | 0.157 | 0.533 | 0.419 |
| Ren-11 | 0.381 | 0.438 | 0.444 | 0.407 | 0.386 | 0.406 | 0.435 | 0.419 | 0.391 | 0.148 | 0.530 | 0.395 |
| Rhen-1 | 0.383 | 0.436 | 0.437 | 0.403 | 0.381 | 0.389 | 0.397 | 0.362 | 0.312 | 0.185 | 0.586 | 0.442 |
| Ri-0 | 0.397 | 0.428 | 0.433 | 0.406 | 0.404 | 0.415 | 0.400 | 0.352 | 0.298 | 0.152 | 0.526 | 0.414 |
| RLD-1 | 0.438 | 0.487 | 0.494 | 0.447 | 0.393 | 0.396 | 0.433 | 0.395 | 0.296 | 0.141 | 0.536 | 0.378 |
| RLD-2 | 0.453 | 0.494 | 0.489 | 0.446 | 0.400 | 0.390 | 0.410 | 0.378 | 0.299 | 0.159 | 0.556 | 0.400 |
| Rmx-A02 | 0.433 | 0.448 | 0.442 | 0.429 | 0.405 | 0.407 | 0.406 | 0.343 | 0.272 | 0.187 | 0.604 | 0.433 |
| Rmx-A18( | 0.419 | 0.470 | 0.462 | 0.434 | 0.417 | 0.417 | 0.414 | 0.371 | 0.287 | 0.178 | 0.557 | 0.449 |
| Rou-0 | 0.396 | 0.439 | 0.431 | 0.400 | 0.384 | 0.384 | 0.387 | 0.345 | 0.267 | 0.150 | 0.515 | 0.411 |
| RRS-10 | 0.373 | 0.429 | 0.449 | 0.411 | 0.389 | 0.385 | 0.401 | 0.388 | 0.313 | 0.151 | 0.535 | 0.402 |
| RRS-7 | 0.371 | 0.410 | 0.432 | 0.412 | 0.392 | 0.385 | 0.389 | 0.369 | 0.312 | 0.146 | 0.509 | 0.404 |
| Rsch-0 | 0.430 | 0.476 | 0.472 | 0.425 | 0.382 | 0.379 | 0.407 | 0.388 | 0.328 | 0.138 | 0.520 | 0.378 |
| Rsch-4 | 0.443 | 0.481 | 0.473 | 0.440 | 0.419 | 0.418 | 0.411 | 0.350 | 0.246 | 0.154 | 0.533 | 0.408 |
| Rubeszhno | 0.413 | 0.458 | 0.467 | 0.440 | 0.396 | 0.392 | 0.412 | 0.367 | 0.280 | 0.141 | 0.518 | 0.377 |
| S96 | 0.418 | 0.453 | 0.450 | 0.421 | 0.415 | 0.403 | 0.400 | 0.373 | 0.327 | 0.191 | 0.578 | 0.452 |
| Santa Clara | 0.379 | 0.408 | 0.415 | 0.422 | 0.428 | 0.396 | 0.343 | 0.343 | 0.371 | 0.166 | 0.554 | 0.424 |
| Sap-0 | 0.422 | 0.464 | 0.470 | 0.444 | 0.432 | 0.420 | 0.410 | 0.375 | 0.295 | 0.193 | 0.587 | 0.457 |
| Sapporo-0 | 0.409 | 0.445 | 0.446 | 0.415 | 0.378 | 0.387 | 0.405 | 0.354 | 0.285 | 0.172 | 0.546 | 0.442 |
| Sav-0 | 0.400 | 0.424 | 0.429 | 0.422 | 0.409 | 0.401 | 0.392 | 0.357 | 0.270 | 0.193 | 0.595 | 0.455 |
| Sav-0 | 0.427 | 0.450 | 0.459 | 0.434 | 0.413 | 0.396 | 0.389 | 0.376 | 0.315 | 0.191 | 0.598 | 0.434 |
| Se-0 | 0.392 | 0.443 | 0.447 | 0.406 | 0.385 | 0.384 | 0.381 | 0.328 | 0.259 | 0.212 | 0.613 | 0.477 |
| Sei-0 | 0.412 | 0.440 | 0.463 | 0.435 | 0.420 | 0.414 | 0.420 | 0.391 | 0.322 | 0.178 | 0.576 | 0.436 |
| Sg-1 | 0.418 | 0.449 | 0.437 | 0.406 | 0.394 | 0.388 | 0.389 | 0.364 | 0.296 | 0.191 | 0.609 | 0.436 |
| Sh-0 | 0.428 | 0.458 | 0.449 | 0.418 | 0.399 | 0.400 | 0.415 | 0.390 | 0.327 | 0.176 | 0.560 | 0.440 |
| Shahdara | 0.388 | 0.420 | 0.432 | 0.413 | 0.389 | 0.376 | 0.391 | 0.377 | 0.311 | 0.205 | 0.615 | 0.464 |
| Si-0 | 0.438 | 0.474 | 0.471 | 0.440 | 0.423 | 0.412 | 0.406 | 0.373 | 0.327 | 0.152 | 0.546 | 0.394 |
| SLSP-30 | 0.417 | 0.444 | 0.443 | 0.417 | 0.399 | 0.400 | 0.404 | 0.360 | 0.273 | 0.171 | 0.559 | 0.424 |

| accession | RGR08_10 | RGR09_11 | RGR10_12 | RGR11_13 | RGR12_14 | RGR13_15 | RGR14_16 | RGR15_17 | RGR16_18 | SA | SL | SW |
|---|---|---|---|---|---|---|---|---|---|---|---|---|
| Sorbo | 0.380 | 0.438 | 0.441 | 0.405 | 0.381 | 0.375 | 0.367 | 0.313 | 0.225 | 0.210 | 0.625 | 0.466 |
| Sp-0 | 0.438 | 0.492 | 0.506 | 0.468 | 0.439 | 0.422 | 0.434 | 0.412 | 0.337 | 0.177 | 0.564 | 0.438 |
| Sq-1 | 0.388 | 0.452 | 0.464 | 0.414 | 0.387 | 0.390 | 0.414 | 0.378 | 0.294 | 0.188 | 0.570 | 0.458 |
| Sq-8 | 0.421 | 0.465 | 0.460 | 0.430 | 0.417 | 0.413 | 0.410 | 0.364 | 0.276 | 0.201 | 0.606 | 0.457 |
| St-0 | 0.434 | 0.470 | 0.453 | 0.427 | 0.417 | 0.404 | 0.413 | 0.381 | 0.286 | 0.175 | 0.557 | 0.441 |
| Ste-0 | 0.432 | 0.509 | 0.482 | 0.436 | 0.413 | 0.398 | 0.396 | 0.357 | 0.284 | 0.191 | 0.574 | 0.456 |
| Ste-3 | 0.386 | 0.503 | 0.517 | 0.458 | 0.427 | 0.400 | 0.405 | 0.406 | 0.369 | 0.171 | 0.577 | 0.415 |
| Stw-0 | 0.418 | 0.450 | 0.446 | 0.419 | 0.409 | 0.403 | 0.390 | 0.345 | 0.270 | 0.186 | 0.585 | 0.437 |
| Ta-0 | 0.411 | 0.433 | 0.433 | 0.419 | 0.406 | 0.383 | 0.367 | 0.340 | 0.271 | 0.224 | 0.631 | 0.489 |
| TAMM-2 | 0.322 | 0.423 | 0.463 | 0.433 | 0.399 | 0.376 | 0.407 | 0.386 | 0.292 | 0.200 | 0.614 | 0.451 |
| Tamm-27 | 0.324 | 0.453 | 0.449 | 0.403 | 0.384 | 0.396 | 0.396 | 0.350 | 0.265 | 0.223 | 0.643 | 0.477 |
| TDr-1 | 0.435 | 0.492 | 0.481 | 0.447 | 0.420 | 0.405 | 0.394 | 0.340 | 0.283 | 0.220 | 0.648 | 0.472 |
| TDr-3 | 0.423 | 0.467 | 0.459 | 0.430 | 0.414 | 0.417 | 0.420 | 0.380 | 0.317 | 0.199 | 0.596 | 0.457 |
| Te-0 | 0.416 | 0.484 | 0.482 | 0.439 | 0.407 | 0.386 | 0.381 | 0.331 | 0.251 | 0.279 | 0.703 | 0.546 |
| Tha-1 | 0.429 | 0.453 | 0.441 | 0.411 | 0.399 | 0.393 | 0.408 | 0.396 | 0.312 | 0.192 | 0.610 | 0.438 |
| Ting-1 | 0.411 | 0.478 | 0.452 | 0.420 | 0.399 | 0.389 | 0.386 | 0.362 | 0.295 | 0.221 | 0.611 | 0.502 |
| Tiv-1 | 0.391 | 0.457 | 0.465 | 0.423 | 0.404 | 0.389 | 0.377 | 0.350 | 0.296 | 0.225 | 0.646 | 0.478 |
| Tol-0 | 0.389 | 0.441 | 0.430 | 0.412 | 0.394 | 0.377 | 0.382 | 0.345 | 0.263 | 0.235 | 0.617 | 0.521 |
| Tottarp-2 | 0.441 | 0.466 | 0.455 | 0.424 | 0.405 | 0.407 | 0.412 | 0.382 | 0.318 | 0.188 | 0.566 | 0.467 |
| TOU-A1-1 | 0.404 | 0.439 | 0.448 | 0.425 | 0.405 | 0.397 | 0.386 | 0.359 | 0.318 | 0.232 | 0.636 | 0.501 |
| TOU-A1-1 | 0.417 | 0.438 | 0.433 | 0.417 | 0.403 | 0.395 | 0.395 | 0.385 | 0.352 | 0.212 | 0.603 | 0.491 |
| TOU-A1-4 | 0.430 | 0.455 | 0.451 | 0.422 | 0.401 | 0.395 | 0.390 | 0.339 | 0.268 | 0.228 | 0.662 | 0.475 |
| TOU-A1-6 | 0.394 | 0.427 | 0.427 | 0.401 | 0.381 | 0.383 | 0.402 | 0.385 | 0.314 | 0.212 | 0.640 | 0.461 |
| TOU-A1-9 | 0.409 | 0.440 | 0.434 | 0.397 | 0.375 | 0.374 | 0.387 | 0.371 | 0.311 | 0.215 | 0.649 | 0.472 |
| TOU-C-3 | 0.388 | 0.415 | 0.416 | 0.401 | 0.385 | 0.388 | 0.390 | 0.353 | 0.291 | 0.227 | 0.614 | 0.510 |
| TOU-E-11 | 0.426 | 0.460 | 0.465 | 0.440 | 0.402 | 0.386 | 0.386 | 0.335 | 0.253 | 0.191 | 0.604 | 0.445 |
| TOU-H-13 | 0.401 | 0.432 | 0.438 | 0.416 | 0.408 | 0.402 | 0.410 | 0.380 | 0.308 | 0.202 | 0.612 | 0.459 |
| TOU-I-17 | 0.427 | 0.449 | 0.437 | 0.409 | 0.394 | 0.392 | 0.396 | 0.371 | 0.310 | 0.158 | 0.555 | 0.401 |
| TOU-I-2 | 0.418 | 0.442 | 0.421 | 0.379 | 0.355 | 0.362 | 0.361 | 0.328 | 0.291 | 0.228 | 0.640 | 0.498 |
| TOU-I-6 | 0.463 | 0.484 | 0.465 | 0.435 | 0.413 | 0.405 | 0.387 | 0.328 | 0.245 | 0.226 | 0.641 | 0.481 |

| accession | RGR08_10 | RGR09_11 | RGR10_12 | RGR11_13 | RGR12_14 | RGR13_15 | RGR14_16 | RGR15_17 | RGR16_18 | SA | SL | SW |
|---|---|---|---|---|---|---|---|---|---|---|---|---|
| Ts-1 | 0.408 | 0.449 | 0.435 | 0.399 | 0.380 | 0.370 | 0.385 | 0.382 | 0.309 | 0.222 | 0.638 | 0.490 |
| Ts-5 | 0.357 | 0.459 | 0.458 | 0.414 | 0.404 | 0.389 | 0.401 | 0.377 | 0.292 | 0.236 | 0.692 | 0.474 |
| Tscha-1 | 0.408 | 0.451 | 0.450 | 0.414 | 0.403 | 0.392 | 0.378 | 0.363 | 0.325 | 0.189 | 0.603 | 0.439 |
| Tsu-0 | 0.454 | 0.481 | 0.466 | 0.429 | 0.409 | 0.415 | 0.393 | 0.321 | 0.243 | 0.193 | 0.591 | 0.453 |
| Tsu-1 | 0.443 | 0.501 | 0.446 | 0.416 | 0.409 | 0.410 | 0.379 | 0.308 | 0.223 | 0.225 | 0.630 | 0.491 |
| Tu-0 | 0.458 | 0.493 | 0.454 | 0.424 | 0.404 | 0.401 | 0.392 | 0.335 | 0.244 | 0.198 | 0.620 | 0.449 |
| Tul-0 | 0.404 | 0.434 | 0.433 | 0.414 | 0.390 | 0.388 | 0.385 | 0.341 | 0.257 | 0.207 | 0.613 | 0.471 |
| Ty-0 | 0.392 | 0.430 | 0.435 | 0.412 | 0.389 | 0.380 | 0.382 | 0.348 | 0.289 | 0.216 | 0.636 | 0.465 |
| Udul1-34 | 0.418 | 0.475 | 0.473 | 0.434 | 0.408 | 0.405 | 0.408 | 0.388 | 0.359 | 0.180 | 0.580 | 0.435 |
| Uk-1 | 0.430 | 0.469 | 0.458 | 0.420 | 0.408 | 0.421 | 0.436 | 0.403 | 0.334 | 0.184 | 0.598 | 0.427 |
| Uk-2 | 0.438 | 0.449 | 0.422 | 0.396 | 0.391 | 0.399 | 0.412 | 0.396 | 0.337 | 0.160 | 0.565 | 0.400 |
| Uk-4 | 0.461 | 0.473 | 0.449 | 0.416 | 0.414 | 0.399 | 0.410 | 0.407 | 0.340 | 0.174 | 0.576 | 0.424 |
| UKID48 | 0.401 | 0.443 | 0.435 | 0.391 | 0.375 | 0.372 | 0.352 | 0.327 | 0.275 | 0.188 | 0.595 | 0.447 |
| UKNW06- | 0.411 | 0.444 | 0.463 | 0.423 | 0.400 | 0.397 | 0.401 | 0.380 | 0.328 | 0.164 | 0.543 | 0.421 |
| UKNW06- | 0.410 | 0.450 | 0.441 | 0.418 | 0.399 | 0.398 | 0.404 | 0.376 | 0.324 | 0.171 | 0.540 | 0.442 |
| UKNW06- | 0.373 | 0.430 | 0.439 | 0.399 | 0.398 | 0.407 | 0.401 | 0.374 | 0.305 | 0.201 | 0.613 | 0.451 |
| UKSE06-4 | 0.378 | 0.421 | 0.417 | 0.388 | 0.385 | 0.398 | 0.411 | 0.403 | 0.339 | 0.164 | 0.551 | 0.416 |
| UKSE06-4 | 0.398 | 0.455 | 0.457 | 0.432 | 0.405 | 0.391 | 0.384 | 0.367 | 0.299 | 0.189 | 0.619 | 0.422 |
| UKSE06-4 | 0.415 | 0.442 | 0.429 | 0.397 | 0.408 | 0.428 | 0.406 | 0.361 | 0.300 | 0.184 | 0.576 | 0.444 |
| UKSE06-5 | 0.386 | 0.436 | 0.446 | 0.410 | 0.382 | 0.379 | 0.384 | 0.379 | 0.362 | 0.163 | 0.533 | 0.431 |
| UKSE06-6 | 0.397 | 0.432 | 0.436 | 0.406 | 0.403 | 0.416 | 0.403 | 0.358 | 0.293 | 0.196 | 0.583 | 0.465 |
| Ull2-3 | 0.432 | 0.460 | 0.447 | 0.426 | 0.414 | 0.413 | 0.416 | 0.374 | 0.286 | 0.167 | 0.545 | 0.426 |
| Ull-2-5 | 0.404 | 0.455 | 0.451 | 0.418 | 0.405 | 0.391 | 0.386 | 0.367 | 0.329 | 0.183 | 0.583 | 0.436 |
| Uod-1 | 0.427 | 0.508 | 0.461 | 0.414 | 0.406 | 0.421 | 0.415 | 0.361 | 0.289 | 0.139 | 0.515 | 0.377 |
| Uod-7 | 0.360 | 0.424 | 0.456 | 0.433 | 0.417 | 0.402 | 0.401 | 0.384 | 0.327 | 0.196 | 0.624 | 0.436 |
| Utrecht | 0.406 | 0.458 | 0.454 | 0.437 | 0.415 | 0.411 | 0.409 | 0.355 | 0.299 | 0.177 | 0.590 | 0.421 |
| Van-0 | 0.433 | 0.482 | 0.454 | 0.424 | 0.411 | 0.405 | 0.404 | 0.370 | 0.307 | 0.184 | 0.581 | 0.442 |
| Var-2-1 | 0.351 | 0.383 | 0.386 | 0.368 | 0.364 | 0.356 | 0.349 | 0.328 | 0.274 | 0.224 | 0.659 | 0.466 |
| Ven-1 | 0.388 | 0.433 | 0.437 | 0.413 | 0.392 | 0.372 | 0.362 | 0.315 | 0.254 | 0.221 | 0.639 | 0.479 |
| Wa-1 | 0.423 | 0.463 | 0.455 | 0.439 | 0.420 | 0.392 | 0.382 | 0.359 | 0.313 | 0.252 | 0.659 | 0.525 |

| accession | RGR08_10 | RGR09_11 | RGR10_12 | RGR11_13 | RGR12_14 | RGR13_15 | RGR14_16 | RGR15_17 | RGR16_18 | SA | SL | SW |
|---|---|---|---|---|---|---|---|---|---|---|---|---|
| Wa-1 | 0.418 | 0.452 | 0.451 | 0.431 | 0.406 | 0.390 | 0.385 | 0.345 | 0.258 | 0.252 | 0.672 | 0.516 |
| Wag-3 | 0.403 | 0.462 | 0.473 | 0.439 | 0.402 | 0.388 | 0.393 | 0.365 | 0.304 | 0.205 | 0.589 | 0.478 |
| Wag-4 | 0.400 | 0.464 | 0.470 | 0.436 | 0.407 | 0.410 | 0.394 | 0.324 | 0.238 | 0.253 | 0.659 | 0.523 |
| Wag-5 | 0.396 | 0.444 | 0.452 | 0.417 | 0.389 | 0.395 | 0.397 | 0.345 | 0.278 | 0.189 | 0.589 | 0.452 |
| WAR | 0.412 | 0.450 | 0.448 | 0.420 | 0.406 | 0.396 | 0.390 | 0.353 | 0.298 | 0.215 | 0.622 | 0.476 |
| Wc-2 | 0.448 | 0.476 | 0.471 | 0.450 | 0.439 | 0.419 | 0.392 | 0.349 | 0.266 | 0.198 | 0.582 | 0.468 |
| Wei-(1) | 0.418 | 0.438 | 0.440 | 0.410 | 0.394 | 0.395 | 0.356 | 0.265 | 0.201 | 0.246 | 0.642 | 0.521 |
| Wei-0 | 0.415 | 0.439 | 0.442 | 0.411 | 0.402 | 0.414 | 0.413 | 0.353 | 0.268 | 0.189 | 0.574 | 0.458 |
| Wil | 0.466 | 0.505 | 0.488 | 0.448 | 0.417 | 0.401 | 0.404 | 0.387 | 0.329 | 0.155 | 0.533 | 0.407 |
| Wil-2 | 0.457 | 0.516 | 0.477 | 0.435 | 0.421 | 0.420 | 0.419 | 0.389 | 0.325 | 0.167 | 0.550 | 0.419 |
| Wl-0 | 0.416 | 0.450 | 0.465 | 0.440 | 0.417 | 0.421 | 0.424 | 0.369 | 0.269 | 0.204 | 0.612 | 0.460 |
| Ws | 0.409 | 0.443 | 0.467 | 0.455 | 0.434 | 0.406 | 0.406 | 0.398 | 0.324 | 0.182 | 0.574 | 0.438 |
| Ws-0 | 0.430 | 0.478 | 0.468 | 0.435 | 0.428 | 0.423 | 0.385 | 0.320 | 0.286 | 0.196 | 0.615 | 0.438 |
| Ws-2 | 0.408 | 0.451 | 0.463 | 0.448 | 0.434 | 0.414 | 0.418 | 0.411 | 0.337 | 0.170 | 0.555 | 0.425 |
| Ws-3 | 0.412 | 0.464 | 0.466 | 0.441 | 0.413 | 0.394 | 0.406 | 0.380 | 0.294 | 0.204 | 0.617 | 0.458 |
| Wt-3 | 0.457 | 0.485 | 0.465 | 0.418 | 0.408 | 0.422 | 0.391 | 0.317 | 0.254 | 0.181 | 0.573 | 0.440 |
| Wt-5 | 0.400 | 0.433 | 0.442 | 0.421 | 0.418 | 0.413 | 0.420 | 0.400 | 0.316 | 0.204 | 0.582 | 0.485 |
| Yo-0 | 0.417 | 0.444 | 0.433 | 0.410 | 0.395 | 0.399 | 0.393 | 0.340 | 0.273 | 0.202 | 0.582 | 0.486 |
| Zdr-1 | 0.432 | 0.454 | 0.442 | 0.423 | 0.420 | 0.420 | 0.420 | 0.391 | 0.340 | 0.224 | 0.616 | 0.501 |
| Zdr-6 | 0.428 | 0.469 | 0.452 | 0.414 | 0.392 | 0.381 | 0.373 | 0.343 | 0.310 | 0.188 | 0.602 | 0.434 |
| Zdrl2-24 | 0.427 | 0.446 | 0.440 | 0.413 | 0.390 | 0.385 | 0.392 | 0.350 | 0.265 | 0.211 | 0.597 | 0.484 |
| Zdrl2-25 | 0.452 | 0.473 | 0.451 | 0.418 | 0.413 | 0.416 | 0.417 | 0.373 | 0.301 | 0.175 | 0.569 | 0.425 |
| Zü-1 | 0.399 | 0.433 | 0.444 | 0.419 | 0.387 | 0.380 | 0.376 | 0.333 | 0.273 | 0.244 | 0.632 | 0.522 |
