## Supplementary Table S3 for "Temporal dynamics of QTL effects on vegetative growth in *Arabidopsis thaliana*"

| MTA | trait | SNP | Chrom | position | p.value | q.value | effect | PVE% | PC | category |
|---|---|---|---|---|---|---|---|---|---|---|
| | PLA11 | M1_00210424 | 1 | 210,424 | 2.41E-07 | 6.38E-03 | 1.416 | 1.567 | 3 | |
| | RGR09_11 | M1_00410242 | 1 | 410,242 | 4.82E-09 | 3.53E-04 | 0.009 | 0.412 | 4 | |
| | RGR14_16 | M1_01037424 | 1 | 1,037,424 | 1.65E-06 | 3.19E-02 | -0.004 | 0.894 | 3 | |
| | RGR10_12 | M1_01877206 | 1 | 1,877,206 | 2.87E-07 | 1.52E-02 | 0.009 | 2.635 | 4 | |
| | **PLA10** | **M1_02376375** | **1** | **2,376,375** | **1.90E-06** | **4.03E-02** | **-0.542** | **0.762** | **2** | dynamic |
| | **PLA11** | **M1_02376375** | **1** | **2,376,375** | **5.31E-14** | **1.13E-08** | **-1.379** | **2.925** | **3** | co-loc |
| 1-01 | **PLA12** | **M1_02376375** | **1** | **2,376,375** | **1.66E-12** | **3.52E-07** | **-1.993** | **2.766** | **3** | |
| | **PLA14** | **M1_02376375** | **1** | **2,376,375** | **1.95E-06** | **4.59E-02** | **-2.966** | **1.182** | **3** | |
| | **RGR15_17** | **M1_02376375** | **1** | **2,376,375** | **2.84E-08** | **8.60E-04** | **0.005** | **2.061** | **4** | |
| 1-06 | **SA** | **M1_05508567** | **1** | **5,508,567** | **1.45E-06** | **3.08E-02** | **0.009** | **1.741** | **5** | co-loc |
| | **RGR10_12** | **M1_05510455** | **1** | **5,510,455** | **2.84E-07** | **1.52E-02** | **0.005** | **1.284** | **4** | |
| | RGR12_14 | M1_08705575 | 1 | 8,705,575 | 6.00E-08 | 2.54E-03 | 0.004 | 3.339 | 3 | |
| | RGR09_11 | M1_08974266 | 1 | 8,974,266 | 3.99E-07 | 1.69E-02 | 0.004 | 2.256 | 4 | |
| | SW | M1_09513015 | 1 | 9,513,015 | 6.43E-07 | 1.95E-02 | -0.008 | 1.575 | 2 | |
| 1-04 | **SA** | **M1_10358650** | **1** | **10,358,650** | **7.90E-07** | **1.86E-02** | **-0.007** | **1.109** | **5** | co-loc |
| | **SW** | **M1_10359910** | **1** | **10,359,910** | **5.51E-08** | **2.92E-03** | **-0.008** | **1.924** | **2** | |
| | PLA18 | M1_11687852 | 1 | 11,687,852 | 3.84E-07 | 7.50E-03 | 9.427 | 2.314 | 3 | |
| | RGR12_14 | M1_12191525 | 1 | 12,191,525 | 1.80E-07 | 4.25E-03 | 0.005 | 0.344 | 3 | |
| | PLA18 | M1_13874540 | 1 | 13,874,540 | 7.52E-07 | 1.33E-02 | -10.747 | 2.560 | 3 | |
| | RGR09_11 | M1_18192320 | 1 | 18,192,320 | 9.61E-09 | 5.10E-04 | -0.008 | 4.332 | 4 | |
| | RGR09_11 | M1_19567276 | 1 | 19,567,276 | 1.52E-06 | 2.94E-02 | 0.005 | 0.744 | 4 | |
| | SW | M1_19639293 | 1 | 19,639,293 | 7.51E-09 | 7.97E-04 | 0.012 | 2.342 | 2 | |
| | RGR09_11 | M1_21234054 | 1 | 21,234,054 | 5.00E-09 | 3.53E-04 | -0.007 | 3.662 | 4 | |
| | RGR15_17 | M1_21332233 | 1 | 21,332,233 | 2.21E-06 | 3.90E-02 | -0.004 | 0.161 | 4 | |
| | DW20 | M1_21696415 | 1 | 21,696,415 | 4.11E-08 | 1.25E-03 | -0.748 | 0.912 | 3 | |
| | **PLA13** | **M1_22361861** | **1** | **22,361,861** | **9.79E-09** | **5.19E-04** | **2.975** | **3.324** | **3** | dynamic |
| | **PLA14** | **M1_22361861** | **1** | **22,361,861** | **1.73E-09** | **1.22E-04** | **4.676** | **2.627** | **3** | |
| | **PLA15** | **M1_22361861** | **1** | **22,361,861** | **4.68E-08** | **1.65E-03** | **6.235** | **2.897** | **3** | |
| | **PLA16** | **M1_22362243** | **1** | **22,362,243** | **9.69E-09** | **1.03E-03** | **7.915** | **2.220** | **4** | |
| 1-02 | **PLA17** | **M1_22362243** | **1** | **22,362,243** | **2.48E-07** | **1.32E-02** | **8.040** | **1.934** | **3** | |

| MTA | trait | SNP | Chrom | position | p.value | q.value | effect | PVE% | PC | category |
|---|---|---|---|---|---|---|---|---|---|---|
| | **PLA18** | **M1_22362243** | **1** | **22,362,243** | **4.92E-10** | **2.09E-05** | **12.163** | **3.194** | **3** | |
| | **PLA10** | **M1_22363423** | **1** | **22,363,423** | **1.14E-11** | **1.21E-06** | **-0.837** | **2.796** | **2** | |
| | **PLA11** | **M1_22363423** | **1** | **22,363,423** | **1.12E-11** | **1.19E-06** | **-1.310** | **1.948** | **3** | |
| | **PLA12** | **M1_22363423** | **1** | **22,363,423** | **1.16E-11** | **1.23E-06** | **-1.977** | **2.063** | **3** | |
| 1-03 | **RGR09_11** | **M1_22381328** | **1** | **22,381,328** | **1.53E-06** | **2.94E-02** | **0.005** | **0.426** | **4** | dynamic |
| | **RGR08_10** | **M1_22382076** | **1** | **22,382,076** | **2.36E-12** | **5.00E-07** | **-0.007** | **2.019** | **3** | co-loc |
| | SL | M1_23244664 | 1 | 23,244,664 | 8.39E-07 | 1.62E-02 | 0.010 | 2.449 | 3 | |
| | RGR14_16 | M1_24088811 | 1 | 24,088,811 | 2.11E-09 | 7.46E-05 | -0.005 | 3.649 | 3 | |
| | PLA08 | M1_24292814 | 1 | 24,292,814 | 3.49E-08 | 1.85E-03 | -0.445 | 1.541 | 3 | |
| | PLA17 | M1_24698430 | 1 | 24,698,430 | 8.15E-07 | 2.88E-02 | 7.907 | 0.185 | 3 | |
| | SL | M1_25191846 | 1 | 25,191,846 | 1.16E-12 | 2.45E-07 | 0.011 | 3.453 | 3 | |
| | SA | M1_25694302 | 1 | 25,694,302 | 1.08E-07 | 4.73E-03 | 0.010 | 1.877 | 5 | |
| | DW20 | M1_26935804 | 1 | 26,935,804 | 2.77E-08 | 9.80E-04 | -0.462 | 1.860 | 3 | |
| | RGR12_14 | M1_27458647 | 1 | 27,458,647 | 1.74E-07 | 4.25E-03 | 0.012 | 3.098 | 3 | |
| | RGR12_14 | M1_27742038 | 1 | 27,742,038 | 3.79E-10 | 4.02E-05 | -0.008 | 1.148 | 3 | |
| | PLA07 | M1_27942612 | 1 | 27,942,612 | 7.00E-07 | 1.65E-02 | 0.204 | 1.879 | 3 | |
| | RGR11_13 | M1_29064168 | 1 | 29,064,168 | 5.91E-10 | 4.18E-05 | -0.014 | 3.710 | | |
| | PLA09 | M1_29363132 | 1 | 29,363,132 | 6.33E-07 | 2.69E-02 | -0.500 | 2.011 | 3 | |
| | RGR14_16 | M1_30223019 | 1 | 30,223,019 | 8.17E-08 | 2.17E-03 | 0.005 | 1.947 | 3 | |
| | SL | M1_30276962 | 1 | 30,276,962 | 2.05E-07 | 6.20E-03 | -0.008 | 1.062 | 3 | |
| | RGR16_18 | M1_30345995 | 1 | 30,345,995 | 1.01E-06 | 2.67E-02 | 0.011 | 0.904 | 3 | |
| | RGR07_09 | M2_00167648 | 2 | 167,648 | 6.19E-08 | 4.37E-03 | 0.004 | 1.596 | 3 | |
| | SL | M2_00772623 | 2 | 772,623 | 2.64E-07 | 6.99E-03 | -0.021 | 2.339 | 3 | |
| 2-01 | **PLA13** | **M2_01550771** | **2** | **1,550,771** | **1.72E-06** | **4.05E-02** | **-2.353** | **2.469** | **3** | dynamic |
| | **PLA14** | **M2_01550771** | **2** | **1,550,771** | **4.47E-07** | **1.35E-02** | **-3.906** | **2.593** | **3** | |
| 2-02 | **PLA07** | **M2_02536548** | **2** | **2,536,548** | **1.18E-09** | **8.34E-05** | **-0.323** | **1.600** | **3** | co-loc |
| | **PLA09** | **M2_02536548** | **2** | **2,536,548** | **1.97E-08** | **3.25E-03** | **-0.662** | **1.141** | **3** | |
| | **PLA11** | **M2_02536548** | **2** | **2,536,548** | **1.34E-07** | **4.06E-03** | **-1.585** | **0.681** | **3** | |
| | RGR15_17 | M2_02703452 | 2 | 2,703,452 | 5.31E-17 | 1.13E-11 | -0.009 | 3.039 | 4 | |
| 2-05 | **SA** | **M2_04367512** | **2** | **4,367,512** | **5.14E-10** | **3.64E-05** | **-0.008** | **1.969** | **5** | co-loc |

| MTA | trait | SNP | Chrom | position | p.value | q.value | effect | PVE% | PC | category |
|---|---|---|---|---|---|---|---|---|---|---|
| 2-0[illegible] | **SW** | **M2_04367512** | **2** | **4,367,512** | **2.25E-09** | **4.78E-04** | **-0.010** | **2.668** | **2** | |
| | PLA11 | M2_04418829 | 2 | 4,418,829 | 9.39E-08 | 3.32E-03 | -1.005 | 1.534 | 3 | |
| | PLA09 | M2_04871627 | 2 | 4,871,627 | 1.00E-06 | 3.54E-02 | 0.385 | 2.916 | 3 | |
| | PLA07 | M2_05370870 | 2 | 5,370,870 | 9.00E-08 | 2.73E-03 | 0.189 | 1.280 | 3 | |
| | RGR12_14 | M2_05929477 | 2 | 5,929,477 | 7.93E-08 | 2.80E-03 | -0.003 | 1.083 | 3 | |
| | RGR16_18 | M2_07773633 | 2 | 7,773,633 | 2.94E-08 | 1.04E-03 | 0.010 | 2.051 | 3 | |
| | SL | M2_07928447 | 2 | 7,928,447 | 2.26E-06 | 4.00E-02 | 0.009 | 0.082 | 3 | |
| | PLA11 | M2_11140052 | 2 | 11,140,052 | 3.45E-07 | 8.12E-03 | 0.906 | 1.091 | 3 | |
| 2-03 | **PLA17** | **M2_11534875** | **2** | **11,534,875** | **5.02E-08** | **3.55E-03** | **-22.681** | **2.238** | **3** | co-loc |
| | **RGR09_11** | **M2_11534875** | **2** | **11,534,875** | **1.46E-06** | **2.94E-02** | **-0.009** | **1.955** | **4** | |
| | PLA11 | M2_11553940 | 2 | 11,553,940 | 5.26E-11 | 3.72E-06 | 1.934 | 2.098 | 3 | |
| | RGR14_16 | M2_11647693 | 2 | 11,647,693 | 9.16E-10 | 3.89E-05 | -0.004 | 2.093 | 3 | |
| | RGR07_09 | M2_11691472 | 2 | 11,691,472 | 2.33E-06 | 4.49E-02 | 0.004 | 1.256 | 3 | |
| | PLA13 | M2_11760256 | 2 | 11,760,256 | 7.60E-07 | 2.30E-02 | 2.300 | 1.401 | 3 | |
| | PLA10 | M2_12008468 | 2 | 12,008,468 | 2.07E-07 | 6.28E-03 | 0.624 | 2.222 | 2 | |
| 2-04 | **PLA16** | **M2_13077250** | **2** | **13,077,250** | **2.85E-07** | **7.55E-03** | **9.452** | **1.541** | **4** | dynamic |
| | **PLA17** | **M2_13077250** | **2** | **13,077,250** | **5.68E-09** | **1.20E-03** | **11.866** | **2.682** | **3** | |
| | DW20 | M2_13167359 | 2 | 13,167,359 | 8.74E-11 | 6.18E-06 | -0.546 | 2.665 | 3 | |
| | SW | M2_13226456 | 2 | 13,226,456 | 8.02E-08 | 3.40E-03 | 0.011 | 3.325 | 2 | |
| | SA | M2_15289966 | 2 | 15,289,966 | 2.17E-06 | 4.19E-02 | -0.006 | 2.712 | 5 | |
| | PLA08 | M2_16072194 | 2 | 16,072,194 | 4.60E-07 | 1.95E-02 | -0.262 | 0.579 | 3 | |
| | SL | M2_17711412 | 2 | 17,711,412 | 1.90E-07 | 6.20E-03 | 0.009 | 3.119 | 3 | |
| | PLA18 | M2_18850087 | 2 | 18,850,087 | 2.94E-12 | 6.25E-07 | 29.534 | 2.823 | 3 | |
| | RGR12_14 | M2_19036527 | 2 | 19,036,527 | 3.86E-09 | 2.73E-04 | 0.010 | 5.546 | 3 | |
| | DW20 | M2_19627477 | 2 | 19,627,477 | 1.21E-06 | 2.56E-02 | -0.328 | 2.407 | 3 | |
| 3-01 | **RGR15_17** | **M3_02221399** | **3** | **2,221,399** | **5.36E-09** | **2.27E-04** | **0.008** | **3.174** | **4** | dynamic |
| | **RGR16_18** | **M3_02221399** | **3** | **2,221,399** | **9.02E-09** | **4.79E-04** | **0.011** | **1.244** | **3** | |
| | RGR12_14 | M3_03527195 | 3 | 3,527,195 | 1.44E-07 | 4.25E-03 | -0.005 | 1.381 | 3 | |
| | PLA10 | M3_04265482 | 3 | 4,265,482 | 5.96E-07 | 1.40E-02 | 0.760 | 2.080 | 2 | |
| | PLA18 | M3_04294414 | 3 | 4,294,414 | 3.89E-07 | 7.50E-03 | 13.165 | 2.964 | 3 | |

| MTA | trait | SNP | Chrom | position | p.value | q.value | effect | PVE% | PC | category |
|---|---|---|---|---|---|---|---|---|---|---|
| 3-02 | **PLA13** | **M3_04302571** | **3** | **4,302,571** | **1.73E-07** | **6.11E-03** | **3.131** | **3.302** | **3** | co-loc |
| | **PLA16** | **M3_04302571** | **3** | **4,302,571** | **1.71E-08** | **1.21E-03** | **10.726** | **4.600** | **4** | |
| | RGR09_11 | M3_04402193 | 3 | 4,402,193 | 1.21E-06 | 2.94E-02 | -0.004 | 3.561 | 4 | |
| | RGR14_16 | M3_04412292 | 3 | 4,412,292 | 2.95E-14 | 3.13E-09 | -0.012 | 5.950 | 3 | |
| 3-03 | **RGR11_13** | **M3_04759689** | **3** | **4,759,689** | **8.75E-08** | **4.64E-03** | **0.004** | **3.321** | | co-loc |
| | **PLA18** | **M3_04763256** | **3** | **4,763,256** | **3.03E-06** | **4.29E-02** | **-9.639** | **1.942** | **3** | |
| | DW20 | M3_06362499 | 3 | 6,362,499 | 6.25E-09 | 3.31E-04 | -0.357 | 2.481 | 3 | |
| | PLA15 | M3_06754875 | 3 | 6,754,875 | 1.93E-07 | 5.12E-03 | -6.671 | 3.012 | 3 | |
| | PLA16 | M3_07522893 | 3 | 7,522,893 | 2.98E-08 | 1.58E-03 | -12.917 | 1.359 | 4 | |
| | RGR12_14 | M3_08038617 | 3 | 8,038,617 | 1.93E-06 | 3.41E-02 | -0.004 | 0.723 | 3 | |
| 3-04 | **PLA07** | **M3_08552586** | **3** | **8,552,586** | **3.71E-07** | **9.84E-03** | **-0.243** | **0.785** | **3** | dynamic |
| | **PLA09** | **M3_08553143** | **3** | **8,553,143** | **3.23E-08** | **3.25E-03** | **-0.533** | **4.199** | **3** | |
| | **PLA10** | **M3_08553143** | **3** | **8,553,143** | **2.01E-07** | **6.28E-03** | **-0.776** | **3.701** | **2** | |
| | RGR15_17 | M3_08553143 | 3 | 8,553,143 | 2.42E-08 | 8.54E-04 | 0.006 | 3.536 | 4 | |
| | RGR15_17 | M3_08746180 | 3 | 8,746,180 | 2.85E-09 | 1.51E-04 | 0.006 | 1.270 | 4 | |
| | RGR12_14 | M3_08886483 | 3 | 8,886,483 | 5.37E-07 | 1.04E-02 | -0.003 | 1.109 | 3 | |
| | RGR11_13 | M3_09144102 | 3 | 9,144,102 | 4.88E-10 | 4.18E-05 | 0.004 | 1.331 | | |
| | SA | M3_09285086 | 3 | 9,285,086 | 1.54E-07 | 5.46E-03 | -0.007 | 2.932 | 5 | |
| | RGR07_09 | M3_10489878 | 3 | 10,489,878 | 1.26E-11 | 1.34E-06 | 0.006 | 4.161 | 3 | |
| | RGR07_09 | M3_10502507 | 3 | 10,502,507 | 3.13E-07 | 1.10E-02 | 0.004 | 0.398 | 3 | |
| | SA | M3_11387053 | 3 | 11,387,053 | 1.11E-07 | 4.73E-03 | 0.008 | 2.552 | 5 | |
| 3-09 | **SL** | **M3_11700017** | **3** | **11,700,017** | **4.26E-10** | **4.52E-05** | **-0.009** | **3.431** | **3** | co-loc |
| | **SW** | **M3_11700017** | **3** | **11,700,017** | **1.44E-08** | **1.02E-03** | **-0.008** | **2.396** | **2** | |
| | PLA17 | M3_11913423 | 3 | 11,913,423 | 1.59E-06 | 4.81E-02 | 7.644 | 1.739 | 3 | |
| | RGR11_13 | M3_12191390 | 3 | 12,191,390 | 1.98E-07 | 8.41E-03 | 0.003 | 1.824 | | |
| | PLA12 | M3_12286474 | 3 | 12,286,474 | 7.92E-07 | 2.10E-02 | 1.500 | 0.487 | 3 | |
| 3-06 | **PLA07** | **M3_12590114** | **3** | **12,590,114** | **6.95E-09** | **2.95E-04** | **-0.581** | **4.066** | **3** | dynamic |
| | **PLA12** | **M3_12590114** | **3** | **12,590,114** | **1.16E-06** | **2.73E-02** | **-4.059** | **4.134** | **3** | |
| | **PLA13** | **M3_12590114** | **3** | **12,590,114** | **1.44E-09** | **1.53E-04** | **-8.315** | **4.348** | **3** | |
| | **PLA15** | **M3_12590114** | **3** | **12,590,114** | **7.58E-07** | **1.79E-02** | **-14.456** | **2.164** | **3** | |

| MTA | trait | SNP | Chrom | position | p.value | q.value | effect | PVE% | PC | category |
|---|---|---|---|---|---|---|---|---|---|---|
| | **PLA16** | **M3_12590114** | **3** | **12,590,114** | **1.41E-07** | **4.47E-03** | **-23.093** | **2.591** | **4** | |
| | RGR07_09 | M3_12820769 | 3 | 12,820,769 | 1.52E-06 | 3.23E-02 | -0.004 | 0.670 | 3 | |
| | RGR14_16 | M3_13969810 | 3 | 13,969,810 | 9.24E-08 | 2.18E-03 | -0.005 | 2.290 | 3 | |
| | RGR07_09 | M3_13978043 | 3 | 13,978,043 | 2.07E-07 | 8.77E-03 | 0.006 | 1.771 | 3 | |
| 3-08 | **PLA07** | **M3_14672819** | **3** | **14,672,819** | **1.14E-06** | **2.42E-02** | **0.239** | **2.625** | **3** | dynamic |
| | **PLA08** | **M3_14672819** | **3** | **14,672,819** | **4.52E-11** | **9.42E-06** | **0.527** | **1.685** | **3** | |
| | RGR16_18 | M3_14902609 | 3 | 14,902,609 | 4.84E-10 | 5.14E-05 | 0.007 | 4.110 | 3 | |
| | RGR13_15 | M3_15966173 | 3 | 15,966,173 | 5.64E-08 | 3.99E-03 | 0.005 | 3.282 | | |
| | SL | M3_16235890 | 3 | 16,235,890 | 1.40E-08 | 9.90E-04 | -0.010 | 1.427 | 3 | |
| | RGR11_13 | M3_16945362 | 3 | 16,945,362 | 1.05E-06 | 3.71E-02 | 0.009 | 3.031 | | |
| | RGR15_17 | M3_17117911 | 3 | 17,117,911 | 6.68E-08 | 1.57E-03 | 0.006 | 1.954 | 4 | |
| | RGR08_10 | M3_17868286 | 3 | 17,868,286 | 8.80E-07 | 3.73E-02 | 0.005 | 0.262 | 3 | |
| | RGR16_18 | M3_18211638 | 3 | 18,211,638 | 2.26E-10 | 4.80E-05 | -0.019 | 3.089 | 3 | |
| 3-07 | **PLA12** | **M3_18315086** | **3** | **18,315,086** | **2.49E-10** | **1.76E-05** | **2.758** | **3.704** | **3** | dynamic |
| | **PLA13** | **M3_18315086** | **3** | **18,315,086** | **8.27E-08** | **3.51E-03** | **3.692** | **4.176** | **3** | |
| | **PLA14** | **M3_18315086** | **3** | **18,315,086** | **1.33E-11** | **1.41E-06** | **7.292** | **5.260** | **3** | |
| | **PLA15** | **M3_18315086** | **3** | **18,315,086** | **6.33E-12** | **6.72E-07** | **11.013** | **5.686** | **3** | |
| | **PLA16** | **M3_18315086** | **3** | **18,315,086** | **4.07E-10** | **8.64E-05** | **14.361** | **4.808** | **4** | |
| | **PLA17** | **M3_18315086** | **3** | **18,315,086** | **5.27E-07** | **2.24E-02** | **12.896** | **4.108** | **3** | |
| | **DW20** | **M3_18315086** | **3** | **18,315,086** | **7.39E-07** | **1.74E-02** | **0.455** | **3.342** | **3** | |
| | **PLA18** | **M3_18315086** | **3** | **18,315,086** | **1.86E-10** | **9.86E-06** | **19.441** | **4.939** | **3** | |
| | PLA10 | M3_18371403 | 3 | 18,371,403 | 2.61E-07 | 6.91E-03 | 0.698 | 3.029 | 2 | |
| | RGR08_10 | M3_21777341 | 3 | 21,777,341 | 7.77E-08 | 4.12E-03 | 0.004 | 0.461 | 3 | |
| | SL | M3_23160992 | 3 | 23,160,992 | 5.88E-08 | 2.50E-03 | 0.022 | 4.826 | 3 | |
| | PLA18 | M4_00143220 | 4 | 143,220 | 3.01E-06 | 4.29E-02 | 8.650 | 1.499 | 3 | |
| | RGR15_17 | M4_00325002 | 4 | 325,002 | 5.50E-08 | 1.46E-03 | 0.008 | 1.378 | 4 | |
| | RGR08_10 | M4_02767296 | 4 | 2,767,296 | 3.33E-11 | 3.53E-06 | -0.008 | 2.705 | 3 | |
| 4-08 | **SL** | **M4_03192692** | **4** | **3,192,692** | **1.42E-07** | **1.51E-02** | **0.008** | **1.451** | **3** | co-loc |
| | **SW** | **M4_03192692** | **4** | **3,192,692** | **1.33E-06** | **3.52E-02** | **0.007** | **1.030** | **2** | |
| | RGR14_16 | M4_04864129 | 4 | 4,864,129 | 3.68E-08 | 1.11E-03 | 0.007 | 2.267 | 3 | |

| MTA | trait | SNP | Chrom | position | p.value | q.value | effect | PVE% | PC | category |
|---|---|---|---|---|---|---|---|---|---|---|
| 4-01 | **RGR13_15** | **M4_05389299** | **4** | **5,389,299** | **6.85E-07** | **2.91E-02** | **-0.006** | **3.106** | | dynamic |
| | **RGR14_16** | **M4_05389975** | **4** | **5,389,975** | **4.39E-11** | **3.10E-06** | **-0.009** | **4.336** | **3** | |
| | RGR07_09 | M4_05433949 | 4 | 5,433,949 | 5.03E-07 | 1.52E-02 | 0.004 | 1.412 | 3 | |
| | PLA18 | M4_06092503 | 4 | 6,092,503 | 1.81E-06 | 2.96E-02 | -11.355 | 1.969 | 3 | |
| 4-02 | **PLA12** | **M4_06682595** | **4** | **6,682,595** | **1.37E-06** | **2.91E-02** | **2.167** | **0.670** | **3** | dynamic |
| | **RGR15_17** | **M4_06682595** | **4** | **6,682,595** | **3.54E-13** | **3.75E-08** | **-0.011** | **2.993** | **4** | co-loc |
| | **RGR16_18** | **M4_06682595** | **4** | **6,682,595** | **1.85E-08** | **7.84E-04** | **-0.011** | **1.014** | **3** | |
| | RGR07_09 | M4_07679303 | 4 | 7,679,303 | 1.52E-06 | 3.23E-02 | 0.009 | 3.391 | 3 | |
| 4-03 | **PLA14** | **M4_07928602** | **4** | **7,928,602** | **4.50E-08** | **1.59E-03** | **-3.606** | **2.815** | **3** | dynamic |
| | **PLA15** | **M4_07928602** | **4** | **7,928,602** | **1.61E-07** | **4.87E-03** | **-5.200** | **3.399** | **3** | |
| 4-04 | **PLA16** | **M4_08007078** | **4** | **8,007,078** | **2.15E-06** | **4.15E-02** | **6.867** | **1.769** | **4** | dynamic |
| | **PLA17** | **M4_08007078** | **4** | **8,007,078** | **2.92E-08** | **3.10E-03** | **9.214** | **0.546** | **3** | |
| | PLA12 | M4_08021147 | 4 | 8,021,147 | 3.59E-07 | 1.27E-02 | 2.681 | 1.316 | 3 | |
| | DW20 | M4_08007078 | 4 | 8,007,078 | 1.23E-08 | 5.23E-04 | 0.362 | 1.071 | 3 | |
| | RGR07_09 | M4_08604864 | 4 | 8,604,864 | 2.43E-12 | 5.16E-07 | -0.009 | 1.701 | 3 | |
| | SL | M4_09030795 | 4 | 9,030,795 | 4.63E-08 | 2.46E-03 | 0.016 | 1.573 | 3 | |
| 4-05 | **PLA08** | **M4_10565446** | **4** | **10,565,446** | **7.92E-10** | **5.60E-05** | **0.446** | **3.067** | **3** | dynamic |
| | **PLA09** | **M4_10565446** | **4** | **10,565,446** | **1.86E-07** | **9.89E-03** | **0.561** | **2.614** | **3** | |
| | **PLA10** | **M4_10565446** | **4** | **10,565,446** | **2.55E-08** | **1.35E-03** | **0.910** | **3.304** | **2** | |
| | **PLA11** | **M4_10565446** | **4** | **10,565,446** | **8.14E-08** | **3.32E-03** | **1.370** | **3.309** | **3** | |
| | **PLA14** | **M4_10565446** | **4** | **10,565,446** | **6.28E-07** | **1.66E-02** | **4.727** | **2.656** | **3** | |
| | PLA07 | M4_10663543 | 4 | 10,663,543 | 4.01E-15 | 8.51E-10 | 0.718 | 2.724 | 3 | |
| | PLA18 | M4_10758117 | 4 | 10,758,117 | 5.80E-11 | 4.10E-06 | 30.628 | 3.220 | 3 | |
| | RGR09_11 | M4_10801467 | 4 | 10,801,467 | 8.94E-07 | 2.94E-02 | -0.007 | 2.399 | 4 | |
| | RGR08_10 | M4_11477061 | 4 | 11,477,061 | 1.07E-08 | 7.59E-04 | 0.009 | 2.548 | 3 | |
| 4-06 | **PLA10** | **M4_16453234** | **4** | **16,453,234** | **2.45E-10** | **1.73E-05** | **1.716** | **2.599** | **2** | dynamic |
| | **PLA11** | **M4_16453234** | **4** | **16,453,234** | **5.15E-09** | **2.73E-04** | **2.639** | **2.261** | **3** | |
| | **PLA12** | **M4_16453234** | **4** | **16,453,234** | **5.51E-10** | **2.92E-05** | **4.387** | **3.232** | **3** | |
| | **PLA13** | **M4_16453234** | **4** | **16,453,234** | **5.62E-13** | **1.19E-07** | **8.256** | **2.933** | **3** | |
| | **PLA14** | **M4_16453234** | **4** | **16,453,234** | **1.43E-15** | **3.03E-10** | **13.992** | **2.953** | **3** | |

| MTA | trait | SNP | Chrom | position | p.value | q.value | effect | PVE% | PC | category |
|---|---|---|---|---|---|---|---|---|---|---|
| | **PLA15** | **M4_16453234** | **4** | **16,453,234** | **8.56E-14** | **1.82E-08** | **19.070** | **2.946** | **3** | |
| | **PLA16** | **M4_16453234** | **4** | **16,453,234** | **1.48E-07** | **4.47E-03** | **18.245** | **3.866** | **4** | |
| | DW20 | M4_16524186 | 4 | 16,524,186 | 1.80E-11 | 1.91E-06 | -1.085 | 2.248 | 3 | |
| | PLA18 | M4_16538982 | 4 | 16,538,982 | 2.70E-07 | 7.16E-03 | -25.426 | 1.978 | 3 | |
| 4-09 | **SA** | **M4_17448257** | **4** | **17,448,257** | **2.11E-07** | **6.38E-03** | **0.006** | **3.144** | **5** | co-loc |
| | **SW** | **M4_17448257** | **4** | **17,448,257** | **1.04E-07** | **3.69E-03** | **0.008** | **2.180** | **2** | |
| 4-07 | **RGR09_11** | **M4_17704759** | **4** | **17,704,759** | **6.22E-14** | **1.32E-08** | **0.011** | **4.340** | **4** | dynamic |
| | **RGR10_12** | **M4_17704759** | **4** | **17,704,759** | **2.43E-08** | **2.58E-03** | **0.007** | **3.873** | **4** | |
| | **RGR11_13** | **M4_17704759** | **4** | **17,704,759** | **4.03E-19** | **8.55E-14** | **0.010** | **6.572** | | |
| | **RGR12_14** | **M4_17704759** | **4** | **17,704,759** | **3.44E-12** | **7.31E-07** | **0.007** | **8.103** | **3** | |
| | **RGR13_15** | **M4_17704759** | **4** | **17,704,759** | **1.79E-09** | **1.90E-04** | **0.007** | **6.204** | | |
| | SA | M4_18448754 | 4 | 18,448,754 | 5.70E-07 | 1.51E-02 | 0.009 | 1.407 | 5 | |
| | SL | M5_00408775 | 5 | 408,775 | 3.38E-07 | 7.98E-03 | -0.019 | 1.538 | 3 | |
| 5-01 | **DW20** | **M5_01196098** | **5** | **1,196,098** | **1.70E-12** | **3.60E-07** | **-0.855** | **6.140** | **3** | dynamic |
| | **PLA13** | **M5_01196098** | **5** | **1,196,098** | **1.18E-06** | **3.14E-02** | **-4.017** | **1.797** | **3** | |
| | **PLA14** | **M5_01196098** | **5** | **1,196,098** | **2.87E-08** | **1.22E-03** | **-7.185** | **2.614** | **3** | |
| | **PLA15** | **M5_01196098** | **5** | **1,196,098** | **7.31E-11** | **5.17E-06** | **-12.655** | **3.357** | **3** | |
| | **PLA16** | **M5_01196098** | **5** | **1,196,098** | **1.14E-07** | **4.47E-03** | **-13.912** | **3.837** | **4** | |
| | **PLA18** | **M5_01196098** | **5** | **1,196,098** | **6.68E-09** | **2.36E-04** | **-20.718** | **3.538** | **3** | |
| | SL | M5_01735888 | 5 | 1,735,888 | 5.39E-07 | 1.14E-02 | 0.016 | 1.245 | 3 | |
| | PLA15 | M5_02262258 | 5 | 2,262,258 | 1.39E-06 | 2.95E-02 | 4.917 | 1.618 | 3 | |
| | RGR16_18 | M5_03632947 | 5 | 3,632,947 | 3.59E-09 | 2.54E-04 | 0.007 | 4.501 | 3 | |
| | SA | M5_03824273 | 5 | 3,824,273 | 1.18E-10 | 1.25E-05 | 0.007 | 2.109 | 5 | |
| | RGR12_14 | M5_04218274 | 5 | 4,218,274 | 3.74E-07 | 7.94E-03 | 0.006 | 1.053 | 3 | |
| | RGR15_17 | M5_05721619 | 5 | 5,721,619 | 6.48E-10 | 4.58E-05 | 0.012 | 0.930 | 4 | |
| | PLA16 | M5_07198259 | 5 | 7,198,259 | 1.18E-06 | 2.50E-02 | 6.774 | 1.465 | 4 | |
| 5-02 | **RGR13_15** | **M5_08237784** | **5** | **8,237,784** | **1.38E-07** | **7.30E-03** | **-0.009** | **4.367** | | dynamic |
| | **RGR14_16** | **M5_08237784** | **5** | **8,237,784** | **4.02E-17** | **8.52E-12** | **-0.016** | **4.618** | **3** | |
| | PLA07 | M5_08325464 | 5 | 8,325,464 | 7.82E-10 | 8.30E-05 | 0.512 | 2.070 | 3 | |
| | SA | M5_09944034 | 5 | 9,944,034 | 3.66E-12 | 7.76E-07 | 0.008 | 2.981 | 5 | |

| MTA | trait | SNP | Chrom | position | p.value | q.value | effect | PVE% | PC | category |
|---|---|---|---|---|---|---|---|---|---|---|
| | RGR14_16 | M5_11037195 | 5 | 11,037,195 | 1.38E-07 | 2.92E-03 | -0.005 | 2.609 | 3 | |
| | PLA10 | M5_12452393 | 5 | 12,452,393 | 4.78E-08 | 2.03E-03 | 0.574 | 0.305 | 2 | |
| | RGR07_09 | M5_12510529 | 5 | 12,510,529 | 2.00E-07 | 8.77E-03 | -0.013 | 6.900 | 3 | |
| | RGR16_18 | M5_13188628 | 5 | 13,188,628 | 3.14E-07 | 9.51E-03 | -0.006 | 0.379 | 3 | |
| | RGR13_15 | M5_13643155 | 5 | 13,643,155 | 2.47E-10 | 5.24E-05 | -0.005 | 2.424 | | |
| 5-03 | **PLA13** | **M5_13811024** | **5** | **13,811,024** | **4.30E-09** | **3.04E-04** | **3.322** | **2.876** | **3** | dynamic |
| | **PLA14** | **M5_13811024** | **5** | **13,811,024** | **1.22E-08** | **6.45E-04** | **4.742** | **3.987** | **3** | |
| | **PLA15** | **M5_13811024** | **5** | **13,811,024** | **3.28E-09** | **1.39E-04** | **7.447** | **3.244** | **3** | |
| | RGR15_17 | M5_14117057 | 5 | 14,117,057 | 3.06E-07 | 6.50E-03 | 0.006 | 0.281 | 4 | |
| | RGR07_09 | M5_14236451 | 5 | 14,236,451 | 6.20E-07 | 1.64E-02 | -0.004 | 1.033 | 3 | |
| | PLA12 | M5_16425089 | 5 | 16,425,089 | 6.42E-07 | 1.95E-02 | 1.310 | 1.397 | 3 | |
| | DW20 | M5_17256023 | 5 | 17,256,023 | 5.04E-08 | 1.34E-03 | -0.380 | 3.066 | 3 | |
| | RGR10_12 | M5_18091009 | 5 | 18,091,009 | 7.18E-09 | 1.52E-03 | -0.008 | 1.420 | 4 | |
| 5-07 | **PLA07** | **M5_18554094** | **5** | **18,554,094** | **3.18E-09** | **1.69E-04** | **0.201** | **3.624** | **3** | dynamic |
| | **PLA08** | **M5_18554094** | **5** | **18,554,094** | **8.88E-11** | **9.42E-06** | **0.335** | **2.530** | **3** | |
| | RGR14_16 | M5_18982082 | 5 | 18,982,082 | 5.20E-10 | 2.76E-05 | 0.007 | 2.791 | 3 | |
| | RGR12_14 | M5_20483858 | 5 | 20,483,858 | 4.52E-08 | 2.39E-03 | 0.006 | 3.160 | 3 | |
| 5-05 | **PLA10** | **M5_20741662** | **5** | **20,741,662** | **2.19E-13** | **4.65E-08** | **-1.105** | **2.573** | **2** | co-loc |
| | **PLA12** | **M5_20741662** | **5** | **20,741,662** | **2.25E-07** | **9.55E-03** | **-1.822** | **1.302** | **3** | |
| | PLA15 | M5_21091035 | 5 | 21,091,035 | 4.22E-10 | 2.24E-05 | -7.901 | 1.493 | 3 | |
| | PLA18 | M5_21144966 | 5 | 21,144,966 | 7.81E-12 | 8.29E-07 | -31.192 | 1.615 | 3 | |
| 5-08 | **PLA07** | **M5_21202492** | **5** | **21,202,492** | **2.27E-08** | **8.04E-04** | **-0.331** | **2.729** | **3** | co-loc |
| | **PLA09** | **M5_21202492** | **5** | **21,202,492** | **4.60E-08** | **3.25E-03** | **-0.757** | **1.526** | **3** | |
| | PLA18 | M5_22458729 | 5 | 22,458,729 | 3.61E-07 | 7.50E-03 | 12.447 | 3.357 | 3 | |
| | RGR09_11 | M5_24460319 | 5 | 24,460,319 | 1.20E-06 | 2.94E-02 | -0.004 | 1.641 | 4 | |
| | RGR15_17 | M5_26848326 | 5 | 26,848,326 | 3.87E-07 | 7.46E-03 | -0.011 | 1.972 | 4 | |
| 5-06 | **PLA16** | **M5_26861998** | **5** | **26,861,998** | **9.18E-07** | **2.16E-02** | **-7.476** | **1.519** | **4** | co-loc |
| | **PLA18** | **M5_26861998** | **5** | **26,861,998** | **7.86E-08** | **2.38E-03** | **-11.074** | **2.500** | **3** | |

| MTA | trait | SNP | Chrom | position | p.value | q.value | effect | PVE% | PC | category |
|---|---|---|---|---|---|---|---|---|---|---|

bold: significant MTA on consecutive days (dynamic) or co-localisation

PC: number of PCs used for population structure correction
