## Supplementary Table S4 for "Temporal dynamics of QTL effects on vegetative growth in *Arabidopsis thaliana*"

| MTA | trait | stage [DAS] | SNP name | SNP | maf% | minor allele | positive allele | location | aa change | loci in interval |
|---|---|---|---|---|---|---|---|---|---|---|
| 1-01 | PLA | 10-13 | M1_0237637 | A/G | 43.72 | A | A | AT1G07680_e | N60 --> D | AT1G07680 |
| | RGR | 15-17 | M1_02376375 | | | | | | | AT1G07670 |
| | | | | | | | | | | AT1G07690 |
| 1-02a | PLA | 10-18 | M1_2236186 | A/G | 22.77 | A | G | AT1G60750_e | G99 --> R | AT1G60750 |
| 1-02b | PLA | 10-18 | M1_2236224 | A/G | 41.23 | A | G | | | AT1G60730 |
| 1-02c | PLA | 10-18 | M1_2236342 | C/G | 32.20 | G | C | | | AT1G60740 |
| 1-03a | RGR | 8-11 | M1_2238132 | A/G | 19.37 | A | G | AT1G60790_exon2 | | AT1G60790 |
| 1-03b | RGR | 8-11 | M1_2238207 | G/T | 17.02 | T | G | AT1G60790_exon1 | | |
| | | | | | | | | | | AT1G60787 |
| | | | | | | | | | | AT1G60800 |
| 2-01 | PLA | 13-14 | M2_0155077 | A/G | 29.97 | A | G | AT2G04470 | | AT2G04470 |
| | | | | | | | | | | AT2G04460 |
| | | | | | | | | | | AT2G04480 |
| 2-04 | PLA | 16-17 | M2_1307725 | C/T | 21.47 | T | T | AT2G30690_e | V341 --> A | AT2G30690 |
| | | | | | | | | | | AT2G30680 |
| | | | | | | | | | | AT2G30695 |
| 3-01 | RGR | 15-18 | M3_0222139 | A/T | 10.99 | A | A | AT3G07020_e | E64 --> D | AT3G07020 |
| | | | | | | | | | | AT3G07025 |
| | | | | | | | | | | AT3G07030 |
| 3-04a | PLA | 7-10 | M3_0855258 | G/T | 11.52 | T | G | AT3G23740_exon6 | | AT3G23740 |
| 3-04b | RGR | 15-17 | M3_0855314 | A/G | 18.72 | A | A | | | AT3G23730 |
| 3-06 | PLA | 12-16 | M3_1259011 | A/G | 2.36 | A | A | intergenic/5'-UTR AT3G3084 | | AT3G30840 |
| | | | | | | | | | | AT3G30841 |
| 3-08 | PLA | 7-8 | M3_1467281 | C/T | 10.60 | T | T | TE | | AT3G42553 |
| | | | | | | | | | | AT3G06405 |
| 3-07 | PLA | 12-20 | M3_1831508 | A/G | 11.26 | A | G | AT3G49390_intron | | AT3G49390 |
| | | | | | | | | | | AT3G49380 |
| | | | | | | | | | | AT3G49400 |
| 4-01a | RGR | 13-16 | M4_0538929 | C/G | 7.85 | G | C | AT4G08480_intron4 | | AT4G08480 |
| 4-01b | RGR | 13-16 | M4_0538997 | C/T | 6.68 | T | T | AT4G08480_exon1 | | AT4G08470 |

| MTA | trait | stage [DAS] | SNP name | SNP | maf% | minor allele | positive allele | location | aa change | loci in interval |
|---|---|---|---|---|---|---|---|---|---|---|
| | | | | | | | | | | AT4G05545 |
| 4-02 | RGR+PLA | 15-18 | M4_06682595 | C/T | 9.42 | C | T | intergenic | | AT4G10860 |
| | | | | | | | | | | AT4G10865 |
| | | | | | | | | | | AT4G10870 |
| 4-03 | PLA | 14-15 | M4_07928602 | A/T | 48.17 | T | A | intergenic | | none/flanking |
| | | | | | | | | flanking genes | | AT4G13620 |
| | | | | | | | | | | AT4G13615 |
| 4-04 | PLA | 16-17 | M4_08007078 | G/T | 35.60 | T | T | AT4G13810_exon | | AT4G13810 |
| | | | | | | | | | | AT4G13800 |
| | | | | | | | | | | AT4G13820 |
| 4-05 | PLA | 8-11 | M4_10565446 | A/G | 13.48 | G | G | intergenic | | AT4G19350 |
| | | | | | | | | | | AT4G19360 |
| | | | | | | | | | | AT4G19370 |
| | | | | | | | | | | AT4G19380 |
| 4-06 | PLA | 10-16 | M4_16453234 | A/T | 4.19 | T | T | AT4G34410_3' UTR | | AT4G34410 |
| | | | | | | | | AT4G34412_intron5 | | AT4G08985 |
| | | | | | | | | | | AT4G34412 |
| | | | | | | | | | | AT4G03375 |
| | | | | | | | | | | AT4G34415 |
| 4-07 | RGR | 9-15 | M4_17704759 | C/G | 10.73 | C | G | intergenic | | AT4G37680 |
| | | | | | | | | | | AT4G37682 |
| | | | | | | | | | | AT4G37685 |
| | | | | | | | | | | AT4G09605 |
| | | | | | | | | | | At4g37900 |
| 5-01b | PLA | 13-20 | M5_01196098 | C/T | 7.59 | T | C | AT5G04290_exon1 | | AT5G04290 |
| | | | | | | | | | | AT5G04280 |
| | | | | | | | | | | AT5G04275 |
| 5-02 | RGR | 13-17 | M5_08237784 | C/G | 3.53 | G | C | AT5G24260_exon1 | | AT5G24260 |
| | | | | | | | | | | AT5G24240 |
| | | | | | | | | | | AT5G24270 |

| MTA | trait | stage [DAS] | SNP name | SNP | maf% | minor allele | positive allele | location | aa change | loci in interval |
|---|---|---|---|---|---|---|---|---|---|---|
| 5-03 | PLA | 13-15 | M5_13811024 | C/T | 21.07 | C | T | intergenic | | none/flanking |
| | | | | | | | | | | AT5G35605 |
| | | | | | | | | | | AT5G35610 |
| 5-07 | PLA | 7-8 | M5_18554094 | A/C | 43.98 | C | C | intergenic | | AT5G06745 |
| | | | | | | | | | | AT5G45730 |
| | | | | | | | | | | AT5G45740 |
| 1-04a | SA | mature seed | M1_10358650 | A/G | 12.565445 | A | A | TE | | AT1G29650 |
| 1-04b | SW | mature seed | M1_10359910 | G/T | 37.04 | G | G | | | AT1G29640 |
| 1-06b | SA | mature seed | M1_05508567 | C/T | 9.95 | T | T | AT1G16060-e | F2 -> S2 | AT1G16060 |
| 1-06a | RGR | 10-12 | M1_05510455 | C/G | 13.22 | G | G | AT1G16060-3'UTR | | AT1G16040 |
| | | | | | | | | | | AT1G16070 |
| 2-05 | SA/SW | mature seed | M2_4367512 | C/G | 23.82 | G | C | TE | | AT2G11020 |
| | | | | | | | | | | AT2G11010 |
| | | | | | | | | | | AT2G11015 |
| | | | | | | | | | | AT2G11030 |
| 3-09 | SL/SW | mature seed | M3_11700017 | A/G | 39.01 | A | A | TE | | AT3G29792 |
| 4-08 | SL/SW | mature seed | M4_3192692 | A/G | 35.73 | G | G | TE | | AT4G06510 |
| | | | | | | | | | | AT4G06509 |
| 4-09 | SA/SW | mature seed | M4_17448257 | A/G | 37.43 | G | G | intergenic | | AT4G37010 |
| | | | | | | | | | | AT4G37020 |
| | | | | | | | | | | AT4G37022 |

| loci in interval | type | candidate | Description | Functions in | Involved in |
|---|---|---|---|---|---|
| AT1G07680 | protein-coding | | transmembrane protein | molecular_function unknown | biological_process unknown |
| AT1G07670 | protein-coding | yes | endomembrane-type CA-ATPase 4 | calcium-transporting ATPase activity | cation transport, ATP biosynthetic process |
| AT1G07690 | protein-coding | | transmembrane protein | molecular_function unknown | biological_process unknown |
| AT1G60750 | protein-coding | | NAD(P)-linked oxidoreductase superfamily protein | oxidoreductase activity | oxidation reduction |
| AT1G60730 | protein-coding | | NAD(P)-linked oxidoreductase superfamily protein | oxidoreductase activity | oxidation reduction |
| AT1G60740 | protein-coding | | Thioredoxin superfamily protein | oxidoreductase activity | cell redox homeostasis |
| AT1G60790 | protein-coding | yes | trichome birefringence-like protein (DUF828) | O-acetyltransferase activity | cell wall organization or biogenesis |
| AT1G60787 | protein-coding | | cysteine/histidine-rich C1 domain protein | | |
| AT1G60800 | protein-coding | | NSP-interacting kinase 3 | kinase activity | protein amino acid phosphorylation |
| AT2G04470 | TE | | copia-like retrotransposon family | | |
| AT2G04460 | TE | | | | |
| AT2G04480 | protein-coding | | hypothetical protein | | |
| AT2G30690 | protein-coding | | lateral signaling target-like protein (Protein of unknown function, DUF593) | unknown | |
| AT2G30680 | protein-coding | | callose synthase-like protein | 1,3-beta-D-glucan synthase activity | |
| AT2G30695 | protein-coding | | bacterial trigger factor | unknown | protein folding, protein transport |
| AT3G07020 | protein-coding | yes | UDP-Glycosyltransferase superfamily protein | transferase activity, transferring glycosyl groups | lipid glycosylation |
| AT3G07025 | pre_trna | | | | |
| AT3G07030 | protein-coding | yes | Alba DNA/RNA-binding protein | nucleic acid binding | |
| AT3G23740 | protein-coding | | hypothetical protein | | |
| AT3G23730 | protein-coding | yes | xyloglucan endotransglucosylase/hydrolase 16 | hydrolase activity, xyloglucan:xyloglucosyl transferase activity | cellular glucan metabolic process |
| AT3G30840 | protein-coding | | hypothetical protein | | |
| AT3G30841 | protein-coding | | Cofactor-independent phosphoglycerate mutase | catalytic activity, metal ion binding | |
| AT3G42553 | TE | | copia-like retrotransposon family | | |
| AT3G06405 | lncRNA | | | | |
| AT3G49390 | protein-coding | yes | CTC-interacting domain 10 | RNA binding | unknown |
| AT3G49380 | protein-coding | | IQ-domain 15 | | |
| AT3G49400 | protein-coding | | Transducin/WD40 repeat-like superfamily protein | nucleotide binding | unknown |

| loci in interval | type | candidate | Description | Functions in | Involved in |
|---|---|---|---|---|---|
| AT4G08480 | protein-coding | yes | mitogen-activated protein kinase kinase kinase 9 | protein serine/threonine kinase activity, protein kinase activity, kinase activity, ATP binding | protein amino acid phosphorylation |
| AT4G08470 | protein-coding | yes | MAPK/ERK kinase kinase 3 | protein serine/threonine kinase activity, protein kinase activity, kinase activity, ATP binding | protein amino acid phosphorylation |
| AT4G05545 | lncRNA | | | | |
| AT4G10860 | protein-coding | | hypothetical protein | | |
| AT4G10865 | TE | | non-LTR retroelement reverse transcriptase | | |
| AT4G10870 | protein-coding | | hypothetical protein | | |
| none/flanking | | | | | |
| AT4G13620 | protein-coding | yes | Integrase-type DNA-binding superfamily protein | sequence-specific DNA binding transcription factor activity | regulation of transcription, DNA-dependent; ethylene-activated signaling pathway |
| AT4G13615 | protein-coding | | Uncharacterized protein family SERF | | |
| AT4G13810 | protein-coding | yes | receptor like protein 47 | | defense response; signal transduction |
| AT4G13800 | protein-coding | | magnesium transporter NIPA (DUF803) | magnesium ion transmembrane transporter activity | |
| AT4G13820 | protein-coding | yes | Leucine-rich repeat (LRR) family protein | kinase activity | signal transduction |
| AT4G19350 | protein-coding | | embryo defective 3006 | | |
| AT4G19360 | protein-coding | | SCD6 protein-like protein | | |
| AT4G19370 | protein-coding | yes | chitin synthase, putative (DUF1218) | unknown | unknown |
| AT4G19380 | protein-coding | yes | Long-chain fatty alcohol dehydrogenase family protein | electron carrier activity, oxidoreductase activity, acting on CH-OH group of donors, FAD binding | |
| AT4G34410 | protein-coding | yes | redox responsive transcription factor 1 | ethylene-activated signalling pathway; vasculature development, cell division | transcription factor |
| AT4G08985 | lncRNA | yes | Natural antisense transcript overlaps with AT4G34410 | | |
| AT4G34412 | protein-coding | | EKC/KEOPS complex subunit tprkb-like protein | | |
| AT4G03375 | protein-coding | | novel_transcribed_region | | |
| AT4G34415 | tRNA | | | | |

| loci in interval | type | candidate | Description | Functions in | Involved in |
|---|---|---|---|---|---|
| AT4G37680 | protein-coding | yes | heptahelical protein 4 | receptor activity | response to hormone stimulus, response to sucrose stimulus; receptor activity |
| AT4G37682 | protein-coding | | hypothetical protein (DUF1399) | unknown | unknown |
| AT4G37685 | protein-coding | | hypothetical protein (DUF1399) | unknown | unknown |
| AT4G09605 | lncRNA | | | | |
| At4g37900 | protein-coding | yes | hypothetical protein (duplicated DUF1399) | | |
| AT5G04290 | protein-coding | yes | kow domain-containing transcription factor 1 | nucleotide binding; RNA binding, DNA binding; chromatin binding; protein binding | production of siRNA involved in RNA interference; RNA-directed DNA methylation; gene silencing |
| AT5G04280 | protein-coding | yes | RNA-binding (RRM/RBD/RNP motifs) family protein with retrovirus zinc finger-like domain-containing protein | RNA binding, zinc ion binding, nucleotide binding, nucleic acid binding | unknown |
| AT5G04275 | miRNA | yes | microRNA172 that targets several genes containing AP2 domains including AP2 | | flower development, gene silencing by miRNA, meristem determinac |
| AT5G24260 | protein-coding | | prolyl oligopeptidase family protein | serine-type peptidase activity | proteolysis |
| AT5G24240 | protein-coding | yes | phosphatidylinositol 4-kinase gamma-like protein | inositol or phosphatidylinositol kinase activity, phosphotransferase activity, alcohol group as acceptor | |
| AT5G24270 | protein-coding | yes | Calcium-binding EF-hand family protein | | |
| none/flanking | | | | | |
| AT5G35605 | pre_trna | | tRNA-Arg | | |
| AT5G35610 | protein-coding | yes | Paired amphipathic helix (PAH2) superfamily protein | | regulation of transcription |
| AT5G06745 | lncRNA | | | | |
| AT5G45730 | protein-coding | yes | Cysteine/Histidine-rich C1 domain family protein | zinc ion binding | |
| AT5G45740 | protein-coding | | Ubiquitin domain-containing protein | | |
| AT1G29650 | TE | | non-LTR retrotransposon family (LINE) | | |
| AT1G29640 | protein-coding | | senescence regulator (Protein of unknown function, DUF584) | | |
| AT1G16060 | protein-coding | yes | ARIA-interacting double AP2 domain protein | | |
| AT1G16040 | protein-coding | yes | phosphatidylinositol-glycan biosynthesis class F-like protein | unknown | GPI anchor biosynthetic process |
| AT1G16070 | protein-coding | yes | tubby like protein 8 | | |

| loci in interval | type | candidate | Description | Functions in | Involved in |
|---|---|---|---|---|---|
| AT2G11020 | TE | | gypsy-like retrotransposon family | | |
| AT2G11010 | protein-coding | | hypothetical protein | | |
| AT2G11015 | protein-coding | | hypothetical protein | | |
| AT2G11030 | TE | | pseudogene, hypothetical protein | | |
| AT3G29792 | TE | | copia-like retrotransposon family | | |
| AT4G06510 | TE | | gypsy-like retrotransposon family | | |
| AT4G06509 | TE | | gypsy-like retrotransposon family | | |
| AT4G37010 | protein-coding | yes | centrin 2 | calcium ion binding | DNA repair |
| AT4G37020 | protein-coding | yes | initiation factor 4A-like protein | | |
| AT4G37022 | protein-coding | yes | hypothetical protein | | |
