## Supplementary Table S5 for "Temporal dynamics of QTL effects on vegetative growth in *Arabidopsis thaliana*"

| MTA | trait | stage | candidate gene | type | SNP impact | type of polymorphism | gene description | symbol | annotation/literature |
|---|---|---|---|---|---|---|---|---|---|
| 1-01 | PLA | 10-13 | AT1G07670 | protein-coding | moderate | 6 missense SNPs cds | endomembrane-type CA-ATPase 4 | ECA4 | calcium ion transport, ATP biosynthetic process |
| 1-03 | RGR | 8-11 | AT1G60790 | protein-coding | moderate | 6 missense SNPs cds; 19 SNPs promoter; 19 SNPs 3'-UTR | trichome birefringence-like protein (DUF828) | TBL2 | cell wall organization or biogenesis |
| 1-06 | SA | seed | AT1G16060 | protein-coding | moderate | 34 missense SNPs cds | ARIA-interacting double AP2 domain protein | WRI3 | positive regulator of the ABA response, regulating seedling growth. |
| | RGR | 10-12 | AT1G16040 | protein-coding | | | phosphatidylinositol-glycan biosynthesis class F-like protein | | |
| | | | AT1G16070 | protein-coding | | | tubby like protein 8 | TLP8 | Member of TLP family |
| 3-01 | RGR | 15-18 | AT3G07020 | protein-coding | moderate | 10 missense SNPs cds | UDP-Glycosyltransferase superfamily protein | UGT80A2 | lipid glycosylation; reduced seed size |
| | | | AT3G07030 | protein-coding | high; moderate | 1 STOP-gain, 2 frameshift, 1 STOP-loss; 53 missense | Alba DNA/RNA-binding protein | Alba | nucleic acid binding |
| 3-04b | RGR | 15-17 | AT3G23730 | protein-coding | moderate | 25 missense | xyloglucan endotransglucosylase/hydrolase 16 | XTH16 | cell wall modification |
| 3-07 | PLA | 12-20 | AT3G49390 | protein-coding | moderate | 9 missense | CTC-interacting domain 10 | CID10 | RNA-binding protein RBP37 (PAB2) |
| 4-01a | RGR | 13-16 | AT4G08480 | protein-coding | high; moderate | 1 frameshift, 1 in-frame insertion, 1 STOP gain; 87 missense | mitogen-activated protein kinase kinase kinase 9 | MEKK2/SUMM1/ MAPKKK9 | signalling |
| 4-01b | RGR | 13-16 | AT4G08470 | protein-coding | high; moderate | 3 frameshift, 5 splice site, 9 STOP gain; 73 missense, 1 disruptive_inframe_insertion | MAPK/ERK kinase kinase 3 | MEKK3/MAPKKK10 | signalling |
| 4-03 | PLA | 14-15 | AT4G13620 | protein-coding | high; moderate | 9 frameshift, 84 missense | Integrase-type DNA-binding superfamily protein | ERF062 | transcription regulation |

| MTA | trait | stage | candidate gene | type | SNP impact | type of polymorphism | gene description | symbol | annotation/literature |
|---|---|---|---|---|---|---|---|---|---|
| 4-04 | PLA | 16-17 | AT4G13810 | protein-coding | high; moderate | 3 frameshift, 1 splice site, 7 STOP gain; 237 missense | receptor like protein 47 | RLP47 | defense response; signal transduction |
| | | | AT4G13820 | protein-coding | high; moderate | 5 frameshift; 90 missense | Leucine-rich repeat (LRR) family protein | | signal transduction |
| 4-05 | PLA | 8-11 | AT4G19370 | protein-coding | moderate | 13 missense | chitin synthase, putative (DUF1218) | MWL-2 | altered secondary cell wall lignin content |
| | | | AT4G19380 | protein-coding | moderate | 37 missense | Long-chain fatty alcohol dehydrogenase family protein | FAO4A | involved in cell wall establishment or modification |
| 4-06 | PLA | 10-16 | AT4G34410 | protein-coding | moderate | 1 intron variant, 5 downstream, 1 inframe deletion | redox responsive transcription factor 1 | RRTF1; ERF109 | ethylene-activated signalling pathway |
| | | | AT4G08985 | lncRNA | | | Natural antisense transcript overlaps with AT4G34410 | | |
| 4-07 | RGR | 9-15 | AT4G37680 | protein-coding | | | heptahelical protein 4 | HHP4 | |
| | | | At4g37900 | protein-coding | moderate | 78 missense | hypothetical protein (duplicated DUF1399) | GRDP2 | involved in many aspects of plant growth and development as well as response to salt stress |
| 4-09 | SA/SW | mature se | AT4G37010 | protein-coding | | | centrin 2 | CEN2 | Encodes a member of the Centrin family. Mutants are hypersensitive to UV and prone to UV induced DNA damage. Based on sequence similarity and mutant phenotype CEN2 is thought to be involved in nucelotide excision repair/DNA repair. |
| | | | AT4G37020 | protein-coding | | | initiation factor 4A-like protein | | |
| | | | AT4G37022 | protein-coding | | | hypothetical protein | | |

| MTA | trait | stage | candidate gene | type | SNP impact | type of polymorphism | gene description | symbol | annotation/literature |
|---|---|---|---|---|---|---|---|---|---|
| 5-01b | PLA | 13-20 | AT5G04290 | protein-coding | high; moderate | 1 frameshift; 65 missense | kow domain-containing transcription factor 1 | KTF1 / SPT5L | KTF1 is an adaptor protein that binds scaffold transcripts generated by Pol V and recruits AGO4 and AGO4-bound siRNAs to form an RdDM effector complex. Encodes SPT5-Like, a member of the nuclear SPT5 (Suppressor of Ty insertion 5) RNA polymerase (RNAP) elongation factor family that is characterized by the presence of a carboxy-terminal extension with more than 40 WG/GW motifs. Interacts with AGO4. Required for RNA-directed DNA methylation. |
| 5-01a | metabolite | seed/15 | AT5G04280 | protein-coding | moderate | 6 missense | RNA-binding (RRM/RBD/RNP motifs) family protein with retrovirus zinc finger-like domain-containing protein | RZ-1c | Encodes one of the zinc finger-containing glycine-rich RNA-binding proteins involved in cold tolerance: AT3G26420 (ATRZ-1A), AT1G60650 (AtRZ-1b), AT5G04280 (AtRZ-1c). |
| | | | AT5G04275 | miRNA | | | | MIR172B | Encodes a microRNA that targets several genes containing AP2 domains including AP2. Mature sequence: AGAAUCUUGAUGAUGCUGCAU. Pri-mRNA coordinates for MIR172b (converted to TAIR10 based on PMID19304749): Chr5: 1188916-1187500 (reverse), length: 1417 bp; exon coordinates: exon 1: 1188916 to 1188742, exon 2: 1188623 to 1188583, exon 3: 1188383 to 1188133, exon 4: 1187852 to 1187500; mature miRNA and miRNA* are located on exon 3. |

| MTA | trait | stage | candidate gene | type | SNP impact | type of polymorphism | gene description | symbol | annotation/literature |
|---|---|---|---|---|---|---|---|---|---|
| 5-02 | RGR | 13-17 | AT5G24240 | protein-coding | high; moderate | 10 frameshift; 50 missense, 3 disruptive_inframe_deletion | phosphatidylinositol 4-kinase gamma-like protein | PI4Kγ3 | Encodes PI4Kc3, localizes to the nucleus and has autophosphorylation activity, but no lipid kinase activity. Overexpression mutants display late-flowering phenotype. |
| | | | AT5G24270 | protein-coding | moderate | 4 missense | Calcium-binding EF-hand family protein | SOS3; CBL4 | Encodes a calcium sensor that is essential for K+ nutrition, K+/Na+ selectivity, and salt tolerance. The protein is similar to calcineurin B. Lines carrying recessive mutations are hypersensitive to Na+ and Li+ stresses and is unable to grow in low K+. The growth defect is rescued by extracellular calcium. |
| 5-03 | PLA | 13-15 | AT5G35610 | protein-coding | high; moderate | 2 frame shift, 3 splice site, 5 STOP gain, 2 STOP loss; 47 missense | Paired amphipathic helix (PAH2) superfamily protein | | |
| 5-07 | PLA | 7-8 | AT5G45730 | protein-coding | high; moderate | 7 frameshift, 1 START loss, 2 STOP gain; 94 missense | Cysteine/Histidine-rich C1 domain family protein | | |
