## Supplementary Figures for "Temporal dynamics of QTL effects on vegetative growth in *Arabidopsis thaliana*"

Figure S1: Geographic origin of the 382 analysed Arabidopsis accessions

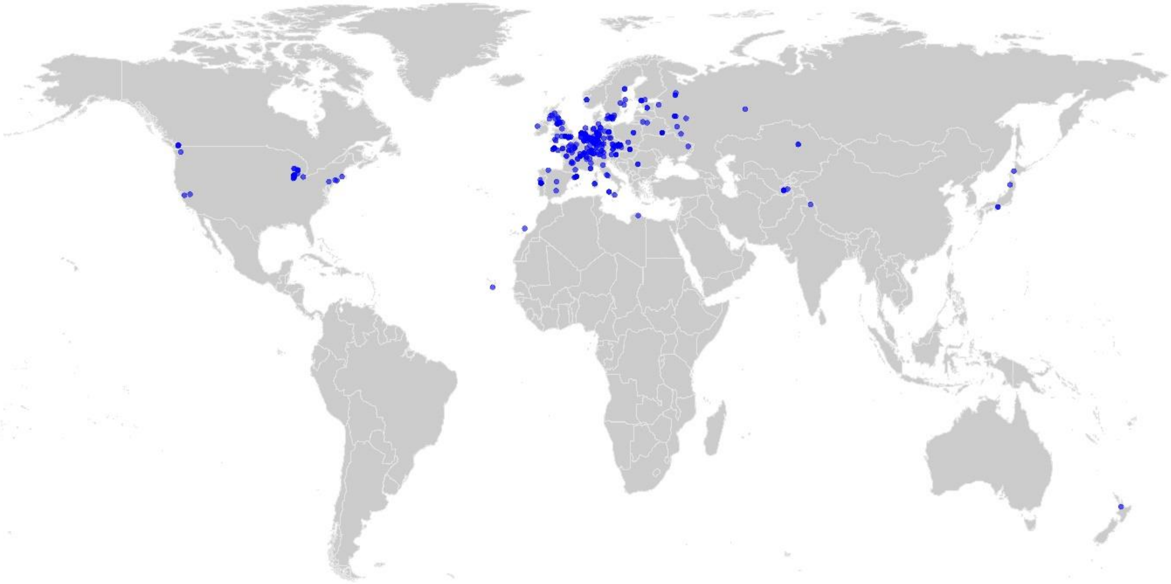

Each blue dot indicates the origin of one accession.

Figure S2: Assessment of population structure

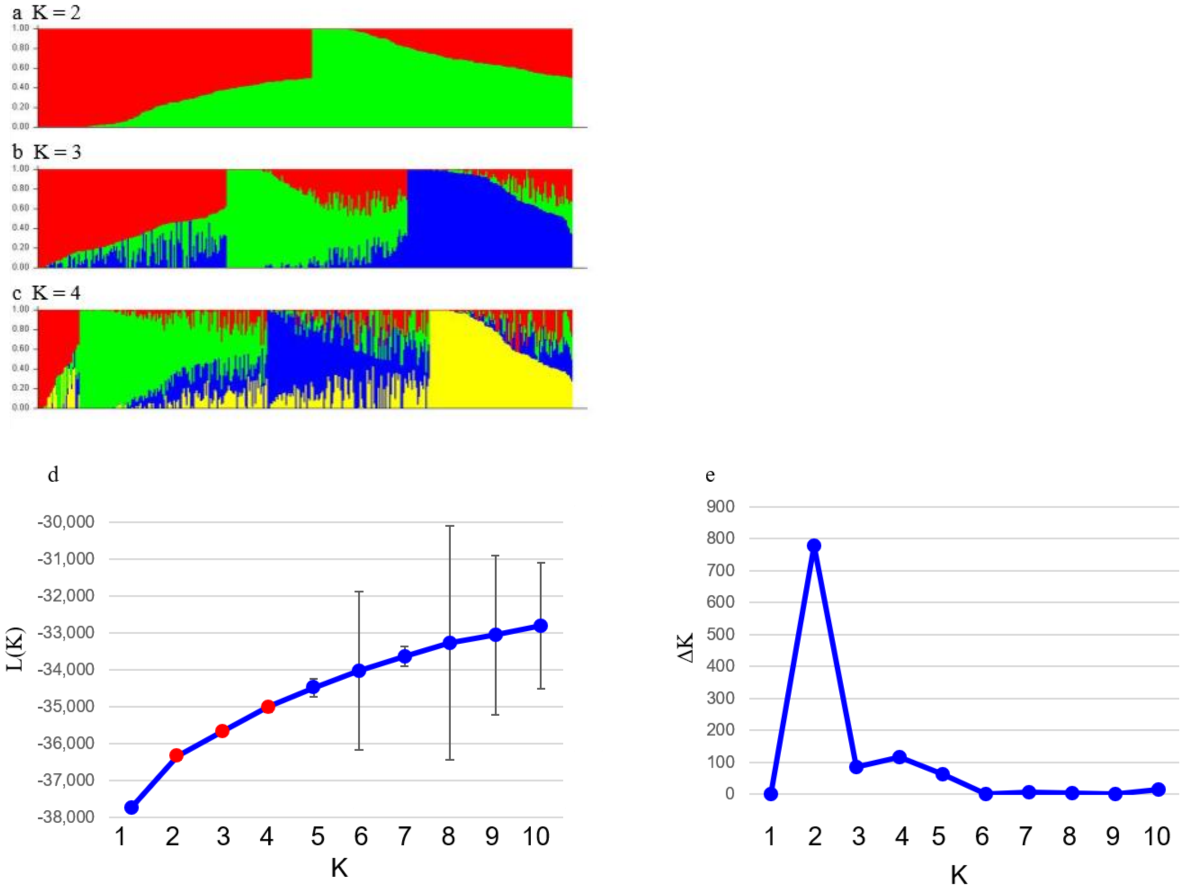


Population structure of the 382 Arabidopsis accession used in the GWAS was assessed using the programme STRUCTURE. Population clustering for *K*= 1 to 10 was performed using the ‘admixture’ model with a burn-in period of 50.000, 50.000 MCMC replications and five iterations per K. The lambda parameter was set to $\lambda$= 0.4619. Subfigures **a-c** shows plots for *K*= 2 to 4. Genotypes were sorted by their ancestry vector (Q). Each genotype is represented by a thin vertical line. Each colour represents a population, and the colour of individual genotypes represents their proportional membership in the different populations. In the subfigures **d-e** statistics used to select K are shown: **d** mean Ln probability L(K) and standard deviation (n = 5); **e** ΔK as the mean difference between successive likelihood values of K (L(K)) averaged over the five runs divided by the standard deviation of L(K)

Figure S3: QQ plots with inclusion of PCs for population structure correction

For each trait the optimum quantile-quantile (QQ) plot after incorporation of the appropriate number of PCs in the model is shown.

The QQ plots compare the expected *p*-values under the null hypothesis (no association) on the x-axis with the observed distribution on the y-axis. The *p*-values are -log10 transformed for easier interpretation. Without bias introduced e.g. by population structure, the dots should follow the red line representin X = Y; the sharp deviation at the upper right corner corresponds to the small number of of true associations aming the 212,142 SNPs tested.

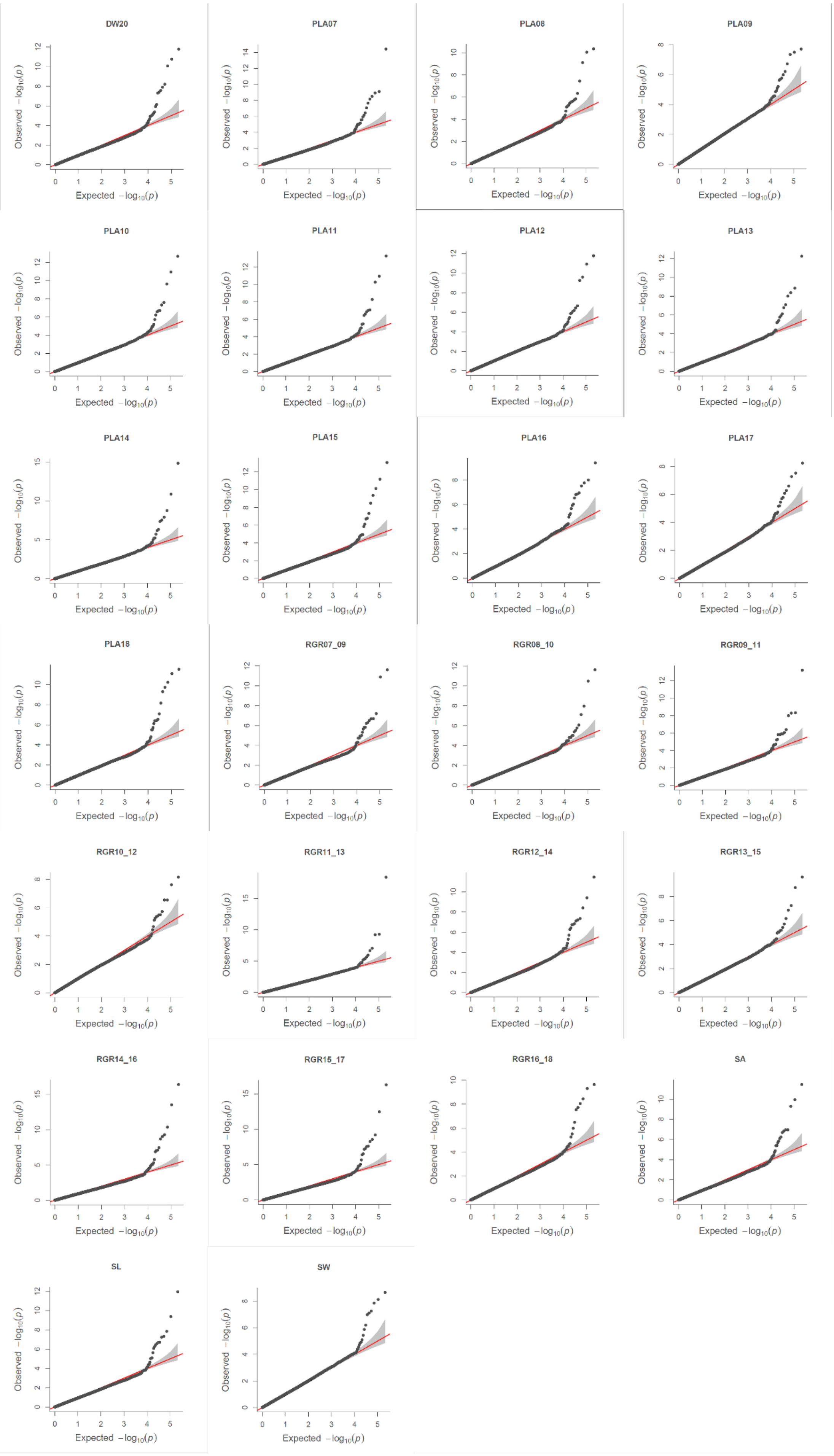
DW20
PLA07
PLA08
PLA09
PLA10
PLA11
PLA12
PLA13
PLA14
PLA15
PLA16
PLA17
PLA18
RGR07_09
RGR08_10
RGR09_11
RGR10_12
RGR11_13
RGR12_14
RGR13_15
RGR14_16
RGR15_17
RGR16_18
SA
SL
SW
Observed $-\log_{10}(p)$
Expected $-\log_{10}(p)$

Figure S4: Duration of dynamic and specific MTA and co-localisation with known QTL

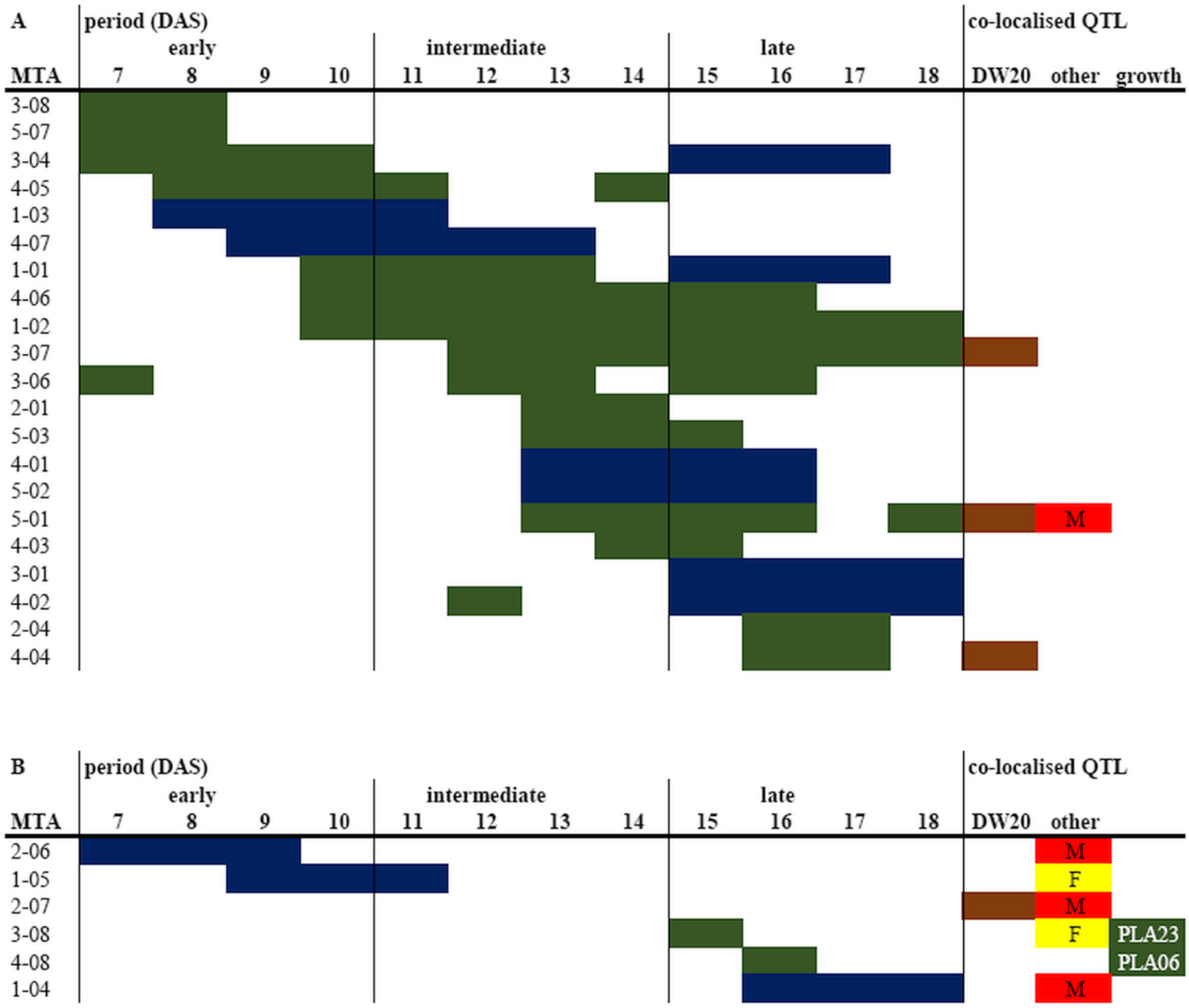


A) The 21 dynamic MTAs significant on at least two consecutive days are represented by horizontal bars across the relevant time period indicated in days after sowing (DAS). green: MTA for projected leaf area; blue: MTA for relative growth rate; brown: MTA for endpoint biomass at 20 DAS; red: metabolic QTL from Knoch et al. (2020) and Lisec et al. (2008).

B) Panel B illustrates co-localisations between single time point MTAs and known QTL. Colour code as above; yellow: flowering; PLA23: QTL 3.68 from Bac-Molenaar et al. (2015); PLA06: QTL from Meyer et al. (2010).

Bac-Molenaar JA, Vreugdenhil D, Granier C, Keurentjes JJB. 2015. Genome-wide association mapping of growth dynamics detects time-specific and general quantitative trait loci. Journal of Experimental Botany 66, 5567-5580.

Knoch D, Abbadi A, Grandke F, Meyer RC, Samans B, Werner CR, Snowdon RJ, Altmann T. 2020. Strong temporal dynamics of QTL action on plant growth progression revealed through high-throughput phenotyping in canola. Plant Biotechnology Journal 18, 68-82.

Lisec J, Meyer RC, Steinfath M, Redestig H, Becher M, Witucka-Wall H, Fiehn O, Törjék O, Selbig J, Altmann T, Willmitzer L. 2008. Identification of metabolic and biomass QTL in Arabidopsis thaliana in a parallel analysis of RIL and IL populations. Plant Journal 53, 960-972.

Meyer RC, Kusterer B, Lisec J, Steinfath M, Becher M, Scharr H, Melchinger AE, Selbig J, Schurr U, Willmitzer L, Altmann T. 2010. QTL analysis of early stage heterosis for biomass in Arabidopsis. Theoretical and Applied Genetics 120, 227-237.
